## Supplemental figures and tables for "Herbivorous insects independently evolved salivary effectors to regulate plant immunity by destabilizing the malectin-LRR RLP NtRLP4"

Xin Wang *et al.*

**This PDF file includes:**

1. Supplementary Figure 1-32
2. Supplementary Table 1-4

### 1. Supplementary Figures

**a**

cleavage site

```
  1: MHKLL AFLVL AGVAL AVPTA TRRCA NYNHY NDWKG NYLVD KEWVV AASTA
 51: SGSVF GWRND YEFCD AYFTY GVPQC GSCTF IRFIP NSRYS GDFGG YTMVQ
101: YAWNS NRQDY DYTEN FIGRA YNGYN APVYR CYTRE PFDRT YFGYK DVMDP
151: DFVPY DAWSN DLTYS AGVLS YCPKN NIVIL VICNN FQYMN YVDNY PIVVV
201: LAQDY KLNAD QKNYI VSELE KNNIN PYGVI LRDNY NCKAR TLYNP SIYL
```

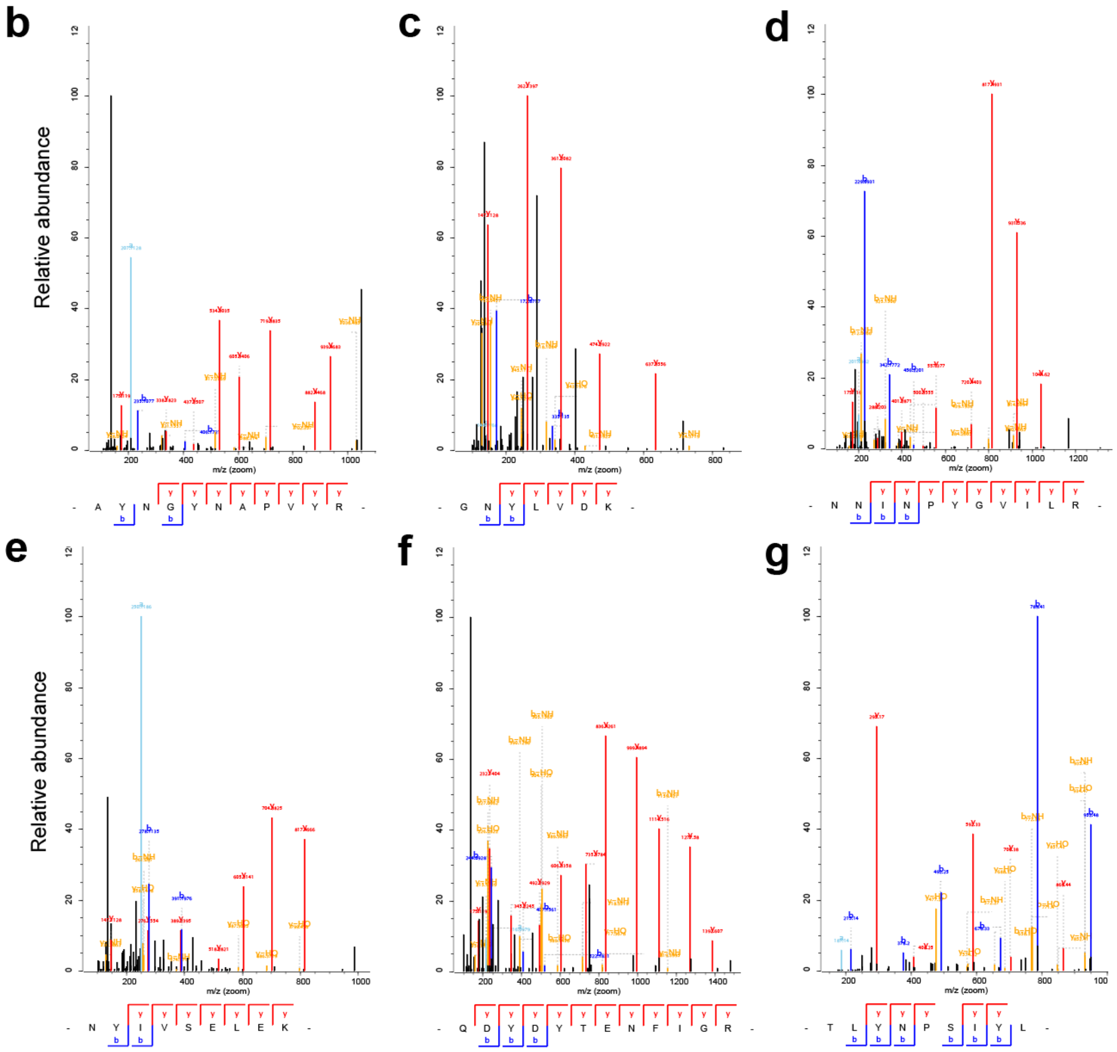


**Fig. S1. Characteristic of BtRDP. (a)** Deduced amino acid sequence of BtRDP. Arrow indicates the signal peptide cleavage site. **(b-g)** Mass spectrums of the identified unique peptides in watery saliva of *Bemisia tabaci*.

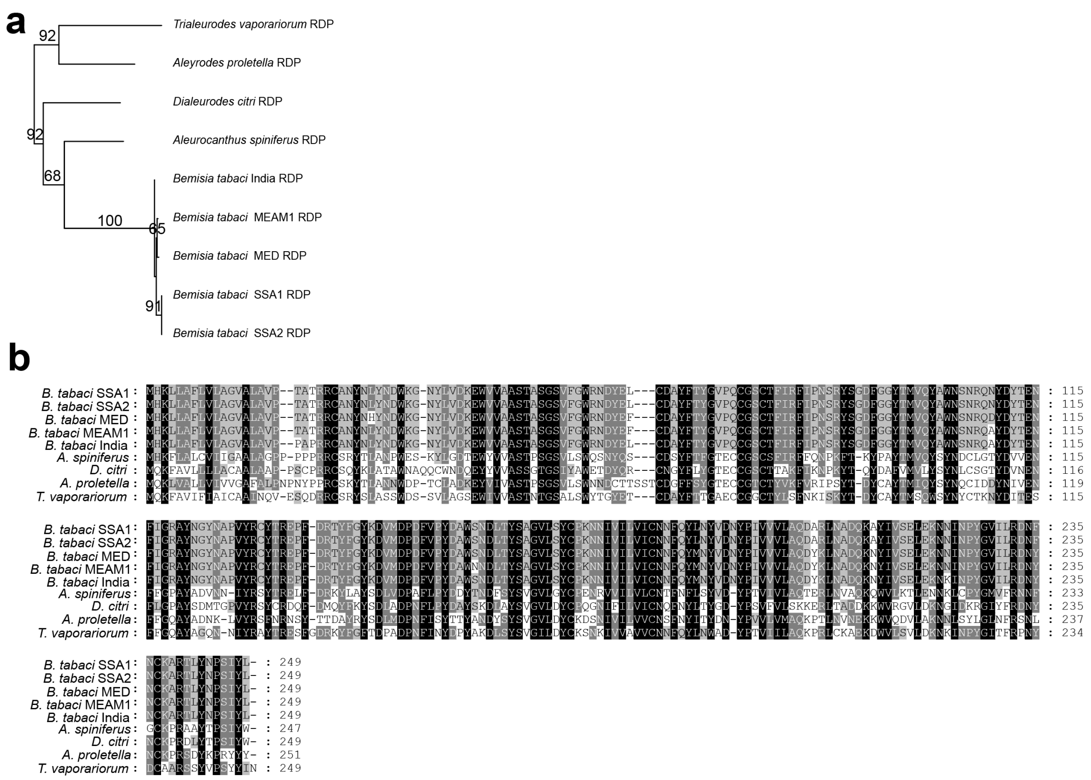


**Fig. S2. Analysis of insect RDP.** (**a**) Phylogenetic tree of RDP sequences from Aleyrodidae species. The trees are constructed by RAxML v0.9.0 using the maximum likelihood method with 1000 bootstrap replicates. Nodes with bootstrap values greater than 50 are displayed. (**b**) Sequence alignments of RDPs. The amino acid sequence of RDPs used for phylogenetic tree construction are aligned using ClusterX software. Black shades indicate the conserved regions.

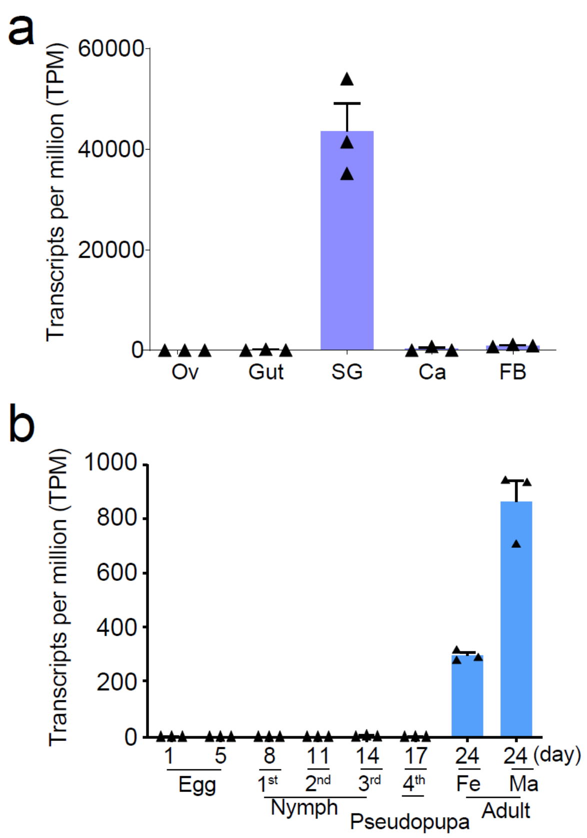


**Fig. S3. Expression patterns of *Bemisia tabaci BtRDP*.** Transcripts per million (TPM) expression values of *BtRDP* in different tissues (**a**) and at different developmental stages (**b**) are determined based on the transcriptomic data. Data are presented as mean values ± SEM (n=3 independent biological replicates).

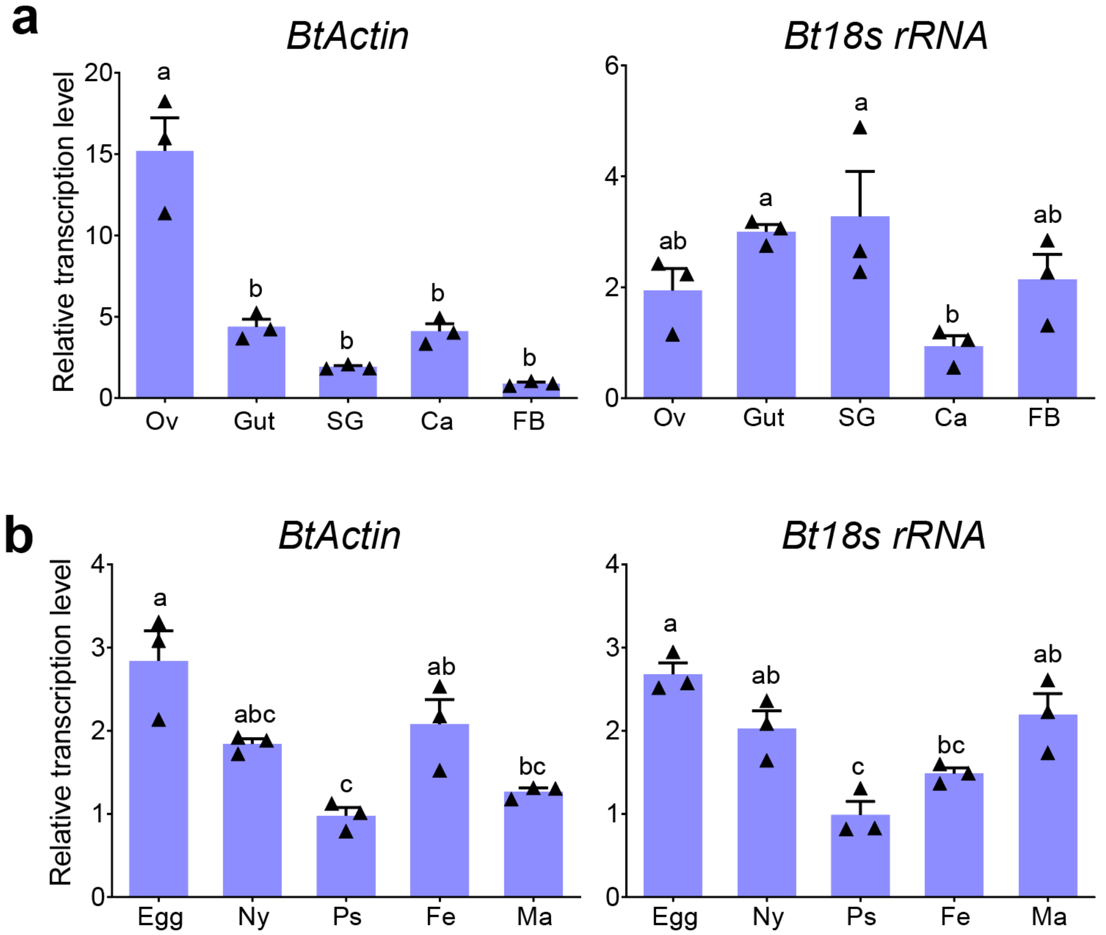


**Fig. S4. Expression patterns of *BtActin* and *Bt18s rRNA*.** The transcript level of each gene in different tissues (**a**) and at different developmental stages (**b**) is quantified by qRT-PCR. *Bemisia tabaci tubulin* is used as an internal control. The relative quantitative method ($2^{-\Delta\Delta Ct}$) is used to evaluate the quantitative variation. Ov, ovary; SG, salivary gland; Ca, carcass; FB, fat body; Ny, nymph; Ps, pseudopupa; Fe, female; Ma, male. Data are presented as mean values ± SEM (n=3 independent biological replicates). Different lowercase letters indicate statistically significant differences at $P < 0.05$ level according to one-way ANOVA test followed by Tukey's multiple comparisons test.

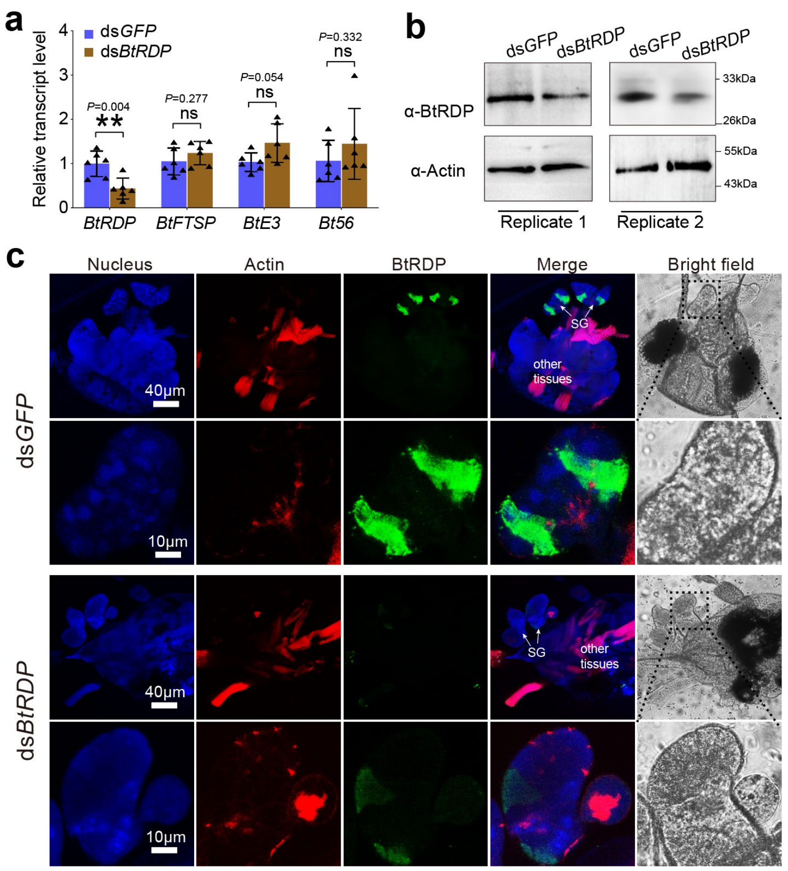


**Fig. S5. BtRDP is efficiently and specifically suppressed by ds*BtRDP*.** (**a**) Relative transcript level of *BtRDP*, *BtFTSP*, *BtE3*, and *Bt56* after dsRNA treatments. *Bemisia tabaci* are treated with ds*GFP* and ds*BtRDP*. The expression pattern of each gene is determined 4 days post treatment using qRT-PCR methods. *B. tabaci tubulin* is used as an internal control. The relative quantitative method ($2^{-\Delta\Delta Ct}$) is used to evaluate the quantitative variation. Data are presented as mean values ± SEM (n= 3 independent biological replicates) (**b**) Protein level of BtRDP after dsRNA treatments. Actin is used as an internal control. Two independent biological replicates are displayed. (**c**) Immunohistochemical staining of BtRDP after dsRNA treatment. The salivary gland (SG) and its nearby tissues are collected from ds*GFP*- and ds*BtRDP*-treated *Bemisia tabaci*. The samples are incubated with anti-BtRDP serum conjugated with Alexa Fluor™ 488 NHS Ester (green) and actin dye phalloidinrhodamine (red), and examined by Leica SP8. The nucleus is stained with DAPI (blue). The lower image in each treatment represents the enlarged images of the boxed area in the upper image. Experiments are repeated twice, with each inspecting more than 10 salivary glands.

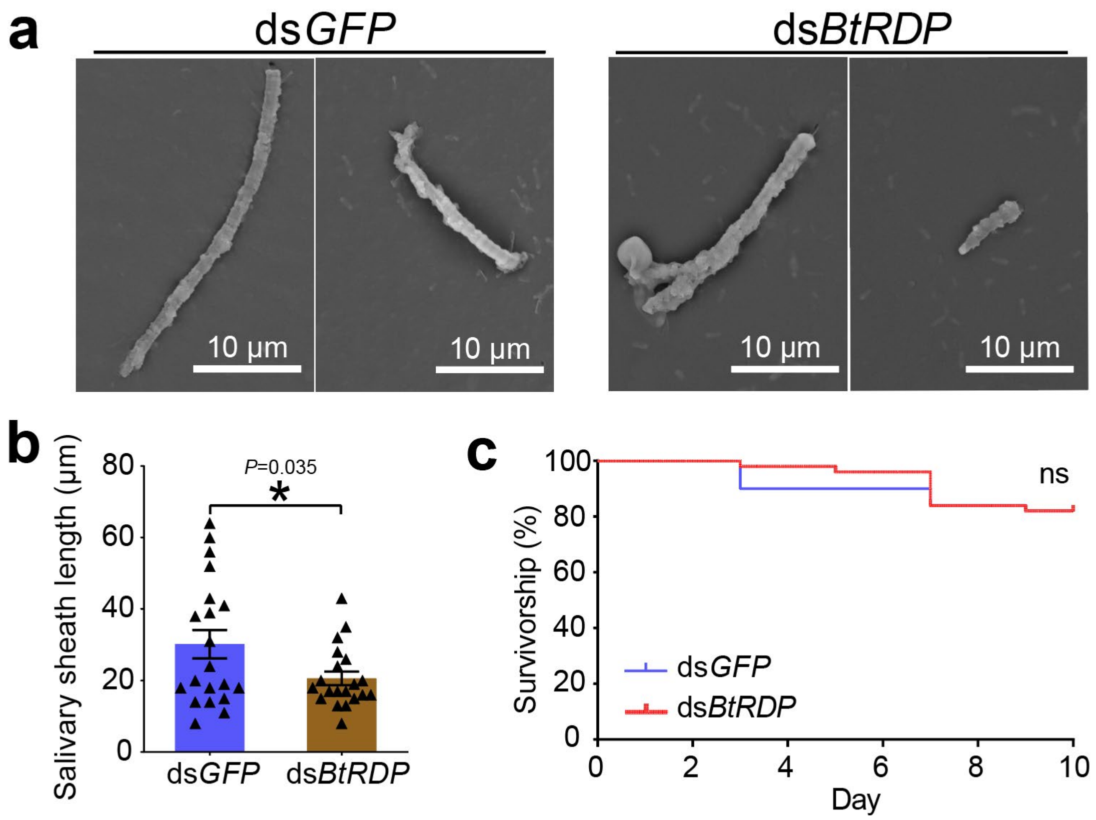


**Fig. S6. Effects of dsRNA treatment on insect survivorship and salivary sheath formation.** (**a**, **b**) Effects of dsRNA treatment on salivary sheath formation. Newly emerged *Bemisia tabaci* adults are injected with ds*GFP* and ds*BtRDP*. The dsRNA-treated *B. tabaci* are fed on artificial diets. The salivary sheaths left on parafilm are inspected by scanning electron microscopy **(a)**, and the length of salivary sheath is measured from the top to base of salivary sheath **(b)**. Data are presented as mean values ± SEM. Twenty salivary sheaths from each treatment are measured. *P*-value is determined by two-tailed unpaired Student's *t* test. *$P < 0.05$. **(c)** Effects of dsRNA treatment on insect survivorship. A group of 20-30 *B. tabaci* are placed in a leaf cage, and their mortality is recorded for ten consecutive days. Three independent biological replications are performed. Differences in survivorship between the two treatments are tested by log-rank test. ns, not significant.

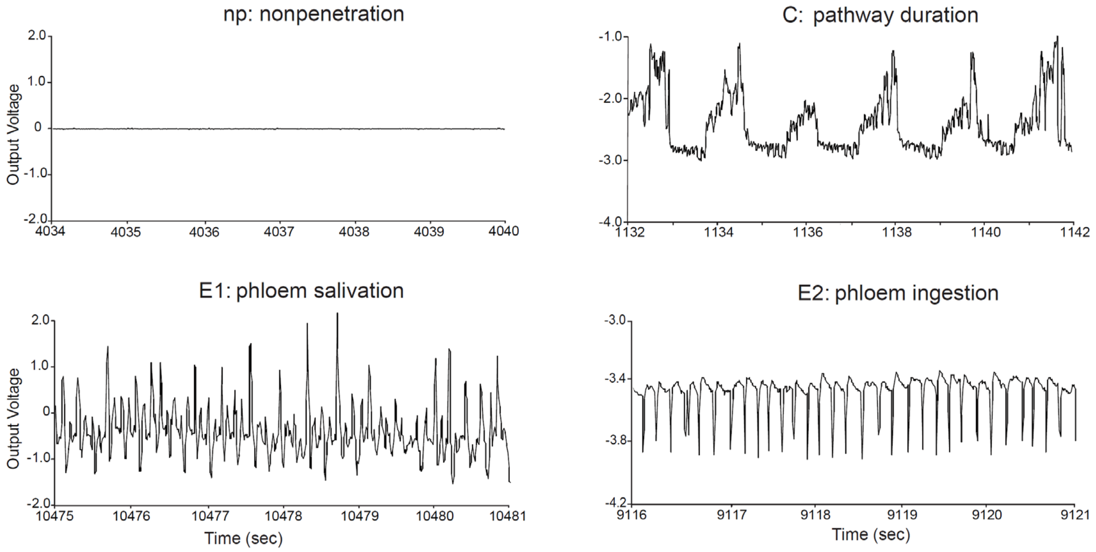


**Fig. S7. Typical EPG waveforms for *Bemisia tabaci* feeding on *Nicotiana tabacum*.** The *B. tabaci* feeding behavior can be classified into nonpenetration (np), pathway duration (C), phloem salivation (E1) and phloem ingestion (E2).

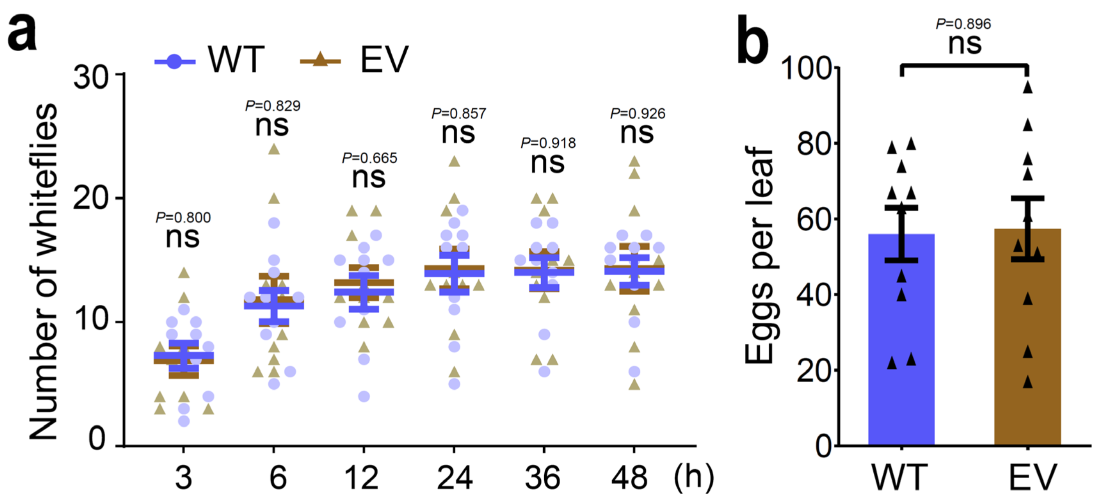


**Fig. S8. Effects of empty vector (EV) transgenic plants on *Bemisia tabaci*.** Two-choice experiment is performed to investigate the attraction of wild type (WT) and EV transgenic plants to *B. tabaci*. A group of 40 female *B. tabaci* are released into a device containing EV and WT leaves. The number of insects settling on each leaf is counted at 3, 6, 12, 24, 36, and 48 h (**a**). After 48 h, the number of eggs on each leaf is counted (**b**). Data are presented as mean values ± SEM. Ten independent biological replicates are performed. *P*-values are determined by two-tailed unpaired Student's *t* test. ns, not significant.

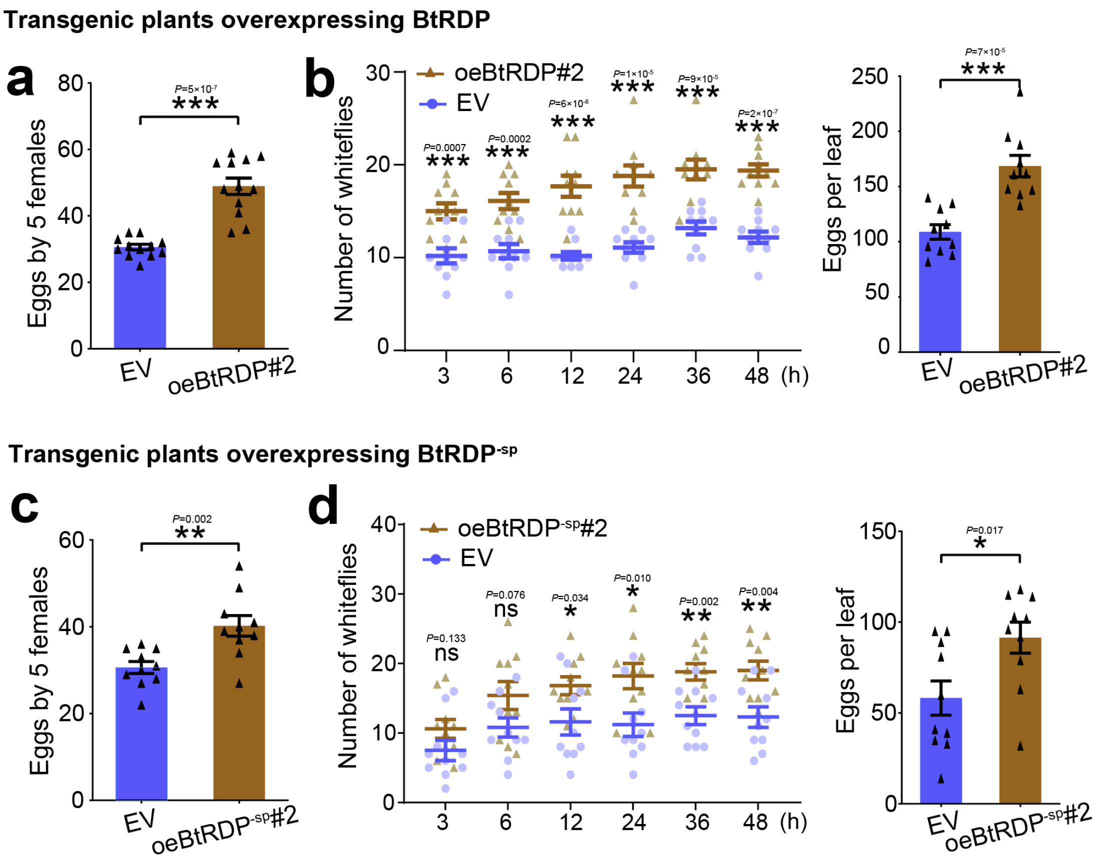


**Fig. S9. Effects of BtRDP and BtRDP$^{-sp}$ overexpression on *Bemisia tabaci*.** Insect performance on oeBtRDP#2 (**a, b**) and oeBtRDP$^{-sp}$#2 (**c, d**) transgenic plants are tested. **(a, c)** Comparison of insect reproduction on empty vector (EV) and oeBtRDP#2/oeBtRDP$^{-sp}$#2 transgenic plants. Five *B. tabaci* individuals are confined to indicated plants for 3 days, and the oviposited eggs are counted. Twelve independent biological replicates are performed. **(b, d)** Attraction of EV and oeBtRDP#2/oeBtRDP$^{-sp}$#2 leaves to *B. tabaci* in a two-choice equipment. A group of 40 female *B. tabaci* are released into a device containing indicated leaves. The number of insects settling on each leaf is counted at 3, 6, 12, 24, 36, and 48 h. After 48 h, the number of eggs on each leaf is counted. Data are presented as mean values ± SEM. Ten independent biological replicates are performed. *P*-values are determined by two-tailed unpaired Student's *t* test. ***$P < 0.001$.

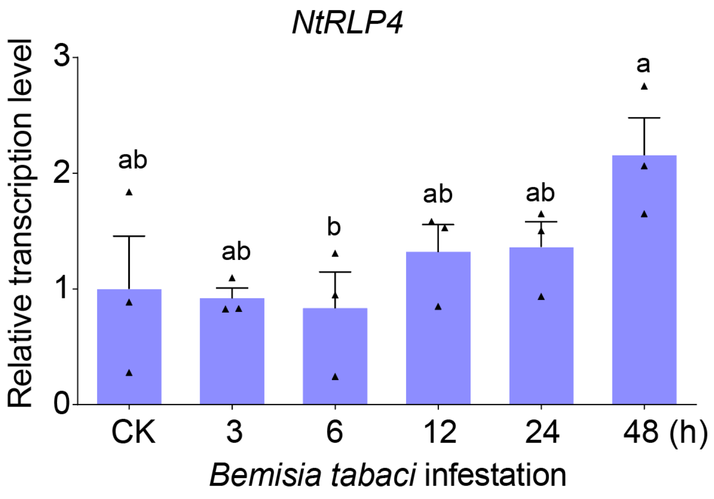


**Fig. S10. Expression patterns of *NtRLP4* in response to *Bemisia tabaci* infestation.** Relative transcript level of *NtRLP4* in response to *B. tabaci* infestation are determined by qRT-PCR. Data are presented as mean values ± SEM. Different lowercase letters indicate statistically significant differences at $P < 0.05$ level according to one-way ANOVA test followed by Tukey's multiple comparisons test. Three independent biological replicates are performed.

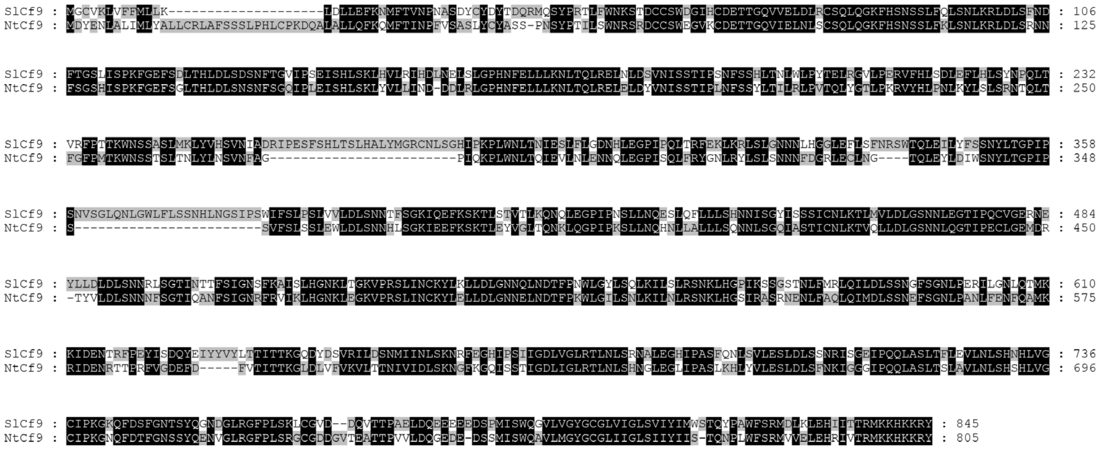

**Fig. S11.** Sequence alignments of SlCf9 and NtCf9. The amino acid sequence of SlCf9 and NtCf9 are aligned using ClusterX software. Black shades indicate the conserved regions.

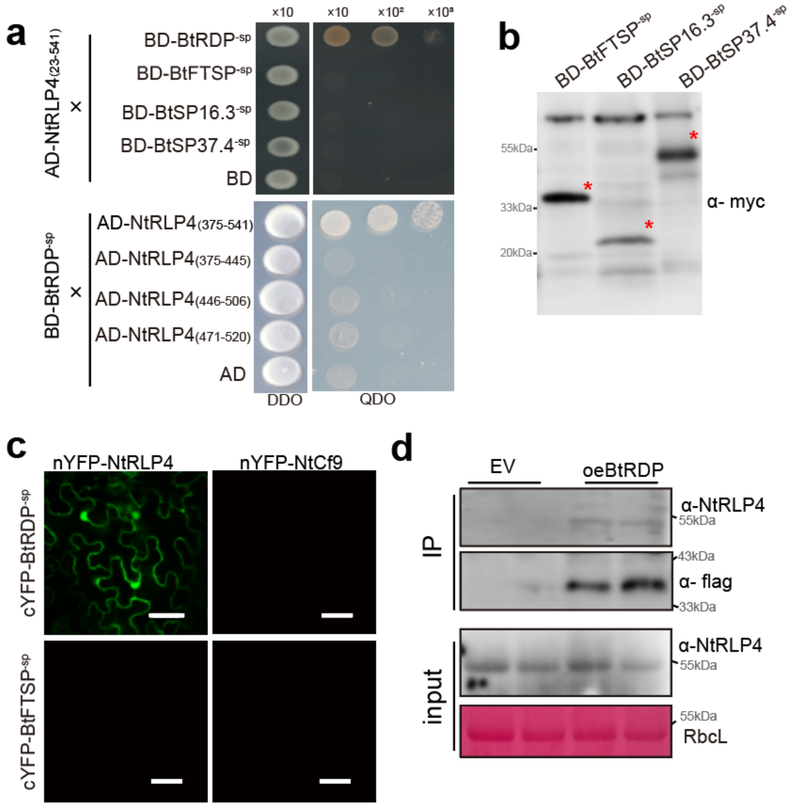


**Fig. S12. BtRDP interacts with NtRLP4.** (**a**) Yeast two hybrid assays showing the interaction between NtRLP4 and different whitefly salivary proteins (BtFTSP$^{-sp}$, BtSP16.3$^{-sp}$ and BtSP37.4$^{-sp}$, without signal peptides), as well as BtRDP$^{-sp}$ and truncated NtRLP4. The different combinations of constructs are transformed into yeast cells, and are grown on the selective medium SD/-Trp/-Leu (DDO), and the interactions are tested with SD/-Trp/-Leu/-His/-Ade (QDO). (**b**) Expression of three control salivary proteins at the protein level. The target bands are indicated by asterisks. **(c)** Bimolecular fluorescence complementation assay showing the specific interaction between BtRDP and NtRLP4. YFP fluorescence is observed when co-expressing the N-terminal nYFP tag fused NtRLP4 and N-terminal cYFP tag fused BtRDP$^{-sp}$. Bar =40 μm. **(d)** Co-immunoprecipitation (Co-IP) assays on oeBtRDP transgenic plants. oeBtRDP transgenic plants that overexpressing BtRDP-flag are incubated with anti-flag beads. Endogenous NtRLP4 can be immunoprecipitated by BtRDP-flag in oeBtRDP transgenic plants, while the flag tag alone in EV plants fails to immunoprecipitate endogenous NtRLP4. Rubisco staining (RbcL) is used to visualize the amount of sample loading.

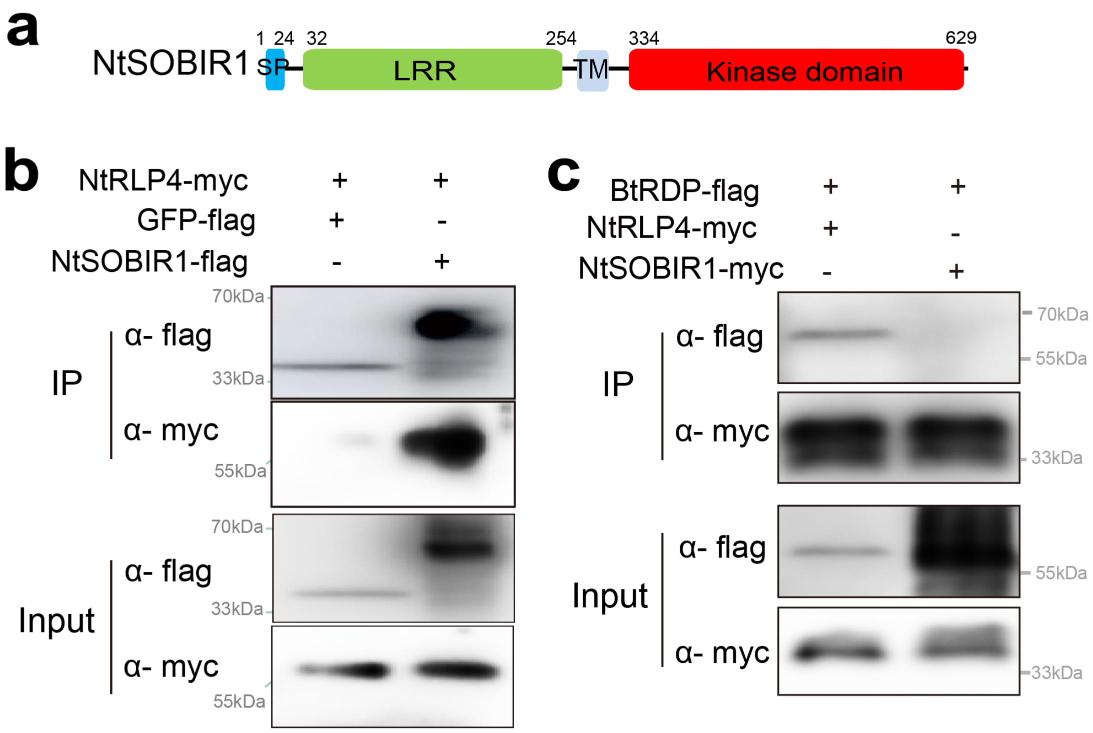


**Fig. S13. NtSOBIR1 interacts with NtRLP4 but not BtRDP. (a)** Domain organization of NtSOBIR1. NtSOBIR1 contains a predicted N-terminal signal peptide (SP), a leucine-rich repeat (LRR) domain, a transmembrane (TM) domain, and a kinase domain. **(b)** Co-immunoprecipitation assay showing the interaction between NtSOBIR1 and NtRLP4. **(c)** Co-immunoprecipitation assay showing the interaction between NtRLP4 and BtRDP, but not NtSOBIR1 and BtRDP. The complete coding region of *NtSOBIR1* and *NtRLP4* are fused with flag and myc tags at the C-terminal ends. Total proteins are extracted from *N. benthamiana* leaves co-expressing NtRLP4-myc with GFP-flag or NtSOBIR1-flag. Precipitation is performed using flag beads. The samples are probed with anti-flag and anti-myc antibodies for immunoblot analysis. Experiments are repeated twice with the similar results.

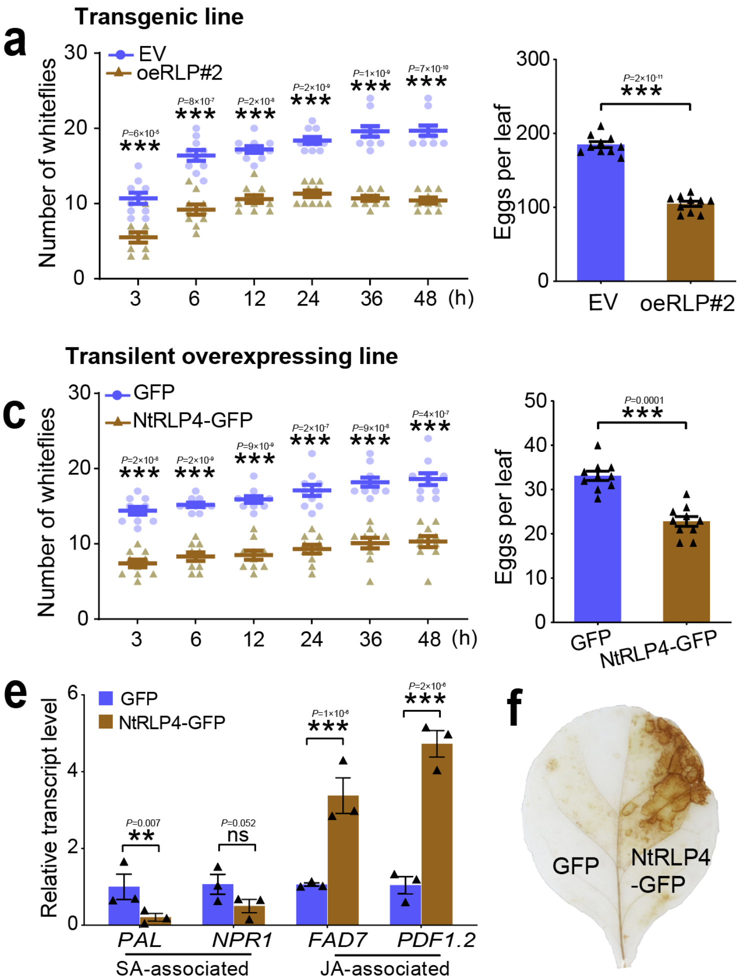


**Fig. S14. Effect of NtRLP4 overexpression on *Bemisia tabaci* performance. (a)** Attraction of empty vector (EV) and oeNtRLP4#2 (oeRLP#2) transgenic *Nicotiana tabacum* to *B. tabaci* in a two-choice equipment. A group of 40 female *B. tabaci* are released into a device containing oeRLP#2 and EV leaves. The number of insects settling on each leaf is counted at 3, 6, 12, 24, 36, and 48 h. After 48 h, the number of eggs on each leaf is counted. **(b)** Comparison of insect reproduction on EV and *oeRLP*#2 transgenic plants. Five *B. tabaci* individuals are confined to indicated plants for 3 days, and the oviposited eggs are counted. **(c)** Attraction of leaves transiently overexpressing GFP and NtRLP4-GFP to *B. tabaci* in a two-choice equipment. **(d)** Comparison of insect reproduction on *N. tabacum* leaves transiently overexpressing GFP and NtRLP4-GFP. **(e)** Relative transcript level of salicylic acid (SA)- and jasmonic acid (JA)-associated genes in *N. tabacum* leaves transiently overexpressing GFP and NtRLP4-GFP. PAL, phenylalanine ammonia lyase; NPR1, nonexpressor of pathogenesis-related protein 1; FAD7, fatty acid desaturase 7; PDF1.2, plant defensin

1.2. **(f)** Transiently overexpressing NtRLP4-GFP induces $H_2O_2$ accumulation. For bioassays in (**a**), (**b**), (**c**), and (**d**), ten independent biological replicates are performed. For qPCR analysis in (**e**), three independent biological replicates are performed. Data are presented as mean values ± SEM. *P*-values are determined by two-tailed unpaired Student's *t* test. ***$P < 0.001$. The experiment in (**f**) is repeated five times with the similar results.

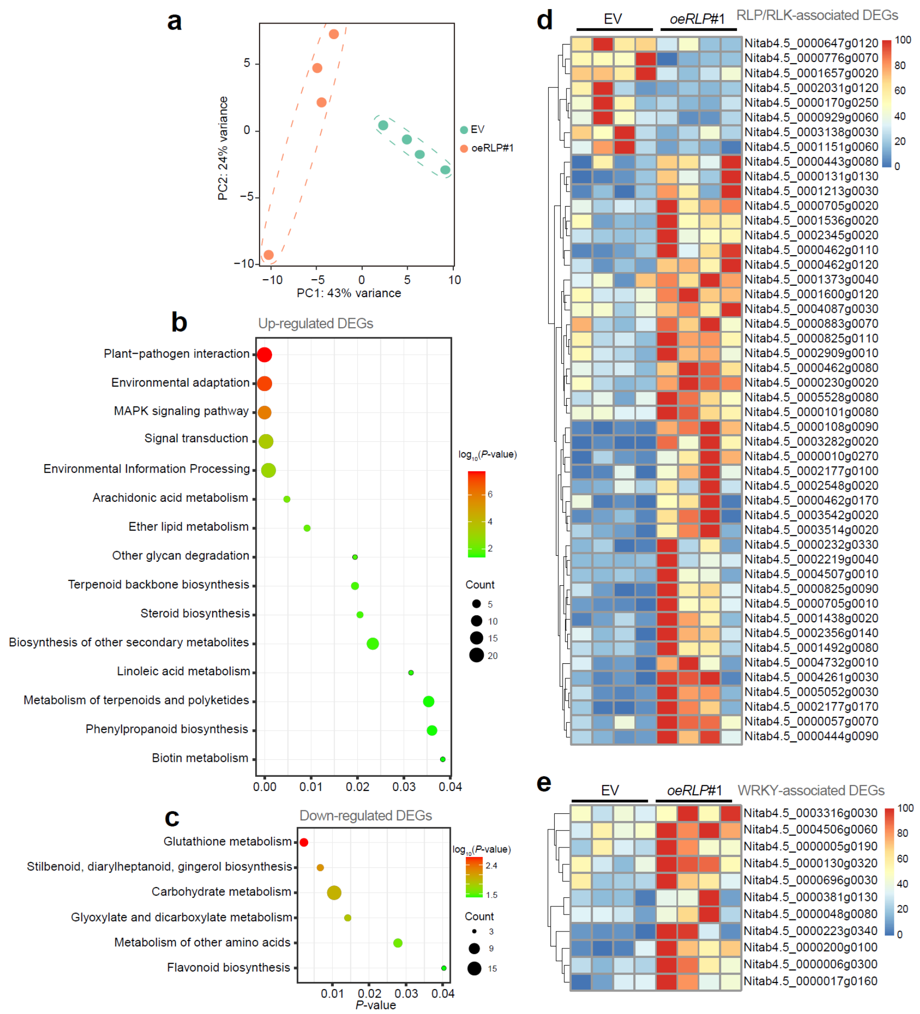


**Fig. S15. Transcriptomic comparison of empty vector (EV) and oeNtRLP4#1 (oeRLP#1) transgenic plants.** (**a**) Principal component analysis (PCA) of gene expression patterns in EV and oeRLP#1 transgenic *Nicotiana tabacum* plants. The first two principal components (PC1 and PC2) based on transcriptomic results are shown. Four independent biological replicates are performed. (**b**, **c**) Kyoto Encyclopedia of Genes and Genomes (KEGG) pathway enrichment analysis of differentially expressed genes (DEGs) that significantly up-regulated (**b**) and down-regulated (**c**). Enriched *P*-values are calculated according to one-sided hypergeometric test using TBtools software. (**d, e**) Expression pattens of DEGs that annotated as RLK/RLP (**d**) or WRKY transcription factor (**e**). Gene expression patterns are illustrated by a heatmap. The max transcripts per million value of each gene is set as 100.

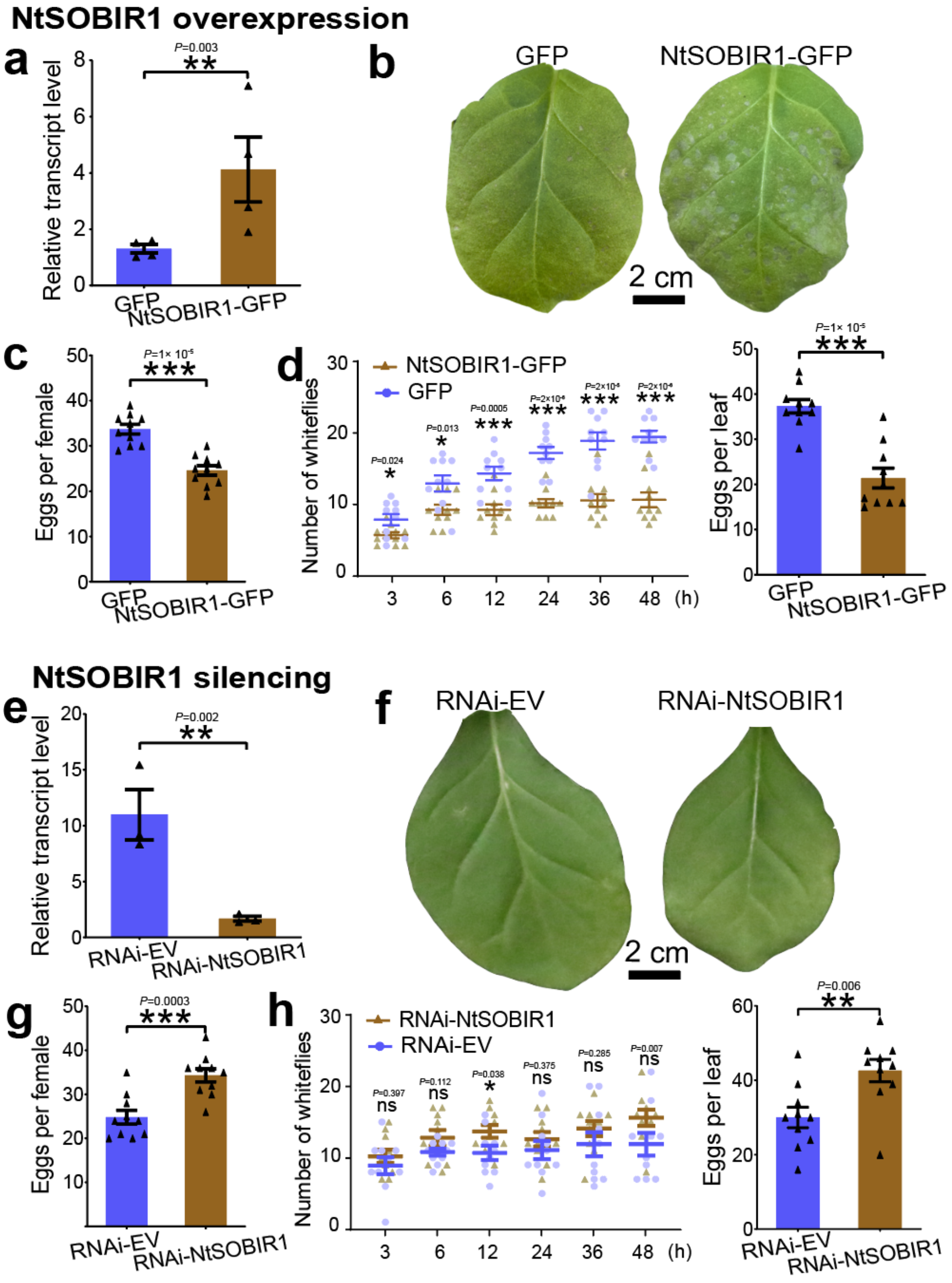


**Fig. S16. Effect of NtSOBIR1 overexpression and silencing on *Bemisia tabaci* performance.** (**a**) Validation of NtSOBIR1 transient overexpression using qRT-PCR (n= 4 biological replicates). (**b**) NtSOBIR1 transient overexpression induces cell death phenotype five days post agro-injection. **(c)** Comparison of insect reproduction on *Nicotiana tabacum* transiently overexpressing GFP and NtSOBIR1-GFP. Five *B. tabaci* individuals are confined to indicated plants for 3 days, and the oviposited eggs are counted. **(d)** Attraction of GFP and NtSOBIR1-GFP leaves to *B. tabaci* in a two-choice equipment. A group of 40 female *B. tabaci* are released into a device containing

GFP and NtSOBIR1-GFP leaves. The number of insects settling on each leaf is counted at 3, 6, 12, 24, 36, and 48 h (left). After 48 h, the number of eggs on each leaf is counted (right). (**e**) Silencing efficiency of hairpin NtSOBIR1 using qRT-PCR (n= 3 independent biological replicates). (**f**) Transiently silencing NtSOBIR1 affect growth of infiltrated leaves. **(g)** Comparison of insect reproduction on *Nicotiana tabacum* leaves infiltrated with hairpin empty vector (EV) and NtSOBIR1. **(h)** Attraction of hairpin EV and NtSOBIR1-treated leaves to *B. tabaci* in a two-choice equipment. The experiments in (**b**) and (**f**) are repeated ten times with the similar results. In (**c**), (**d**), (**g**), and (**h**), ten independent biological replicates are performed. *P*-values are determined by two-tailed unpaired Student's *t* test. ****P* < 0.001; ***P* < 0.01.

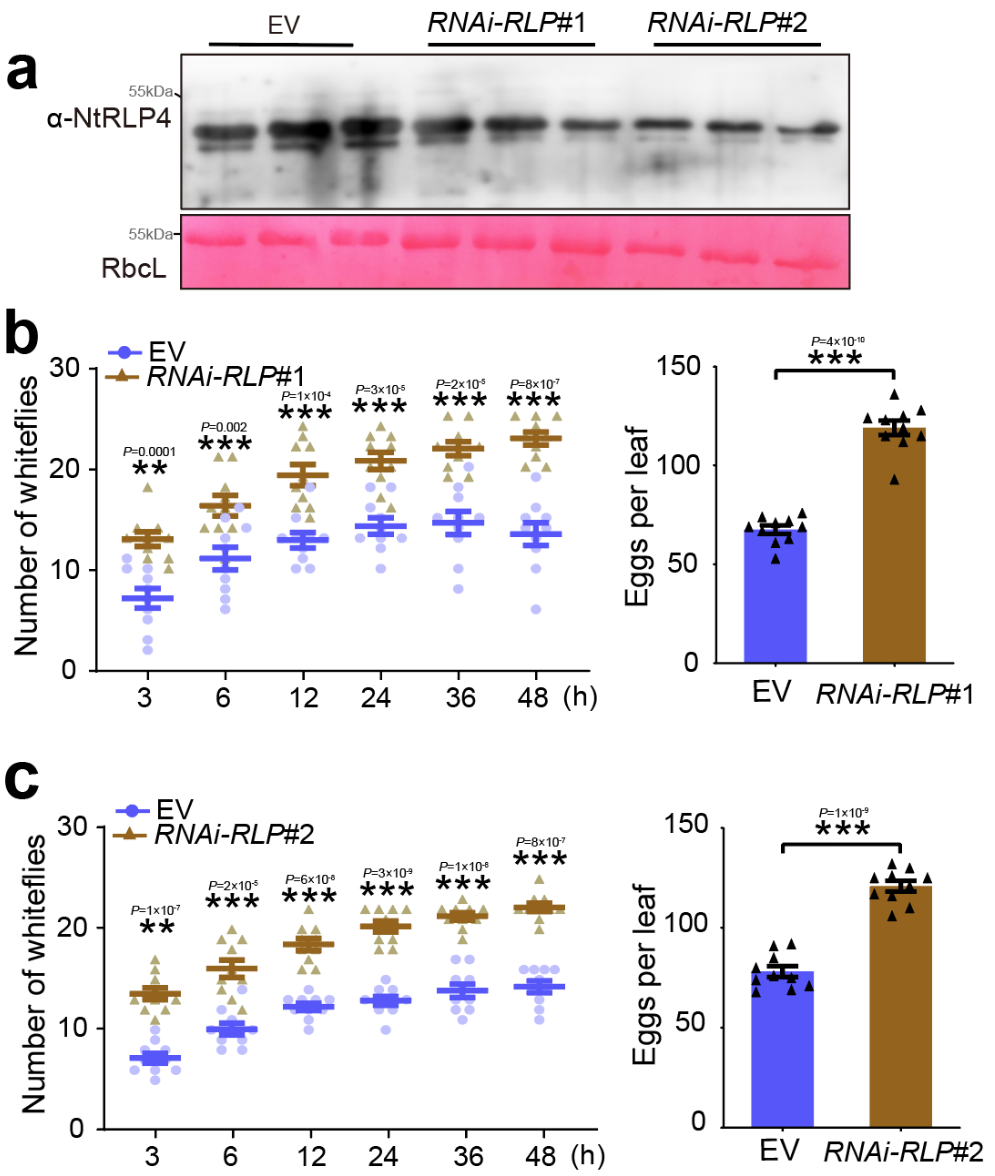


**Fig. S17. Effects of *RLP* silencing on *Bemisia tabaci* performance**. (**a**) Detection of NtRLP4 protein in empty vector (EV) and *NtRLP4*-silenced (*RNAi-RLP*) transgenic *Nicotiana tabacum*. Rubisco staining (RbcL) is conducted to visualize the amount of sample loading. **(b**, **c)** Attraction of EV and *RNAi-RLP* plants to *B. tabaci* in a two-choice equipment. Two NtRLP4-silenced plants (RNAi-RLP#1 in **b**; RNAi-RLP#2 in **c**) are assayed. A group of 40 female *B. tabaci* are released into a device containing two leaves. The number of insects settling on each leaf is counted at 3, 6, 12, 24, 36, and 48 h. After 48 h, the number of eggs on each leaf is counted. Ten independent biological replicates are performed. Data are presented as mean values ± SEM. *P*-values are determined by two-tailed unpaired Student's *t* test. ***$P < 0.001$.

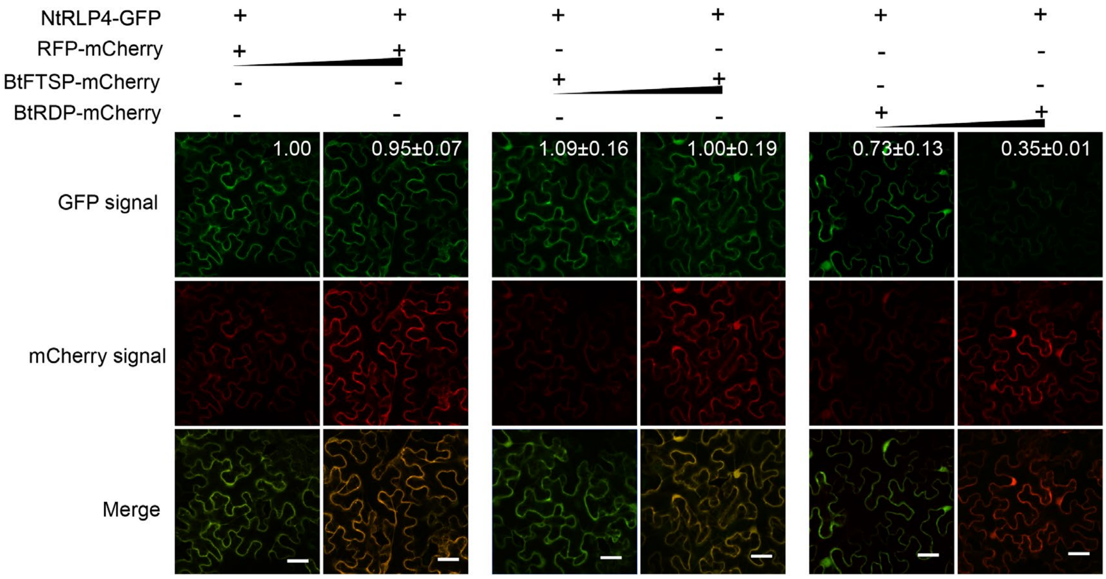


**Fig. S18. Influence of BtRDP on NtRLP4 accumulation by fluorescent analysis.** The NtRLP4-GFP is transiently co-expressed with red fluorescent protein (RFP)-mCherry, BtFTSP-mCherry, or BtRDP-mCherry via agroinfiltration. The samples are imaged by confocal microscopy at 48 h post injection. RFP-mCherry and BtFTSP-mCherry are used as negative controls. Three independent biological replicates are performed, and three representative images are taken in each biological replicate. Fluorescence intensity is measured using ImageJ. The intensity values from three biological replicates are calculated and the mean value in the upper left image is set at 1.0. Representative fluorescence images are displayed. The small triangle indicates the different concentrations ($OD_{600}$ = 0.1 and 1.0) of *Agrobacterium*. Bar= 40 µm.

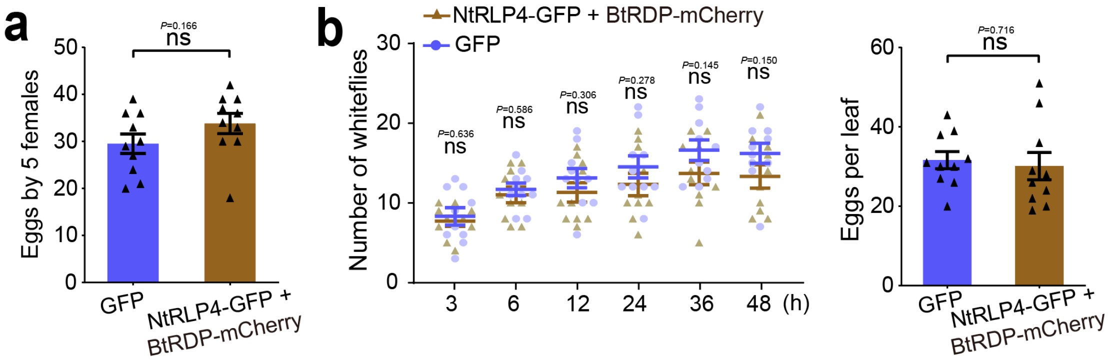


**Fig. S19. Effects of BtRDP on suppressing NtRLP4-associated plant defenses. (a)** Comparison of insect reproduction on *Nicotiana tabacum* plants transiently overexpressing GFP along and BtRDP-mCherry/NtRLP4-GFP. Five *Bemisia tabaci* individuals are confined to indicated plants for 3 days, and the oviposited eggs are counted. **(b)** Attraction of GFP- and BtRDP-mCherry/NtRLP4-GFP-expressed leaves to *B. tabaci* in a two-choice equipment. A group of 40 female *B. tabaci* are released into a device containing two leaves. The number of insects settling on each leaf is counted at 3, 6, 12, 24, 36, and 48 h. After 48 h, the number of eggs on each leaf is counted. Data are presented as mean values ± SEM. Ten independent biological replicates are performed. *P*-values are determined by two-tailed unpaired Student's *t* test. *$P < 0.05$; ns, not significant.

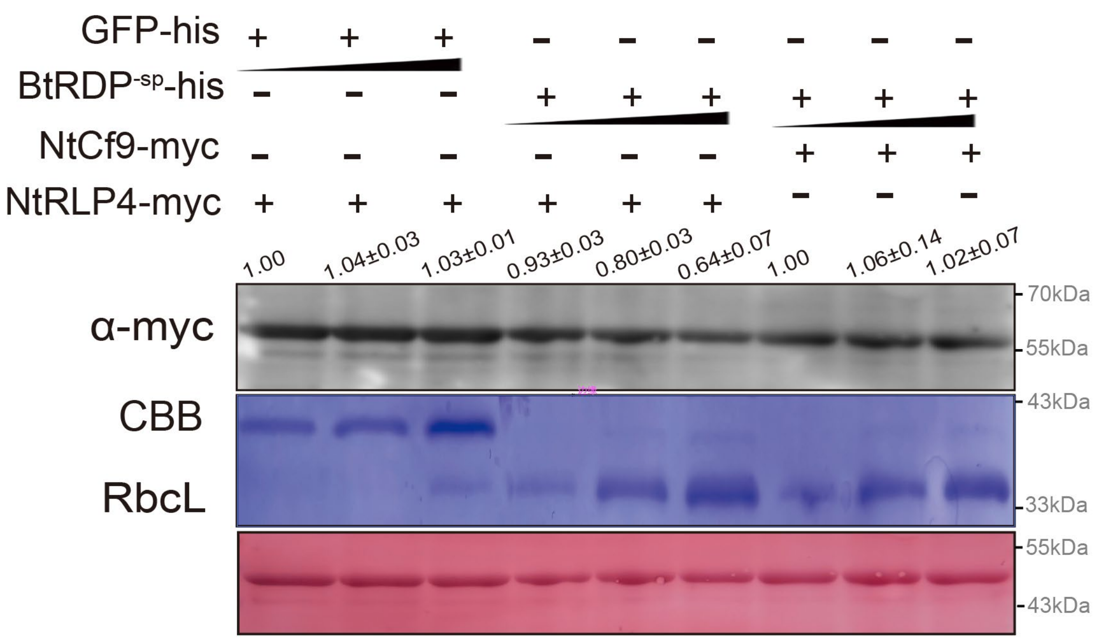


**Fig. S20. Degradation of NtRLP4 by purified BtRDP$^{-sp}$ in *Nicotiana benthamiana* leaves.** Recombinant BtRDP$^{-sp}$-his and GFP-his proteins are expressed in *Escherichia coli*, and purified by Ni-NTA. *N. benthamiana* plants overexpressing NtRLP4-myc or NtCf9-myc are then infiltrated with different concentrations of purified BtRDP$^{-sp}$-his and GFP-his. The samples are probed with anti-myc antibodies for immunoblot analysis. Rubisco staining (RbcL) is conducted to visualize the amount of sample loading. Coomassie brilliant blue (CBB) staining is conducted to visualize the amount of recombinant BtRDP$^{-sp}$-his and GFP-his proteins. Experiments are repeated three times with the similar results. Band density is measured using ImageJ. The density values from three biological replicates are calculated and the mean value in the first lane is set at 1.0. Data are presented as mean values ± SEM.

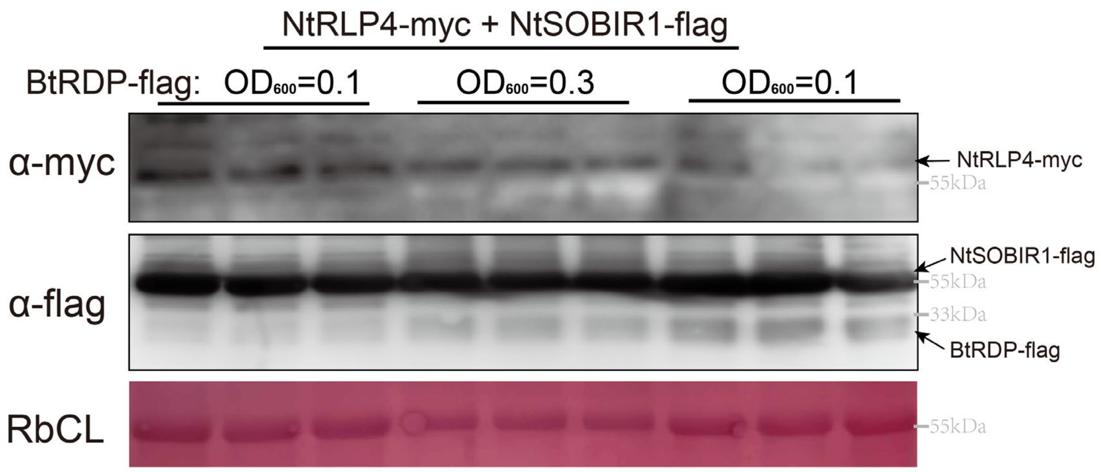


**Fig. S21. Effect of BtRDP on NtRLP4 and NtSOBIR1.** NtRLP4-myc and NtSOBIR1-flag were transiently co-expressed with different concentration of BtRDP-flag in *Nicotiana benthamiana* plants through *Agrobacterium* infiltration. The samples are probed with anti-flag and anti-myc antibodies for immunoblot analysis. Rubisco staining (RbcL) is conducted to visualize the amount of sample loading. The amount of BtRDP were controlled by infiltrating different concentrations of *Agrobacterium* ($OD_{600}$ = 0.1, 0.3, and 1.0).

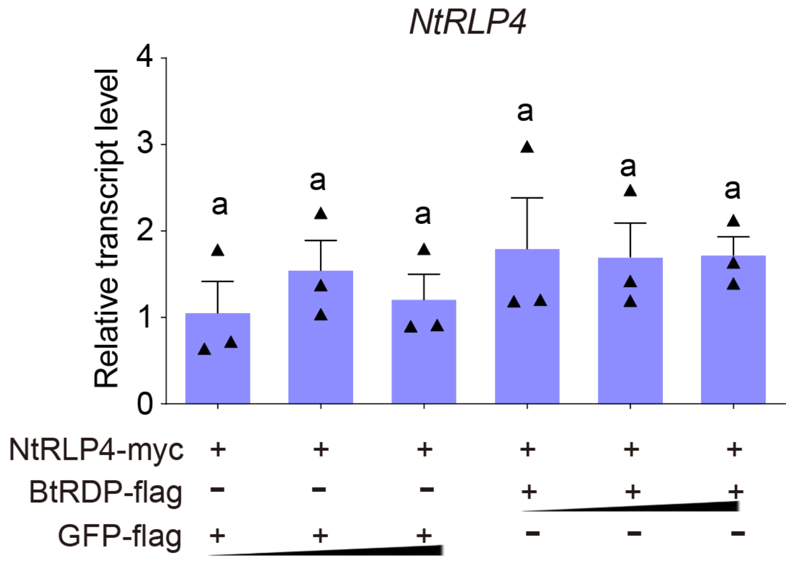


**Fig. S22. Effect of BtRDP on the transcript level of *NtRLP4*.** NtRLP4-myc is agroinfiltrated together with different concentration of BtRDP-flag or GFP-flag into *Nicotiana benthamiana* leaves. The transcript level of *NtRLP4* is determined by qRT-PCR. Data are presented as mean values ± SEM. Three independent biological replicates are performed. The same lowercase letters "a" indicate no statistically significant differences at $P < 0.05$ according to the one-way ANOVA test followed by Tukey's multiple comparisons test.

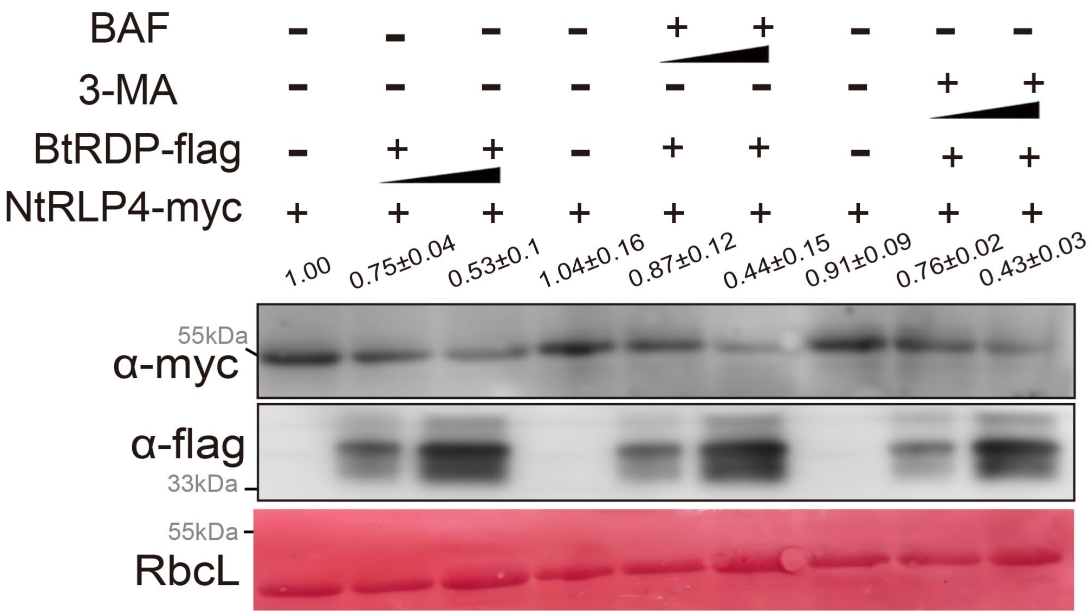


**Fig. S23. Effects of autophagy inhibitor on NtRLP4 accumulation.** NtRLP4-myc and BtRDP-flag were transiently co-expressed in *Nicotiana benthamiana* plants through *Agrobacterium* infiltration. Co-infiltrated leaves are treated with autophagy inhibitor BAF and 3-MA at 24 h post injection. The samples are probed with anti-flag and anti-myc antibodies for immunoblot analysis. Rubisco staining (RbcL) is conducted to visualize the amount of sample loading. The small triangle indicates the different concentrations of *Agrobacterium* ($OD_{600}$ = 0.1 and 1.0). Experiments are repeated three times with the similar results. Band density is measured using ImageJ. The density values from three biological replicates are calculated and the mean value in the first lane is set at 1.0. Data are presented as mean values ± SEM. *P*-values are determined by two-tailed unpaired Student's *t* test. ***$P < 0.001$; **$P < 0.01$; *$P < 0.05$; ns, not significant.

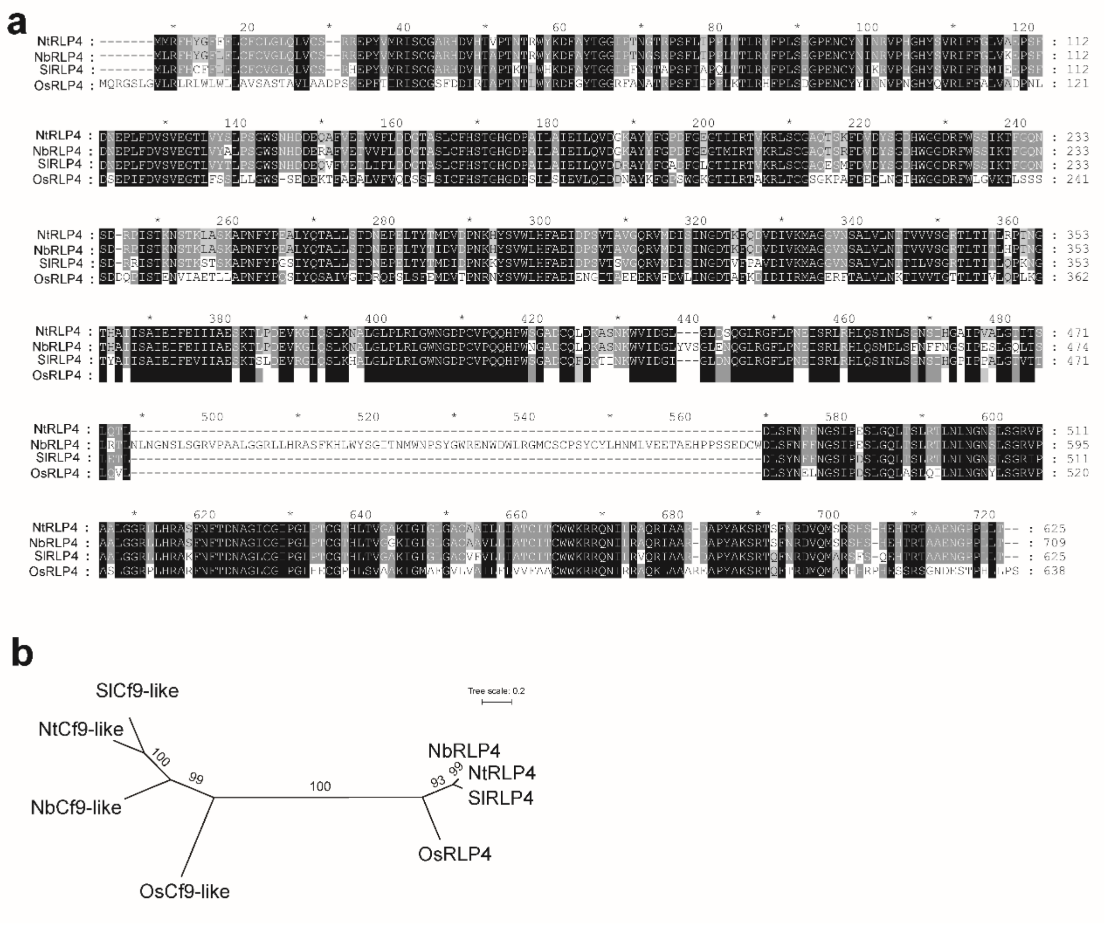


**Fig. S24. Sequence alignment and phylogenetic tree of RLP4 homologs.** (**a**) The amino acid sequence of RLP4s from *Nicotiana tabacum*, *N. benthamiana*, *Solanum lycopersicum*, and *Oryza sativa* are aligned using ClusterX software. Black shades indicate the conserved regions. (**b**) Phylogenetic analysis of RLP4 and Cf9-associated genes used in this study. The unrooted phylogenetic trees are constructed by RAxML v0.9.0 using the maximum likelihood method with 1000 bootstrap replicates.

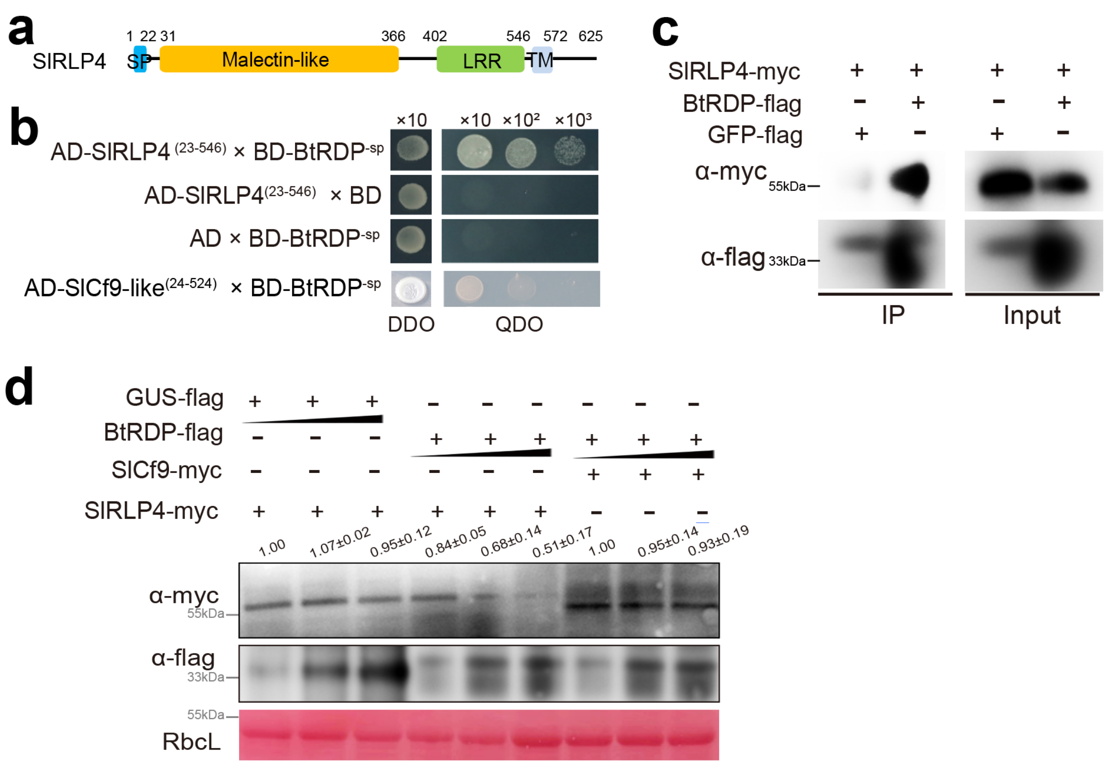


**Fig. S25. Influence of BtRDP on RLP4 homology in *Solanum lycopersicum*. (a)** Domain organization of *S. lycopersicum* RLP4 (SlRLP4). SlRLP4 contains a predicted N-terminal signal peptide (SP), a malectin-like domain, a LRR domain, and a transmembrane (TM) domain. (**b, c**) Yeast two-hybrid and co-immunoprecipitation (Co-IP) assays showing the interaction between BtRDP and SlRLP4. In (**b**), BtRDP is expressed without a signal peptide (BtRDP$^{-sp}$), while SlRLP4 and SlCf9 are expressed without a signal peptide and transmembrane domain (SlRLP4$^{(23\text{-}546)}$ and SlCf9$^{(24\text{-}524)}$). **(d)** Effect of BtRDP on the accumulation of SlRLP4. SlRLP4-myc and SlCf9-myc are agro-injected together with different concentration of BtRDP-flag or GFP-flag. In (**c, d**), the complete coding region of *BtRDP*, *SlRLP4*, *SfCf9* are fused with flag or myc tags at C-terminal ends, respectively. The small triangle indicates the different concentrations ($OD_{600}$ = 0.05, 0.3, and 1.0) of *Agrobacterium*. Rubisco staining (RbcL) is conducted to visualize the amount of sample loading. Experiments are repeated three times with the similar results. Band density is measured using ImageJ. The density values from three biological replicates are calculated. For SlRLP4-myc bands (lane 1-6), the lane 1 is set at 1.0. For SlCf9-myc bands (lane 7-9), the lane 7 is set at 1.0.

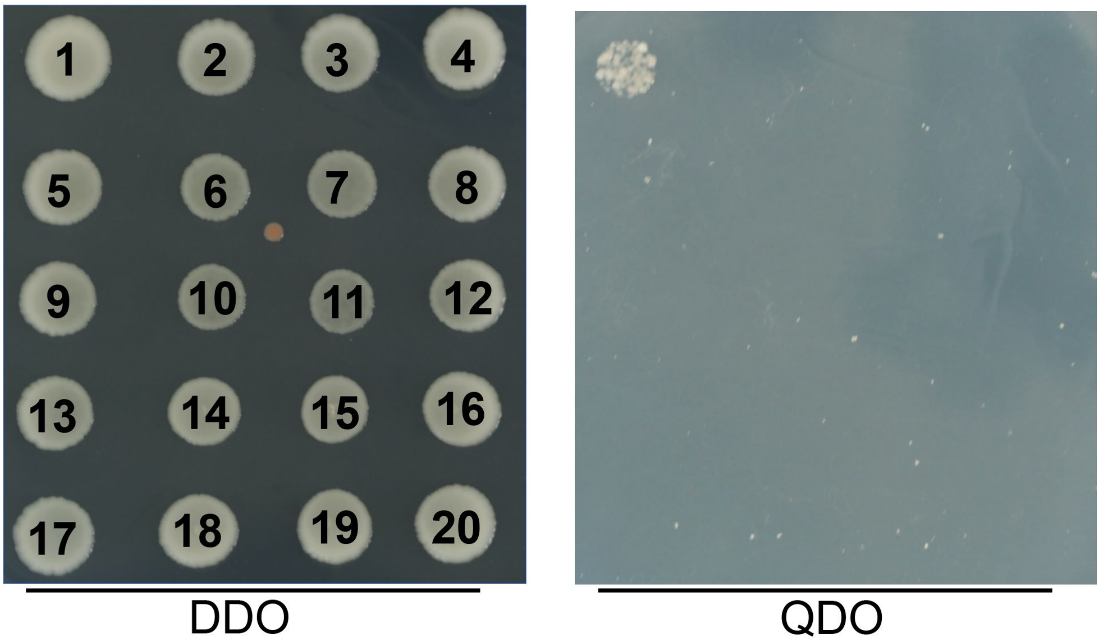


**Fig. S26. Yeast two hybrid assays showing the interaction between OsRLP4 and salivary proteins from planthopper species.** OsRLP4$_{(29\text{-}551)}$ (OsRLP4 without signal peptides and transmembrane domains) is fused to pGADT7 vector, while salivary proteins without signal peptides are fused to pGBKT7 vector, respectively. The different combinations of constructs are transformed into yeast cells, and are grown on the selective medium SD/-Trp/-Leu (DDO), and the interactions are tested with SD/-Trp/-Leu/-His/-Ade (QDO). Twenty salivary proteins are selected. The accession number of each protein is listed as follow: (1) MF278694.1, (2) XP_039291719.1, (3) KU365967.1, (4) MF278706.1, (5) MF278711.1, (6) MF278714.1, (7) MF278715.1, (8) MF278720.1, (9) XP_022195702.1, (10) XP_022196818.2, (11) XP_039284081.1, (12) XP_022192221.2, (13) XP_022207944.2, (14) KT764973,15) RZF44823.1, (16) RZF42644.1, (17) RZF33006.1, (18) RZF48570.1, (19) RZF42817.1, (20) RZF33751.1.

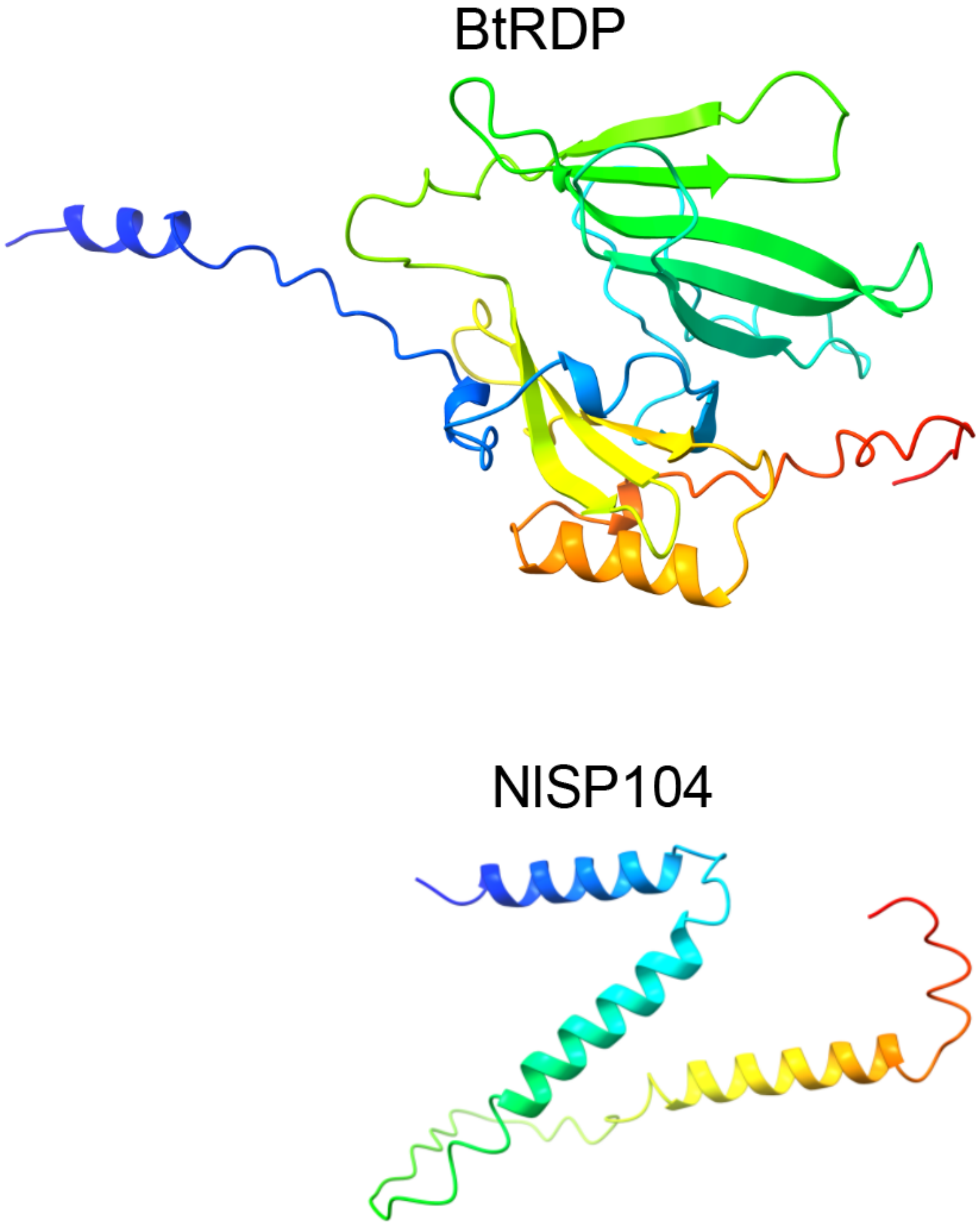


**Fig. S27. Three-dimensional structure of BtRDP and NlSP104.** AlphaFold2 is used to predicate the protein structure. The structures are colored as a spectrum from N-terminus (blue) to C-terminus (red).

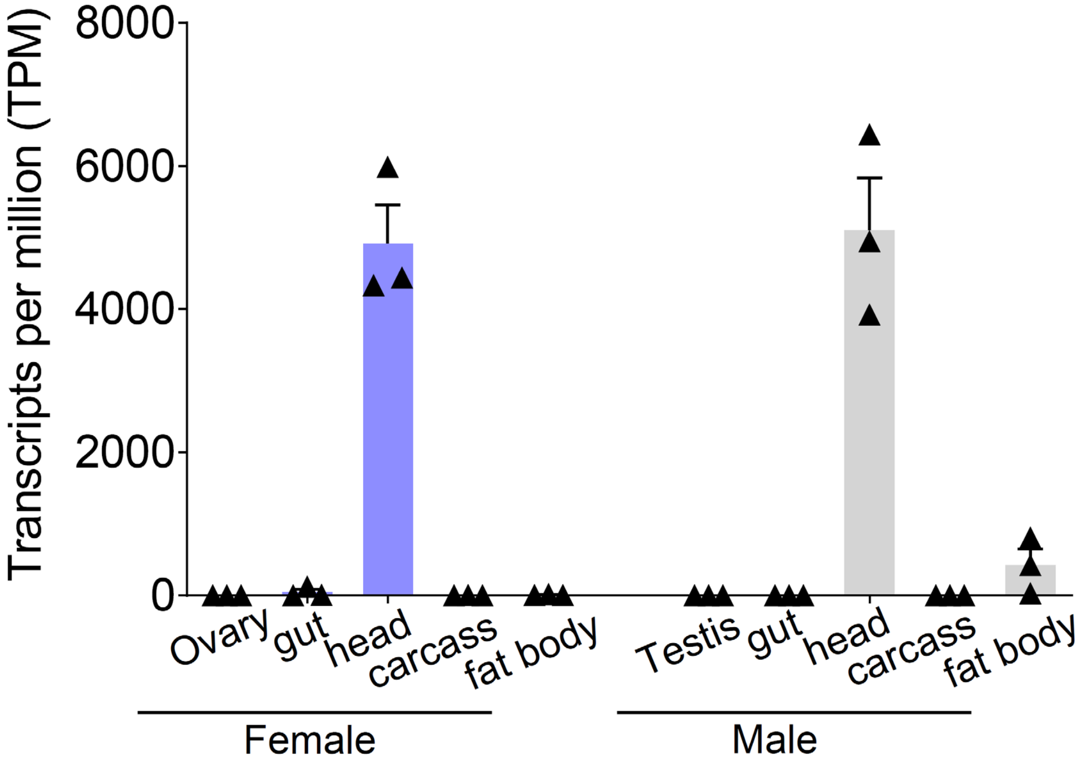


**Fig. S28. Expression patterns of *Nilaparvata lugens NlSP104*.** Transcripts per million (TPM) expression values of *NlSP104* in different tissues are determined based on the transcriptomic data. The head that contains salivary glands exhibits highest *NlSP104* expression in male and female *N. lugens*. Data are presented as mean values ± SEM (n=3 independent biological replicates).

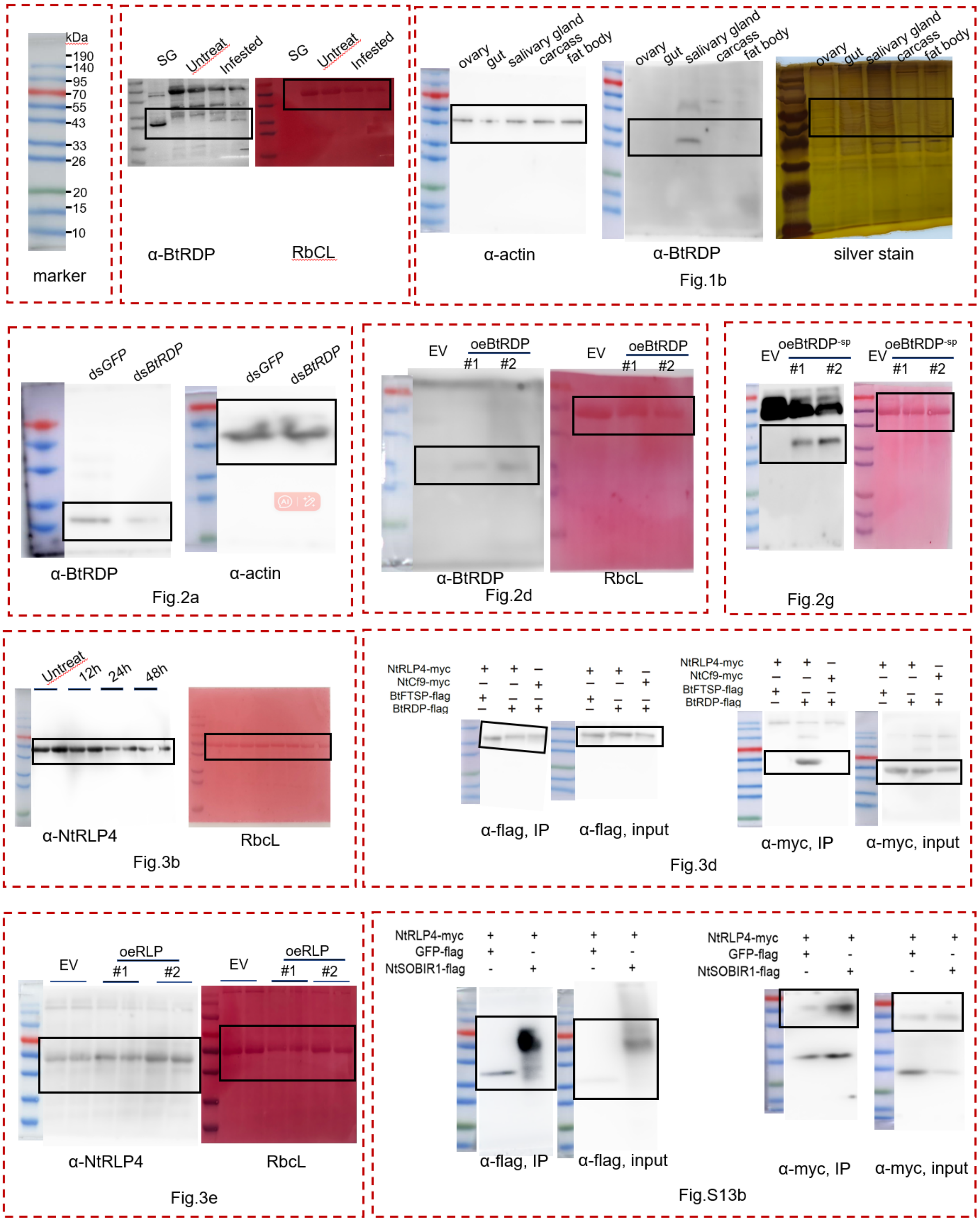


**Fig. S29.** Original images for blots in Figures 1-3 and Supplementary Figure S13b

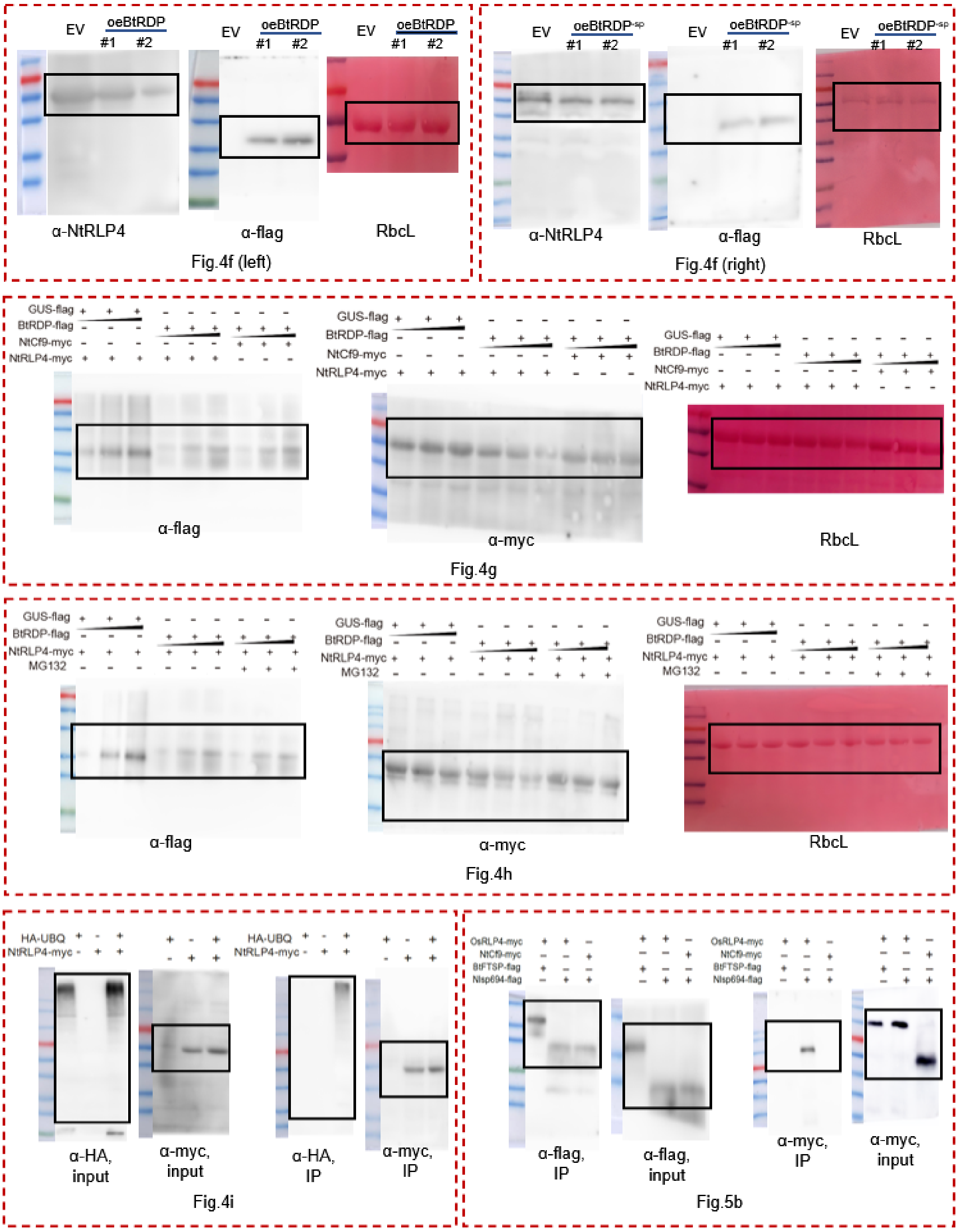


**Fig. S30.** Original images for blots in Figures 4-5

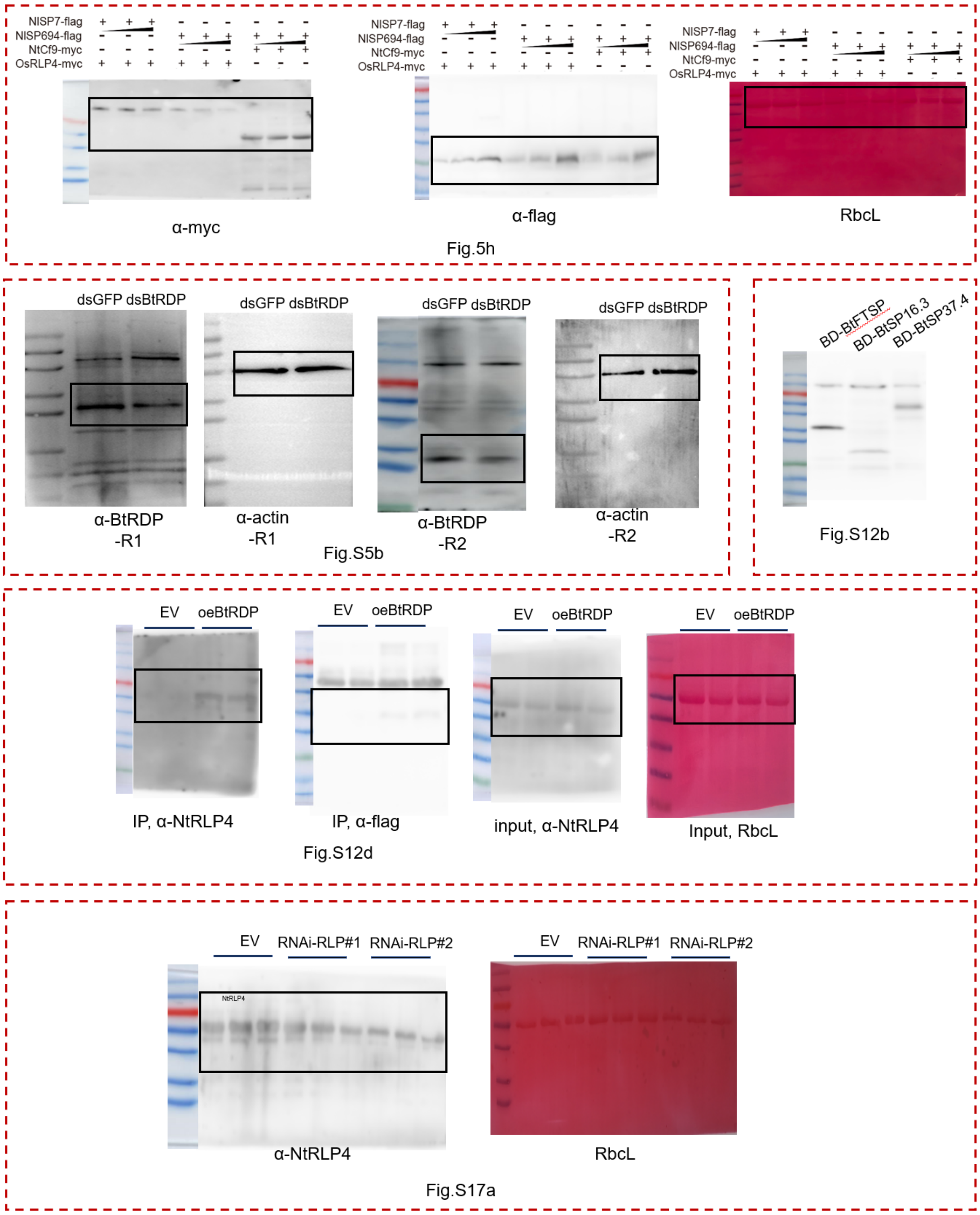


**Fig. S31.** Original images for blots in Figures 5 and Supplementary Figure 5-17.

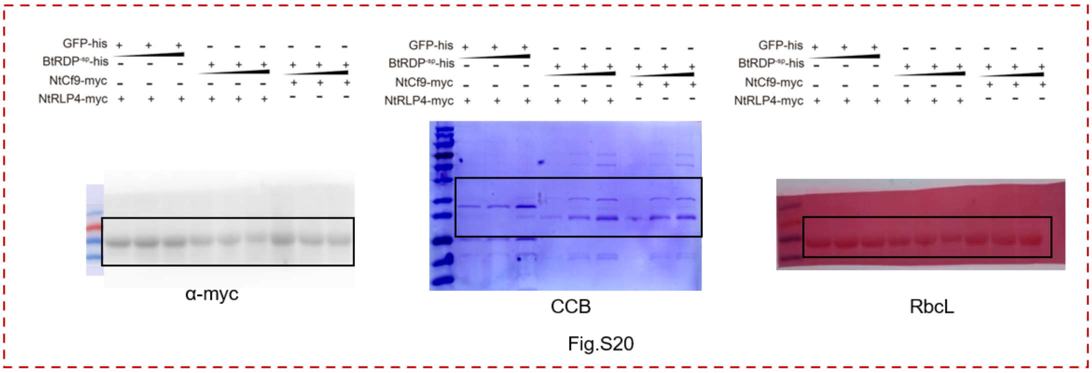


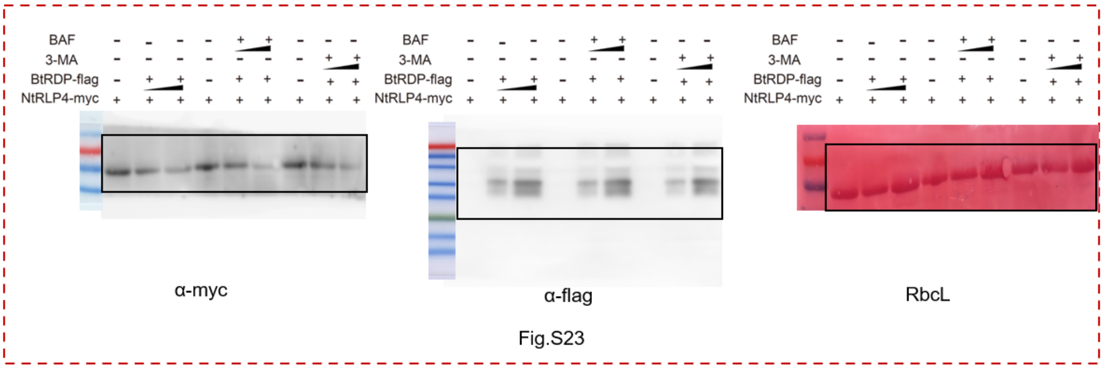


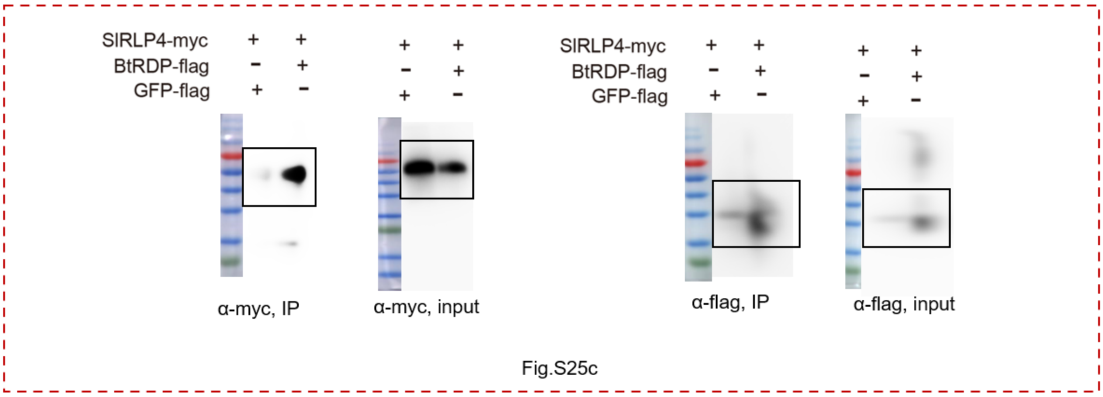


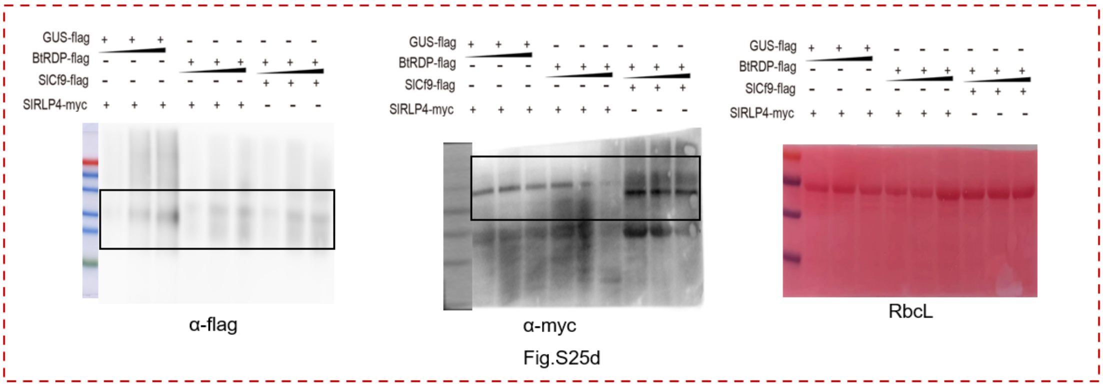


**Fig. S32.** Original images for blots in Supplementary Figure 20-25.

#### 2. Supplementary Tables

**Table S1. Identification of RDP and SP101 homologs in insect species**

| Family | Species | Accession | Total Base | Number of RDP homologs | Number of SP101 homologs | Note |
|---|---|---|---|---|---|---|
| Aleyrodidae | *Bemisia tabaci* MED | GCA_918797505 | 609.6 MB | 1 | 0 | Assembled genome |
| | *Bemisia tabaci* MEAM1 | GCF_001854935 | 600.6 MB | 1 | 0 | Assembled genome |
| | *Bemisia tabaci* India | SRR1159209 | 9.7 GB* | 1 | 0 | Assembled genome |
| | *Bemisia tabaci* SSA1 | GCA_902825415 | 657.8 MB | 1 | 0 | Assembled genome |
| | *Bemisia tabaci* SSA2 | GCA_903994125 | 625.3 MB | 1 | 0 | Assembled genome |
| | *Aleurocanthus spiniferus* | SRR17330024 | 9.2 GB* | 1 | 0 | RNA-seq, whole body |
| | *Aleyrodes proletella* | SRX14998536 | 1.9 GB* | 1 | 0 | RNA-Seq, whole body |
| | *Trialeurodes vaporariorum* | GCA_011764245 | 814.7 MB | 1 | 0 | Assembled genome |
| | *Dialeurodes citri* | SRR2980521 | 5.0 GB* | 1 | 0 | RNA-Seq, whole body |
| Delphacidae | *Nilaparvata lugens* | GCF_014356525 | 1,088 MB | 0 | 1 | Assembled genome |
| | *Laodelphax striatellus* | GCA_017141395 | 510.2 MB | 0 | 1 | Assembled genome |
| | *Sogatella furcifera* | GCA_017141385 | 563.8 MB | 0 | 1 | Assembled genome |
| Aphididae | *Acyrthosiphon pisum* | GCA_005508785 | 533.6 MB | 0 | 0 | Assembled genome |

| | | | | | | |
|---|---|---|---|---|---|---|
| | *Myzus persicae* | GCA_001856785 | 347.3 MB | 0 | 0 | Assembled genome |
| | *Sitobion miscanthi* | GCA_008086715 | 397.9 MB | 0 | 0 | Assembled genome |
| Aphalaridae | *Pachypsylla venusta* | GCA_012654025 | 482.0 MB | 0 | 0 | Assembled genome |
| | *Diaci psyllid* | GCA_000475195 | 485.7 MB | 0 | 0 | Assembled genome |
| Alydidae | *Riptortus pedestris* | GCA_019009955 | 1.0 GB | 0 | 0 | Assembled genome |
| Lygaeidae | *Oncopeltus fasciatus* | GCA_000696205 | 1.1 GB | 0 | 0 | Assembled genome |
| Pentatomidae | *Halyomorpha halys* | GCA_000696795 | 998.2 MB | 0 | 0 | Assembled genome |
| Cimicidae | *Cimex lectularius* | GCA_000648675 | 510.8 MB | 0 | 0 | Assembled genome |
| Reduviidae | *Rhodnius prolixus* | GCA_000181055 | 706.8 MB | 0 | 0 | Assembled genome |
| Gerridae | *Gerris buenoi* | GCA_001010745 | 994.4 MB | 0 | 0 | Assembled genome |
| Drosophilidae | *Drosophila melanogaster* | GCA_002310755 | 120.4 MB | 0 | 0 | Assembled genome |
| Culicidae | *Aedes albopictus* | GCA_018104305 | 1.3 GB | 0 | 0 | Assembled genome |
| Tenebrionidae | *Tribolium castaneum* | GCA_000002335 | 165.9 MB | 0 | 0 | Assembled genome |
| Chrysomeloidea | *Anoplophora glabripennis* | GCA_000390285 | 679.8 MB | 0 | 0 | Assembled genome |
| Bombycidae | *Bombyx mori* | GCA_027366755 | 456.6 MB | 0 | 0 | Assembled genome |
| Noctuidae | *Spodoptera frugiperda* | GCA_023101765 | 383.9 MB | 0 | 0 | Assembled genome |
| Pediculidae | *Pediculus humanus* | GCA_000006295 | 110.8 MB | 0 | 0 | Assembled genome |

[*] The total base of raw data retrieved from SRA database.

**Table S2. Proteins from a *Nicotiana benthamiana* cDNA library screened by yeast two hybrid using BtRDP as a bait**

| ***N. benthamiana* v2.6.1 Accession [1]** | **Annotation** | **Number of colonies** | **Region** |
|---|---|---|---|
| Niben261Chr07g1310001.1 | Putative leucine-rich repeat receptor-like serine/threonine-protein kinase | 3 | 455-709 |
| Niben261Chr13g1396009.1 | Chloroplast stem-loop binding protein of 41 kDa b | 1 | 162-374 |
| Niben261Chr02g0086007.1 | Sorbitol dehydrogenase-like | 1 | 95-305 |
| Niben261Chr17g0179002.1 | Guanine nucleotide-binding protein subunit beta-like protein | 1 | 92-326 |
| Niben261Chr01g1673009.1 | Ribulose bisphosphate carboxylase/oxygenase activase 2 | 2 | 228-420 |
| Niben261Chr07g1128067.1 | Ribulose-1,5-bisphosphate carboxylase/oxygenase small subunit | 5 | 23-181 |
| Niben261Chr14g0161005.1 | Cytochrome b6-f complex iron-sulfur subunit 2 | 5 | 1-216 |

[1] The sequence can be downloaded from Sol Genomics Network (https://solgenomics.net/ftp/genomes/Nicotiana_benthamianaV261)

**Table S3. Differentially expressed genes between empty vector (EV) and oeRLP#1 transgenic plant.**

| Gene ID | oeRLP#1 | | | | EV | | | | Description |
|---|---|---|---|---|---|---|---|---|---|
| | R1 | R2 | R3 | R4 | R1 | R2 | R3 | R4 | |
| Nitab4.5_0000757g0070 | 22 | 11 | 8 | 14 | 0 | 0 | 0 | 0 | SRC2-like protein |
| Nitab4.5_0000014g0130 | 0 | 0 | 4 | 7 | 0 | 0 | 0 | 0 | Unknown Protein |
| Nitab4.5_0001143g0060 | 3 | 6 | 1 | 0 | 0 | 0 | 0 | 0 | PBSP domain-containing protein |
| Nitab4.5_0002689g0030 | 3 | 1 | 1 | 9 | 0 | 0 | 0 | 0 | Non-structural maintenance of chromosome element 4 |
| Nitab4.5_0000171g0100 | 3 | 1 | 5 | 1 | 0 | 0 | 0 | 0 | Two-component response regulator ARR3 |
| Nitab4.5_0000344g0220 | 3 | 0 | 0 | 7 | 0 | 0 | 0 | 0 | LEA-like protein |
| Nitab4.5_0000108g0090 | 7 | 8 | 9 | 7 | 0 | 0 | 0 | 0 | Receptor serine_threonine kinase |
| Nitab4.5_0003282g0020 | 3 | 1 | 3 | 2 | 0 | 0 | 0 | 0 | RLK, Receptor like protein, putative resistance protein with an antifungal domain |
| Nitab4.5_0000287g0160 | 1 | 1 | 2 | 4 | 0 | 0 | 0 | 0 | BHLH transcription factor-like |
| Nitab4.5_0004038g0030 | 2 | 0 | 1 | 4 | 0 | 0 | 0 | 0 | -- |
| Nitab4.5_0001715g0030 | 0 | 0 | 2 | 4 | 0 | 0 | 0 | 0 | Os12g0114200 protein |
| Nitab4.5_0002380g0020 | 1 | 0 | 4 | 3 | 0 | 0 | 0 | 0 | -- |
| Nitab4.5_0000516g0160 | 5 | 1 | 0 | 5 | 0 | 0 | 1 | 0 | Dof zinc finger protein |
| Nitab4.5_0003395g0030 | 5 | 6 | 1 | 3 | 0 | 1 | 0 | 0 | SNARE associated Golgi protein |
| Nitab4.5_0002123g0050 | 2 | 1 | 1 | 2 | 0 | 0 | 0 | 0 | Beta Galactosidase-like protein |
| Nitab4.5_0001286g0080 | 2 | 2 | 2 | 0 | 0 | 0 | 0 | 0 | Alpha mannosidase-like protein |
| Nitab4.5_0001146g0240 | 2 | 1 | 1 | 2 | 0 | 0 | 0 | 0 | Unknown Protein |
| Nitab4.5_0000171g0070 | 2 | 10 | 3 | 2 | 1 | 0 | 0 | 0 | U-box domain-containing protein 13 |
| Nitab4.5_0003558g0030 | 10 | 3 | 1 | 3 | 1 | 0 | 0 | 0 | Zinc finger (Ran-binding) family protein |
| Nitab4.5_0000068g0120 | 0 | 2 | 3 | 0 | 0 | 0 | 0 | 0 | Dof zinc finger protein |
| Nitab4.5_0000725g0100 | 7 | 3 | 2 | 2 | 1 | 0 | 0 | 0 | UDP-glucosyltransferase family 1 protein |
| Nitab4.5_0000282g0040 | 4 | 2 | 2 | 4 | 0 | 1 | 0 | 0 | Cohesin subunit |
| Nitab4.5_0000458g0090 | 0 | 1 | 2 | 2 | 0 | 0 | 0 | 0 | Blue copper protein |

| | | | | | | | | | |
|---|---|---|---|---|---|---|---|---|---|
| Nitab4.5_0000856g0230 | 2 | 3 | 2 | 0 | 0 | 0 | 0 | 0 | Methyltransferase-like protein 6 |
| Nitab4.5_0002244g0110 | 5 | 5 | 4 | 0 | 0 | 0 | 0 | 1 | Isopentenyl-diphosphate delta-isomerase family protein |
| Nitab4.5_0000165g0050 | 4 | 1 | 3 | 0 | 0 | 0 | 0 | 0 | RNA-dependent RNA polymerase family protein |
| Nitab4.5_0000231g0180 | 2 | 2 | 0 | 0 | 0 | 0 | 0 | 0 | Hydrolase alpha_beta fold family protein |
| Nitab4.5_0004261g0030 | 12 | 11 | 12 | 1 | 2 | 0 | 1 | 0 | LRR receptor-like serine_threonine-protein kinase, RLP |
| Nitab4.5_0000705g0010 | 11 | 5 | 6 | 4 | 1 | 1 | 1 | 0 | Receptor like protein kinase |
| Nitab4.5_0000223g0340 | 9 | 9 | 3 | 1 | 1 | 1 | 1 | 0 | WRKY transcription factor-30 |
| Nitab4.5_0002100g0100 | 3 | 0 | 0 | 2 | 0 | 0 | 0 | 0 | N-hydroxycinnamoyl_benzoyltransferase 1 |
| Nitab4.5_0000213g0080 | 4 | 0 | 0 | 2 | 0 | 0 | 0 | 0 | Integrin-linked kinase-associated serine_threonine phosphatase 2C |
| Nitab4.5_0000274g0230 | 3 | 6 | 0 | 1 | 1 | 0 | 0 | 0 | RING finger protein 6 |
| Nitab4.5_0002177g0170 | 11 | 9 | 8 | 2 | 2 | 0 | 0 | 1 | Receptor like kinase, RLK |
| Nitab4.5_0003169g0020 | 9 | 2 | 0 | 4 | 0 | 0 | 1 | 1 | Disease resistance response |
| Nitab4.5_0000200g0030 | 1 | 7 | 4 | 2 | 0 | 0 | 0 | 1 | Aldose 1-epimerase family protein |
| Nitab4.5_0000529g0080 | 5 | 1 | 3 | 7 | 2 | 0 | 0 | 0 | Calmodulin |
| Nitab4.5_0001210g0030 | 2 | 5 | 0 | 1 | 1 | 0 | 0 | 0 | Unknown Protein |
| Nitab4.5_0003051g0020 | 25 | 22 | 23 | 5 | 5 | 0 | 1 | 2 | NBS-LRR class disease resistance protein |
| Nitab4.5_0000856g0110 | 1 | 3 | 2 | 3 | 0 | 0 | 0 | 0 | U-box domain-containing protein |
| Nitab4.5_0003558g0070 | 18 | 17 | 22 | 15 | 5 | 0 | 0 | 3 | Zinc finger (Ran-binding) family protein |
| Nitab4.5_0000495g0060 | 2 | 5 | 2 | 7 | 0 | 0 | 0 | 1 | Ring finger protein |
| Nitab4.5_0002444g0010 | 0 | 3 | 2 | 0 | 0 | 0 | 0 | 0 | -- |
| Nitab4.5_0001863g0220 | 8 | 7 | 9 | 6 | 1 | 2 | 0 | 0 | Genomic DNA chromosome 3 TAC clone K1G2 |
| Nitab4.5_0002221g0060 | 4 | 4 | 1 | 7 | 0 | 0 | 0 | 2 | Cytochrome P450 |
| Nitab4.5_0000246g0210 | 1 | 5 | 0 | 2 | 0 | 1 | 0 | 0 | Pollen allergen Phl p 11 |
| Nitab4.5_0000020g0210 | 2 | 0 | 1 | 2 | 0 | 0 | 0 | 0 | -- |
| Nitab4.5_0001794g0030 | 7 | 8 | 3 | 8 | 0 | 0 | 3 | 0 | Flavoprotein wrbA |
| Nitab4.5_0002156g0030 | 7 | 3 | 1 | 0 | 0 | 1 | 0 | 1 | Genomic DNA chromosome 5 P1 clone MQD19 |

| | | | | | | | | | |
|---|---|---|---|---|---|---|---|---|---|
| Nitab4.5_0000462g0110 | 17 | 6 | 11 | 16 | 1 | 1 | 2 | 3 | Receptor like kinase, RLK |
| Nitab4.5_0002859g0040 | 2 | 0 | 4 | 0 | 0 | 0 | 0 | 0 | Dynein light chain 1 cytoplasmic |
| Nitab4.5_0000131g0130 | 7 | 4 | 2 | 10 | 0 | 0 | 1 | 2 | LRR receptor-like serine_threonine-protein kinase, RLP |
| Nitab4.5_0000278g0050 | 2 | 2 | 1 | 6 | 0 | 0 | 1 | 1 | Os12g0581300 protein (Fragment) |
| Nitab4.5_0004821g0030 | 3 | 4 | 4 | 3 | 0 | 0 | 0 | 1 | Zinc finger family protein |
| Nitab4.5_0006338g0090 | 7 | 0 | 3 | 1 | 0 | 0 | 0 | 1 | Ring H2 finger protein |
| Nitab4.5_0002816g0080 | 6 | 1 | 2 | 3 | 1 | 0 | 0 | 1 | Os04g0461600 protein (Fragment) |
| Nitab4.5_0000028g0360 | 6 | 6 | 9 | 0 | 3 | 0 | 0 | 0 | Unknown Protein |
| Nitab4.5_0000125g0360 | 2 | 4 | 6 | 8 | 1 | 2 | 0 | 0 | Unknown Protein |
| Nitab4.5_0000200g0100 | 6 | 4 | 3 | 4 | 0 | 0 | 0 | 2 | WRKY transcription factor 23 |
| Nitab4.5_0000010g0270 | 1 | 2 | 3 | 2 | 0 | 1 | 0 | 0 | Receptor like kinase, RLK |
| Nitab4.5_0000187g0090 | 3 | 5 | 2 | 2 | 0 | 0 | 1 | 0 | Tumor susceptibility protein 101 (Fragment) |
| Nitab4.5_0000040g0670 | 2 | 2 | 0 | 1 | 0 | 0 | 0 | 0 | GHMP kinase family protein |
| Nitab4.5_0003542g0020 | 11 | 15 | 17 | 0 | 1 | 2 | 3 | 1 | Receptor like kinase, RLK |
| Nitab4.5_0000621g0030 | 15 | 21 | 6 | 27 | 0 | 3 | 6 | 2 | Glutaredoxin |
| Nitab4.5_0002114g0080 | 0 | 2 | 4 | 3 | 0 | 0 | 0 | 0 | Ras-related protein Rab-25 |
| Nitab4.5_0000098g0150 | 11 | 8 | 3 | 16 | 1 | 0 | 0 | 4 | Unknown Protein |
| Nitab4.5_0004136g0020 | 2 | 2 | 3 | 1 | 0 | 0 | 1 | 0 | |
| Nitab4.5_0004197g0020 | 2 | 3 | 4 | 33 | 4 | 2 | 0 | 1 | Patatin-like phospholipase domain-containing protein c |
| Nitab4.5_0000617g0080 | 2 | 4 | 4 | 3 | 1 | 0 | 0 | 1 | Nuclear nucleic acid-binding protein C1D |
| Nitab4.5_0001204g0110 | 2 | 4 | 1 | 1 | 0 | 1 | 0 | 0 | NAC domain transcription factor protein |
| Nitab4.5_0000825g0090 | 15 | 8 | 10 | 5 | 4 | 1 | 0 | 1 | Receptor kinase |
| Nitab4.5_0002905g0040 | 22 | 24 | 14 | 7 | 6 | 5 | 0 | 0 | Unknown Protein |
| Nitab4.5_0001662g0030 | 20 | 21 | 14 | 31 | 4 | 1 | 1 | 8 | Gibberellin 2-oxidase 2 |
| Nitab4.5_0000352g0090 | 3 | 0 | 6 | 3 | 0 | 1 | 1 | 0 | Coatomer alpha subunit-like protein |
| Nitab4.5_0000080g0230 | 1 | 2 | 3 | 2 | 0 | 0 | 0 | 1 | 5_apos-3_apos exoribonuclease 2 |

| | | | | | | | | | |
|---|---|---|---|---|---|---|---|---|---|
| Nitab4.5_0005052g0030 | 26 | 21 | 21 | 4 | 7 | 2 | 1 | 2 | Receptor like kinase, RLK |
| Nitab4.5_0000305g0140 | 2 | 3 | 1 | 4 | 0 | 0 | 1 | 0 | Ethylene-responsive transcription factor 4 |
| Nitab4.5_0000440g0150 | 2 | 5 | 2 | 5 | 1 | 0 | 1 | 0 | 2,3-bisphosphoglycerate-dependent phosphoglycerate mutase |
| Nitab4.5_0000258g0120 | 17 | 6 | 7 | 4 | 1 | 1 | 3 | 0 | Cyclic nucleotide gated channel |
| Nitab4.5_0006460g0020 | 11 | 9 | 7 | 6 | 0 | 0 | 0 | 5 | Unknown Protein |
| Nitab4.5_0000753g0120 | 1 | 3 | 4 | 3 | 0 | 0 | 1 | 1 | Unknown Protein |
| Nitab4.5_0000976g0090 | 1 | 4 | 4 | 2 | 0 | 0 | 1 | 0 | FLORICAULA_LEAFY-like protein |
| Nitab4.5_0000008g0870 | 18 | 27 | 13 | 29 | 4 | 5 | 1 | 5 | Hydroxycinnamoyl transferase |
| Nitab4.5_0001232g0030 | 7 | 19 | 2 | 13 | 0 | 3 | 4 | 0 | Unknown Protein |
| Nitab4.5_0001213g0030 | 2 | 2 | 0 | 3 | 0 | 0 | 0 | 1 | LRR receptor-like serine_threonine-protein kinase, RLP |
| Nitab4.5_0000444g0090 | 18 | 13 | 17 | 6 | 4 | 3 | 1 | 2 | Receptor-like protein kinase At3g21340 |
| Nitab4.5_0005936g0030 | 4 | 10 | 6 | 12 | 1 | 1 | 0 | 4 | F-box family protein |
| Nitab4.5_0000519g0380 | 8 | 0 | 2 | 8 | 0 | 2 | 1 | 0 | Transportin |
| Nitab4.5_0000188g0120 | 8 | 2 | 3 | 3 | 1 | 1 | 0 | 0 | Armadillo_beta-catenin repeat family protein |
| Nitab4.5_0000038g0030 | 8 | 14 | 13 | 16 | 2 | 0 | 6 | 2 | Cysteine-rich receptor-like protein kinase |
| Nitab4.5_0003514g0020 | 8 | 12 | 14 | 2 | 3 | 2 | 1 | 0 | Receptor-like protein kinase At3g21340 |
| Nitab4.5_0000344g0120 | 12 | 14 | 7 | 6 | 3 | 1 | 1 | 2 | Cell division protease ftsH homolog 3 |
| Nitab4.5_0000041g0370 | 2 | 2 | 2 | 1 | 0 | 0 | 1 | 0 | Acetyl esterase |
| Nitab4.5_0003443g0050 | 3 | 4 | 2 | 1 | 0 | 0 | 0 | 1 | Hypoxanthine phosphoribosyltransferase |
| Nitab4.5_0000249g0390 | 1 | 3 | 1 | 3 | 0 | 0 | 0 | 1 | Homeodomain-like |
| Nitab4.5_0000517g0040 | 9 | 4 | 4 | 2 | 1 | 1 | 0 | 1 | ELF4-like protein |
| Nitab4.5_0001438g0020 | 5 | 4 | 3 | 1 | 0 | 1 | 1 | 0 | Receptor-like protein kinase |
| Nitab4.5_0002177g0100 | 1 | 2 | 3 | 1 | 0 | 0 | 1 | 0 | Receptor like kinase, RLK |
| Nitab4.5_0002783g0040 | 1 | 2 | 2 | 6 | 0 | 0 | 0 | 2 | Polygalacturonase |
| Nitab4.5_0004728g0070 | 3 | 1 | 1 | 3 | 0 | 0 | 0 | 1 | Mitochondrial phosphate carrier protein |
| Nitab4.5_0000286g0030 | 6 | 5 | 4 | 2 | 1 | 1 | 1 | 0 | Ankyrin repeat protein |

| | | | | | | | | | |
|---|---|---|---|---|---|---|---|---|---|
| Nitab4.5_0003558g0090 | 16 | 0 | 18 | 3 | 1 | 0 | 5 | 2 | Zinc finger (Ran-binding) family protein |
| Nitab4.5_0000716g0240 | 9 | 2 | 9 | 4 | 2 | 3 | 0 | 0 | Unknown Protein |
| Nitab4.5_0001896g0050 | 2 | 1 | 3 | 2 | 0 | 1 | 0 | 0 | Os06g0220000 protein (Fragment) |
| Nitab4.5_0000082g0400 | 2 | 2 | 6 | 20 | 2 | 1 | 1 | 2 | Patatin-like phospholipase domain-containing protein c |
| Nitab4.5_0006826g0060 | 3 | 12 | 6 | 5 | 2 | 3 | 1 | 0 | |
| Nitab4.5_0003551g0110 | 5 | 3 | 4 | 19 | 4 | 1 | 0 | 1 | Unknown Protein |
| Nitab4.5_0001302g0090 | 3 | 1 | 1 | 2 | 1 | 1 | 0 | 0 | Glycosyltransferase-like protein |
| Nitab4.5_0000462g0120 | 13 | 14 | 10 | 18 | 5 | 1 | 3 | 2 | Receptor like kinase, RLK |
| Nitab4.5_0001608g0050 | 6 | 10 | 14 | 4 | 4 | 1 | 1 | 1 | Cathepsin B-like cysteine proteinase |
| Nitab4.5_0001048g0080 | 0 | 2 | 2 | 1 | 0 | 1 | 0 | 0 | ATP-dependent RNA helicase |
| Nitab4.5_0000404g0060 | 12 | 7 | 6 | 16 | 2 | 2 | 1 | 3 | Unknown Protein |
| Nitab4.5_0001461g0070 | 23 | 17 | 6 | 6 | 5 | 2 | 1 | 2 | cytochrome P450 |
| Nitab4.5_0004692g0020 | 2 | 2 | 5 | 4 | 0 | 0 | 2 | 1 | RING zinc finger-containing protein |
| Nitab4.5_0004732g0010 | 8 | 11 | 5 | 1 | 4 | 1 | 1 | 0 | Receptor-like kinase |
| Nitab4.5_0000262g0190 | 1 | 1 | 3 | 9 | 1 | 0 | 1 | 1 | Pectinesterase |
| Nitab4.5_0003913g0040 | 10 | 9 | 6 | 10 | 1 | 1 | 2 | 3 | Subtilisin-like protease |
| Nitab4.5_0000153g0200 | 3 | 5 | 2 | 5 | 0 | 1 | 3 | 0 | |
| Nitab4.5_0000747g0020 | 2 | 3 | 6 | 1 | 1 | 1 | 0 | 1 | Serine_threonine-protein kinase receptor |
| Nitab4.5_0000232g0330 | 4 | 1 | 2 | 0 | 1 | 1 | 0 | 0 | Receptor like kinase, RLK |
| Nitab4.5_0000174g0170 | 24 | 30 | 12 | 14 | 12 | 2 | 4 | 1 | Unknown Protein |
| Nitab4.5_0000370g0110 | 1 | 0 | 3 | 2 | 0 | 0 | 0 | 0 | Aldose 1-epimerase family protein |
| Nitab4.5_0001439g0050 | 5 | 2 | 3 | 3 | 1 | 1 | 0 | 1 | Cytochrome P450 |
| Nitab4.5_0000262g0080 | 2 | 6 | 5 | 5 | 2 | 1 | 0 | 1 | 8-amino-7-oxononanoate synthase-like protein |
| Nitab4.5_0004800g0070 | 3 | 3 | 1 | 3 | 2 | 0 | 1 | 0 | F-box_kelch-repeat protein At1g22040 |
| Nitab4.5_0000980g0260 | 0 | 2 | 3 | 2 | 1 | 0 | 0 | 0 | Neutral invertase like protein |
| Nitab4.5_0002285g0010 | 8 | 5 | 5 | 3 | 1 | 1 | 2 | 1 | Nodulin-like protein (Fragment) |

| | | | | | | | | | |
|---|---|---|---|---|---|---|---|---|---|
| Nitab4.5_0004506g0120 | 10 | 7 | 11 | 3 | 3 | 1 | 1 | 2 | Cyclic nucleotide gated channel |
| Nitab4.5_0004657g0030 | 68 | 50 | 39 | 150 | 20 | 10 | 21 | 23 | Auxin response factor 14 |
| Nitab4.5_0002328g0020 | 15 | 16 | 17 | 13 | 6 | 4 | 1 | 3 | Heat stress transcription factor-type, DNA-binding |
| Nitab4.5_0000676g0220 | 3 | 0 | 3 | 3 | 0 | 1 | 1 | 1 | Myb family transcription factor |
| Nitab4.5_0002321g0060 | 5 | 7 | 3 | 4 | 1 | 0 | 3 | 0 | NAC domain protein IPR003441 protein |
| Nitab4.5_0000892g0040 | 3 | 2 | 3 | 1 | 0 | 0 | 0 | 1 | Receptor-like protein kinase |
| Nitab4.5_0000146g0180 | 5 | 5 | 1 | 2 | 2 | 1 | 0 | 0 | Oligopeptidase (Protease II) |
| Nitab4.5_0002916g0060 | 10 | 10 | 6 | 14 | 4 | 1 | 4 | 1 | 1-aminocyclopropane-1-carboxylate oxidase-like protein |
| Nitab4.5_0000441g0240 | 12 | 6 | 5 | 8 | 4 | 2 | 0 | 2 | Membrane protein |
| Nitab4.5_0004625g0010 | 5 | 3 | 6 | 1 | 2 | 1 | 0 | 0 | Serine_threonine-protein kinase receptor |
| Nitab4.5_0001143g0050 | 45 | 43 | 25 | 52 | 3 | 9 | 10 | 20 | PBSP domain-containing protein |
| Nitab4.5_0004490g0050 | 4 | 2 | 2 | 3 | 0 | 1 | 0 | 1 | 3-deoxy-D-manno-octulosonic acid transferase-like protein |
| Nitab4.5_0000028g0180 | 134 | 78 | 91 | 41 | 49 | 14 | 14 | 11 | Blue copper protein (Fragment) |
| Nitab4.5_0000989g0020 | 12 | 9 | 5 | 7 | 2 | 1 | 2 | 3 | Glutamate-gated kainate-type ion channel receptor subunit GluR5 |
| Nitab4.5_0002222g0070 | 8 | 6 | 1 | 13 | 1 | 4 | 2 | 0 | F-box family protein |
| Nitab4.5_0002796g0030 | 13 | 18 | 10 | 11 | 0 | 9 | 0 | 5 | Unknown Protein |
| Nitab4.5_0000462g0170 | 2 | 2 | 3 | 0 | 1 | 0 | 0 | 0 | Receptor like kinase, RLK |
| Nitab4.5_0004464g0010 | 4 | 1 | 2 | 4 | 1 | 0 | 1 | 1 | Cytochrome P450 |
| Nitab4.5_0002251g0110 | 83 | 40 | 58 | 39 | 25 | 8 | 11 | 14 | NBS-LRR class disease resistance protein |
| Nitab4.5_0000287g0290 | 10 | 4 | 4 | 3 | 1 | 2 | 2 | 1 | Cytochrome P450 |
| Nitab4.5_0001143g0020 | 57 | 71 | 31 | 152 | 23 | 11 | 15 | 32 | PBSP domain-containing protein |
| Nitab4.5_0001157g0010 | 12 | 4 | 6 | 2 | 2 | 2 | 0 | 3 | Ribosomal-protein-alanine N-acetyltransferase |
| Nitab4.5_0001160g0190 | 5 | 5 | 3 | 5 | 0 | 1 | 2 | 1 | Calcium-binding protein 39 |
| Nitab4.5_0001386g0050 | 6 | 11 | 9 | 29 | 2 | 6 | 0 | 6 | Wound induced protein |
| Nitab4.5_0001423g0140 | 4 | 1 | 2 | 2 | 0 | 2 | 1 | 1 | Phosphoglycerate mutase family protein |
| Nitab4.5_0003063g0040 | 1 | 3 | 5 | 7 | 1 | 1 | 2 | 0 | Dof zinc finger protein 6 |

| | | | | | | | | | |
|---|---|---|---|---|---|---|---|---|---|
| Nitab4.5_0000235g0130 | 59 | 40 | 19 | 13 | 17 | 5 | 3 | 10 | 3-hydroxy-3-methylglutaryl coenzyme A reductase |
| Nitab4.5_0004955g0020 | 3 | 1 | 2 | 9 | 1 | 0 | 2 | 2 | Homeobox leucine zipper protein |
| Nitab4.5_0000231g0240 | 5 | 2 | 2 | 18 | 3 | 3 | 1 | 0 | Ninja-family protein 1 |
| Nitab4.5_0001816g0090 | 4 | 4 | 2 | 13 | 1 | 1 | 2 | 3 | AT-hook DNA-binding protein (Fragment) |
| Nitab4.5_0002405g0110 | 1 | 3 | 2 | 1 | 0 | 1 | 1 | 1 | Tubulin beta chain |
| Nitab4.5_0000008g0370 | 5 | 2 | 2 | 15 | 2 | 0 | 1 | 3 | PAR-1c protein |
| Nitab4.5_0003912g0030 | 3 | 1 | 3 | 4 | 2 | 1 | 0 | 1 | Dihydroflavonol 4-reductase family-binding domain |
| Nitab4.5_0001087g0030 | 72 | 54 | 46 | 34 | 23 | 9 | 8 | 16 | Unknown Protein |
| Nitab4.5_0002817g0050 | 0 | 5 | 2 | 4 | 1 | 0 | 1 | 1 | Heat shock-like protein |
| Nitab4.5_0000057g0070 | 10 | 9 | 9 | 4 | 3 | 1 | 4 | 1 | Receptor-like kinase |
| Nitab4.5_0001952g0050 | 4 | 7 | 4 | 2 | 1 | 1 | 2 | 0 | NHL repeat-containing protein-like |
| Nitab4.5_0002331g0010 | 3 | 3 | 11 | 0 | 1 | 2 | 2 | 0 | Peroxidase 5 |
| Nitab4.5_0000130g0140 | 126 | 71 | 48 | 71 | 64 | 6 | 13 | 7 | 1-aminocyclopropane-1-carboxylate oxidase |
| Nitab4.5_0000970g0150 | 10 | 2 | 3 | 5 | 3 | 0 | 2 | 0 | Galactokinase like protein |
| Nitab4.5_0002356g0140 | 4 | 3 | 3 | 1 | 1 | 1 | 1 | 0 | Receptor like kinase, RLK |
| Nitab4.5_0006034g0010 | 6 | 7 | 10 | 3 | 4 | 2 | 0 | 1 | Cc-nbs-lrr, resistance protein |
| Nitab4.5_0002337g0050 | 4 | 6 | 5 | 3 | 3 | 1 | 0 | 1 | Oxidoreductase 2OG-Fe oxygenase family protein |
| Nitab4.5_0000188g0090 | 39 | 72 | 46 | 85 | 17 | 11 | 17 | 25 | Chaperone protein dnaJ 11 |
| Nitab4.5_0001863g0230 | 7 | 8 | 5 | 7 | 2 | 1 | 2 | 2 | Ring zinc finger protein (Fragment) |
| Nitab4.5_0000391g0120 | 8 | 6 | 3 | 4 | 2 | 2 | 2 | 1 | Unknown Protein |
| Nitab4.5_0001777g0020 | 72 | 118 | 132 | 21 | 58 | 19 | 12 | 11 | Os06g0524700 protein (Fragment) |
| Nitab4.5_0000864g0100 | 4 | 3 | 3 | 5 | 0 | 2 | 1 | 1 | Cold-shock DNA binding protein |
| Nitab4.5_0001461g0050 | 65 | 34 | 17 | 32 | 16 | 7 | 12 | 9 | Alpha-humulene_(-)-(E)-beta-caryophyllene synthase |
| Nitab4.5_0000118g0080 | 27 | 9 | 10 | 26 | 4 | 5 | 5 | 7 | Cytochrome P450 |
| Nitab4.5_0000795g0070 | 41 | 23 | 17 | 24 | 9 | 7 | 9 | 7 | Solute carrier family 2, facilitated glucose transporter member 3 |
| Nitab4.5_0001231g0050 | 35 | 42 | 43 | 88 | 17 | 12 | 11 | 20 | Ethylene responsive transcription factor 2b |

| | | | | | | | | | |
|---|---|---|---|---|---|---|---|---|---|
| Nitab4.5_0000017g0160 | 14 | 12 | 5 | 6 | 0 | 1 | 5 | 5 | WRKY transcription factor 2 |
| Nitab4.5_0000262g0170 | 2 | 1 | 3 | 4 | 0 | 1 | 1 | 1 | Glycerol-3-phosphate transporter |
| Nitab4.5_0001270g0170 | 52 | 44 | 38 | 97 | 18 | 17 | 16 | 18 | Atcambp25-binding protein OF |
| Nitab4.5_0000212g0200 | 2 | 2 | 2 | 2 | 1 | 1 | 0 | 0 | Unknown Protein |
| Nitab4.5_0000655g0030 | 15 | 11 | 13 | 6 | 3 | 3 | 5 | 2 | NaCl-inducible Ca2+-binding protein |
| Nitab4.5_0001223g0090 | 10 | 5 | 5 | 4 | 3 | 1 | 1 | 2 | Serine_threonine-protein kinase receptor |
| Nitab4.5_0000013g0280 | 72 | 63 | 43 | 54 | 13 | 13 | 18 | 27 | Kinase family protein |
| Nitab4.5_0000170g0410 | 10 | 4 | 4 | 3 | 3 | 1 | 0 | 2 | Low affinity zinc transporter |
| Nitab4.5_0003558g0080 | 3 | 6 | 13 | 9 | 2 | 2 | 1 | 4 | Zinc finger (Ran-binding) family protein |
| Nitab4.5_0000230g0020 | 5 | 7 | 6 | 6 | 3 | 2 | 1 | 1 | Receptor like protein kinase |
| Nitab4.5_0000188g0200 | 20 | 17 | 5 | 12 | 5 | 3 | 5 | 3 | Cytochrome P450 |
| Nitab4.5_0000635g0040 | 109 | 66 | 50 | 59 | 29 | 14 | 18 | 26 | Epoxide hydrolase 3 |
| Nitab4.5_0002131g0040 | 3 | 4 | 2 | 1 | 0 | 1 | 1 | 0 | Calcium dependent protein kinase 2 |
| Nitab4.5_0000944g0090 | 7 | 5 | 2 | 5 | 1 | 1 | 3 | 1 | Glutamate decarboxylase |
| Nitab4.5_0000403g0140 | 4 | 2 | 2 | 2 | 0 | 0 | 1 | 2 | Alcohol dehydrogenase zinc-containing |
| Nitab4.5_0001492g0080 | 16 | 10 | 10 | 4 | 4 | 3 | 4 | 1 | Receptor like kinase, RLK |
| Nitab4.5_0000563g0210 | 45 | 27 | 23 | 23 | 20 | 5 | 4 | 7 | Unknown Protein |
| Nitab4.5_0002776g0050 | 26 | 19 | 19 | 16 | 11 | 8 | 1 | 5 | Glutathione peroxidase |
| Nitab4.5_0000342g0270 | 9 | 10 | 6 | 15 | 4 | 3 | 2 | 4 | Ethylene receptor |
| Nitab4.5_0000008g0460 | 2 | 2 | 4 | 2 | 0 | 2 | 0 | 1 | Os03g0169000 protein (Fragment) |
| Nitab4.5_0001030g0120 | 4 | 4 | 9 | 2 | 3 | 1 | 2 | 0 | Microtubule-associated protein TORTIFOLIA1 |
| Nitab4.5_0000006g0300 | 40 | 35 | 23 | 17 | 10 | 6 | 11 | 9 | WRKY transcription factor |
| Nitab4.5_0000381g0130 | 44 | 47 | 116 | 12 | 27 | 20 | 20 | 3 | WRKY transcription factor 6 |
| Nitab4.5_0000742g0110 | 9 | 8 | 9 | 26 | 5 | 5 | 0 | 7 | Chaperone protein dnaJ |
| Nitab4.5_0000059g0360 | 5 | 1 | 4 | 5 | 0 | 2 | 0 | 2 | Glucose transporter 8 |
| Nitab4.5_0001014g0030 | 4 | 2 | 1 | 4 | 1 | 0 | 1 | 1 | Cytochrome P450 |

| | | | | | | | | | |
|---|---|---|---|---|---|---|---|---|---|
| Nitab4.5_0002265g0140 | 8 | 12 | 14 | 6 | 3 | 3 | 7 | 1 | cDNA clone J023121M11 full insert sequence |
| Nitab4.5_0002241g0040 | 8 | 7 | 4 | 6 | 2 | 2 | 4 | 0 | Universal stress protein |
| Nitab4.5_0000117g0060 | 2 | 12 | 4 | 9 | 3 | 2 | 2 | 2 | Unknown Protein |
| Nitab4.5_0000057g0260 | 19 | 21 | 20 | 23 | 8 | 7 | 3 | 8 | Transmembrane BAX inhibitor motif-containing protein 4 |
| Nitab4.5_0000443g0080 | 2 | 2 | 1 | 2 | 0 | 1 | 0 | 0 | LRR receptor-like serine_threonine-protein kinase, RLP |
| Nitab4.5_0002022g0010 | 2 | 2 | 1 | 1 | 1 | 0 | 1 | 0 | Ternary complex factor MIP1 |
| Nitab4.5_0001711g0050 | 7 | 7 | 9 | 80 | 8 | 9 | 9 | 8 | Non-specific lipid-transfer protein |
| Nitab4.5_0003766g0010 | 28 | 63 | 48 | 72 | 19 | 21 | 21 | 10 | Plant-specific domain TIGR01615 family protein |
| Nitab4.5_0001691g0140 | 9 | 9 | 7 | 9 | 6 | 3 | 1 | 1 | Cation diffusion facilitator family transporter |
| Nitab4.5_0001575g0050 | 17 | 8 | 6 | 9 | 7 | 1 | 4 | 1 | U-box domain-containing protein 5 |
| Nitab4.5_0003924g0080 | 7 | 7 | 6 | 10 | 3 | 3 | 3 | 2 | AT5g06970_MOJ9_14 |
| Nitab4.5_0001494g0140 | 7 | 7 | 5 | 7 | 1 | 0 | 4 | 5 | Chaperone protein dnaJ |
| Nitab4.5_0002765g0040 | 5 | 2 | 1 | 1 | 1 | 1 | 0 | 0 | Histone-lysine N-methyltransferase MEDEA |
| Nitab4.5_0002116g0040 | 4 | 3 | 1 | 2 | 0 | 1 | 0 | 2 | Chaperone protein dnaJ |
| Nitab4.5_0001536g0020 | 16 | 8 | 10 | 9 | 8 | 2 | 3 | 2 | Receptor-like protein kinase At3g21340 |
| Nitab4.5_0002682g0060 | 11 | 9 | 6 | 47 | 5 | 4 | 3 | 13 | Alpha-galactosidase |
| Nitab4.5_0002219g0040 | 91 | 23 | 36 | 29 | 15 | 15 | 14 | 16 | LRR receptor-like serine_threonine-protein kinase, RLP |
| Nitab4.5_0001654g0010 | 36 | 43 | 27 | 32 | 16 | 9 | 11 | 11 | Hydroxycinnamoyl transferase |
| Nitab4.5_0007184g0020 | 7 | 7 | 8 | 8 | 2 | 2 | 4 | 2 | Pathogen-induced calmodulin-binding protein (Fragment) |
| Nitab4.5_0004625g0020 | 7 | 4 | 2 | 8 | 1 | 3 | 3 | 1 | CONSTANS-like zinc finger protein |
| Nitab4.5_0005043g0060 | 12 | 6 | 16 | 3 | 3 | 4 | 3 | 2 | Glutaredoxin |
| Nitab4.5_0000430g0110 | 6 | 11 | 9 | 13 | 8 | 1 | 1 | 3 | CHP-rich zinc finger protein-like |
| Nitab4.5_0000209g0240 | 2 | 2 | 2 | 7 | 1 | 1 | 0 | 2 | Genomic DNA chromosome 3 P1 clone MUJ8 |
| Nitab4.5_0001495g0020 | 5 | 8 | 3 | 3 | 1 | 2 | 1 | 3 | Unknown Protein |
| Nitab4.5_0000357g0200 | 11 | 7 | 5 | 3 | 5 | 1 | 0 | 2 | RING finger protein 38 |
| Nitab4.5_0001599g0220 | 3 | 2 | 2 | 2 | 1 | 1 | 1 | 1 | Baculoviral IAP repeat-containing 4 (Predicted) |

| | | | | | | | | | |
|---|---|---|---|---|---|---|---|---|---|
| Nitab4.5_0000462g0080 | 6 | 9 | 8 | 6 | 3 | 3 | 3 | 1 | Receptor like kinase, RLK |
| Nitab4.5_0000615g0040 | 13 | 10 | 7 | 16 | 8 | 2 | 3 | 3 | Short-chain dehydrogenase_reductase family protein |
| Nitab4.5_0000705g0190 | 4 | 1 | 2 | 1 | 1 | 1 | 1 | 1 | GRAS family transcription factor |
| Nitab4.5_0004507g0010 | 8 | 3 | 3 | 1 | 1 | 1 | 2 | 1 | Receptor-like kinase |
| Nitab4.5_0001860g0020 | 24 | 18 | 22 | 17 | 11 | 5 | 8 | 5 | Glycosyl transferase family 17 protein |
| Nitab4.5_0002088g0040 | 22 | 20 | 16 | 11 | 6 | 7 | 8 | 3 | Genomic DNA chromosome 5 P1 clone MRH10 |
| Nitab4.5_0004048g0040 | 15 | 7 | 15 | 11 | 4 | 5 | 4 | 4 | Alcohol dehydrogenase (Fragment) |
| Nitab4.5_0001553g0040 | 9 | 10 | 10 | 1 | 4 | 3 | 2 | 2 | Nbs-lrr, resistance protein |
| Nitab4.5_0004813g0010 | 23 | 15 | 12 | 5 | 4 | 8 | 2 | 6 | VQ motif family protein |
| Nitab4.5_0001714g0100 | 45 | 31 | 26 | 28 | 19 | 11 | 9 | 7 | Phospholipase D |
| Nitab4.5_0002613g0020 | 4 | 2 | 2 | 3 | 1 | 0 | 2 | 0 | Tyrosyl-DNA phosphodiesterase 1 |
| Nitab4.5_0000622g0110 | 5 | 4 | 3 | 1 | 1 | 1 | 1 | 1 | |
| Nitab4.5_0002345g0020 | 16 | 7 | 8 | 9 | 5 | 3 | 3 | 3 | RLK, Receptor like protein, putative resistance protein with an antifungal domain |
| Nitab4.5_0000250g0100 | 3 | 1 | 4 | 6 | 2 | 1 | 1 | 1 | Zinc finger FYVE domain containing 26 |
| Nitab4.5_0000652g0050 | 68 | 65 | 47 | 42 | 22 | 11 | 26 | 22 | Glutathione S-transferase-like protein |
| Nitab4.5_0001049g0140 | 16 | 24 | 52 | 14 | 6 | 13 | 9 | 10 | Pistil extensin like protein (Fragment) |
| Nitab4.5_0000705g0020 | 14 | 8 | 11 | 12 | 5 | 3 | 4 | 4 | Receptor like protein kinase |
| Nitab4.5_0002780g0130 | 46 | 36 | 40 | 26 | 22 | 9 | 12 | 11 | U-box domain-containing protein |
| Nitab4.5_0000962g0010 | 9 | 6 | 4 | 6 | 2 | 1 | 4 | 3 | UDP-glucuronosyltransferase 1-1 |
| Nitab4.5_0001874g0010 | 8 | 6 | 3 | 4 | 3 | 1 | 2 | 2 | NPR1-like protein (Fragment) |
| Nitab4.5_0005528g0080 | 13 | 11 | 9 | 6 | 3 | 5 | 2 | 5 | Receptor-like kinase |
| Nitab4.5_0002232g0030 | 106 | 84 | 51 | 203 | 44 | 26 | 47 | 50 | Unknown Protein |
| Nitab4.5_0003885g0100 | 3 | 2 | 2 | 2 | 0 | 0 | 2 | 1 | Kelch-like protein |
| Nitab4.5_0000175g0060 | 5 | 3 | 4 | 1 | 2 | 0 | 1 | 2 | NB-ARC domain containing protein expressed |
| Nitab4.5_0000250g0040 | 4 | 5 | 5 | 2 | 1 | 2 | 2 | 1 | Os10g0422600 protein (Fragment) |
| Nitab4.5_0000882g0090 | 14 | 26 | 15 | 23 | 14 | 3 | 6 | 7 | Protein transport protein Sec61 beta subunit |

| | | | | | | | | | |
|---|---|---|---|---|---|---|---|---|---|
| Nitab4.5_0001926g0080 | 15 | 10 | 12 | 34 | 5 | 4 | 8 | 9 | Exoribonuclease R |
| Nitab4.5_0001553g0030 | 4 | 7 | 3 | 1 | 2 | 3 | 1 | 1 | Cc-nbs-lrr, resistance protein |
| Nitab4.5_0000575g0130 | 85 | 64 | 85 | 356 | 54 | 44 | 48 | 80 | Aquaporin 2 |
| Nitab4.5_0000134g0180 | 6 | 26 | 8 | 16 | 5 | 4 | 8 | 5 | F-box protein PP2-B1 |
| Nitab4.5_0001326g0060 | 4 | 5 | 4 | 6 | 2 | 1 | 3 | 1 | Pyrimidine 5_apos-nucleotidase |
| Nitab4.5_0000465g0060 | 17 | 10 | 13 | 7 | 8 | 3 | 7 | 1 | Armadillo_beta-catenin repeat family protein |
| Nitab4.5_0000294g0190 | 8 | 10 | 6 | 6 | 4 | 3 | 2 | 3 | COP9 signalosome subunit 6 |
| Nitab4.5_0005325g0010 | 6 | 9 | 9 | 7 | 0 | 6 | 3 | 2 | Unknown Protein |
| Nitab4.5_0004529g0010 | 292 | 211 | 140 | 155 | 96 | 54 | 75 | 85 | 3-hydroxy-3-methylglutaryl coenzyme A reductase |
| Nitab4.5_0000082g0090 | 2 | 4 | 3 | 3 | 2 | 1 | 0 | 1 | Receptor like kinase, RLK |
| Nitab4.5_0000005g0240 | 4 | 3 | 5 | 5 | 2 | 1 | 1 | 2 | U3 small nucleolar RNA-associated protein 6 homolog |
| Nitab4.5_0001030g0010 | 1 | 2 | 3 | 2 | 0 | 1 | 1 | 1 | Tir-nbs-lrr, resistance protein |
| Nitab4.5_0000293g0110 | 3 | 5 | 2 | 2 | 2 | 1 | 1 | 0 | MORN repeat-containing protein |
| Nitab4.5_0000429g0030 | 60 | 62 | 36 | 222 | 43 | 30 | 32 | 45 | ABC transporter G family member 22 |
| Nitab4.5_0003290g0010 | 8 | 5 | 3 | 2 | 2 | 3 | 0 | 2 | Calcium-dependent protein kinase |
| Nitab4.5_0000737g0060 | 25 | 20 | 10 | 45 | 13 | 9 | 8 | 10 | Hydroxycinnamoyl transferase |
| Nitab4.5_0000987g0110 | 6 | 4 | 8 | 6 | 4 | 1 | 3 | 2 | Mitochondrial glycoprotein family protein |
| Nitab4.5_0000044g0240 | 48 | 49 | 57 | 33 | 32 | 18 | 13 | 12 | Inorganic phosphate transporter |
| Nitab4.5_0000336g0020 | 79 | 47 | 37 | 62 | 22 | 23 | 30 | 16 | Universal stress protein |
| Nitab4.5_0000040g0440 | 4 | 4 | 12 | 3 | 1 | 2 | 3 | 3 | Lipase |
| Nitab4.5_0000775g0070 | 39 | 30 | 38 | 46 | 11 | 20 | 13 | 18 | ATP dependent RNA helicase |
| Nitab4.5_0000198g0120 | 29 | 31 | 31 | 31 | 8 | 11 | 15 | 15 | Flavin-binding kelch domain F box protein |
| Nitab4.5_0001357g0030 | 19 | 31 | 32 | 37 | 9 | 9 | 11 | 19 | Aluminum-induced protein-like |
| Nitab4.5_0002548g0020 | 3 | 1 | 4 | 1 | 1 | 1 | 2 | 1 | Receptor like kinase, RLK |
| Nitab4.5_0000397g0280 | 18 | 10 | 11 | 15 | 9 | 7 | 3 | 3 | Ras-related protein Rab-21 |
| Nitab4.5_0001474g0070 | 16 | 49 | 101 | 63 | 37 | 9 | 22 | 25 | |

| | | | | | | | | | |
|---|---|---|---|---|---|---|---|---|---|
| Nitab4.5_0001392g0090 | 7 | 6 | 5 | 12 | 1 | 3 | 2 | 6 | Unknown Protein |
| Nitab4.5_0000258g0290 | 15 | 16 | 9 | 14 | 4 | 6 | 7 | 5 | Glycogen synthase kinase |
| Nitab4.5_0000072g0120 | 2 | 7 | 5 | 4 | 3 | 3 | 1 | 1 | Alpha-L-fucosidase 1 |
| Nitab4.5_0000123g0220 | 44 | 33 | 42 | 122 | 26 | 21 | 25 | 27 | Atcambp25-binding protein OF |
| Nitab4.5_0000416g0140 | 49 | 56 | 44 | 91 | 44 | 21 | 18 | 17 | Chitinase A |
| Nitab4.5_0001014g0010 | 6 | 6 | 5 | 10 | 2 | 3 | 2 | 4 | Cytochrome P450 |
| Nitab4.5_0000200g0140 | 13 | 13 | 12 | 12 | 4 | 7 | 5 | 5 | 2-hydroxyacid dehydrongenase (Fragment) |
| Nitab4.5_0001764g0140 | 19 | 36 | 11 | 20 | 7 | 8 | 2 | 19 | Unknown Protein |
| Nitab4.5_0000825g0110 | 70 | 55 | 57 | 38 | 39 | 18 | 22 | 13 | Receptor serine_threonine kinase |
| Nitab4.5_0002219g0030 | 25 | 19 | 15 | 41 | 14 | 8 | 5 | 14 | Unknown Protein |
| Nitab4.5_0004300g0180 | 4 | 9 | 6 | 5 | 5 | 2 | 1 | 2 | NAC domain transcription factor protein |
| Nitab4.5_0002909g0010 | 66 | 48 | 50 | 27 | 34 | 20 | 16 | 10 | Receptor-like protein kinase |
| Nitab4.5_0000489g0040 | 5 | 3 | 3 | 7 | 1 | 2 | 1 | 3 | Mediator of RNA polymerase II transcription subunit 25 |
| Nitab4.5_0000130g0320 | 25 | 23 | 22 | 13 | 13 | 7 | 4 | 10 | WRKY transcription factor |
| Nitab4.5_0002649g0020 | 20 | 19 | 12 | 25 | 11 | 8 | 8 | 6 | Calcium-dependent protein kinase 4 |
| Nitab4.5_0001133g0040 | 40 | 47 | 33 | 33 | 26 | 7 | 8 | 24 | BURP domain-containing protein |
| Nitab4.5_0000486g0060 | 7 | 15 | 10 | 8 | 3 | 1 | 2 | 11 | Golgi SNAP receptor complex member 1 |
| Nitab4.5_0002192g0050 | 12 | 7 | 3 | 4 | 5 | 2 | 1 | 3 | Serine_threonine-protein kinase receptor |
| Nitab4.5_0001616g0120 | 6 | 5 | 3 | 1 | 2 | 2 | 1 | 2 | Os03g0731050 protein (Fragment) |
| Nitab4.5_0002463g0040 | 3 | 9 | 5 | 3 | 3 | 1 | 3 | 2 | Cathepsin B |
| Nitab4.5_0000280g0250 | 13 | 11 | 12 | 9 | 5 | 6 | 4 | 3 | Cc-nbs-lrr, resistance protein |
| Nitab4.5_0000794g0140 | 29 | 30 | 37 | 28 | 15 | 14 | 11 | 14 | Calmodulin binding protein |
| Nitab4.5_0000064g0010 | 15 | 9 | 8 | 9 | 6 | 5 | 4 | 1 | DMI1 protein (Fragment)-binding domain |
| Nitab4.5_0002238g0030 | 31 | 22 | 16 | 27 | 10 | 12 | 8 | 11 | class I heat shock protein 1 |
| Nitab4.5_0001135g0060 | 88 | 38 | 52 | 37 | 29 | 23 | 20 | 20 | CCR4-NOT transcription complex subunit 7 |
| Nitab4.5_0005003g0010 | 20 | 12 | 24 | 25 | 3 | 15 | 8 | 9 | Mitochondrial import inner membrane translocase subunit tim23 |

| | | | | | | | | | |
|---|---|---|---|---|---|---|---|---|---|
| Nitab4.5_0000588g0160 | 20 | 36 | 26 | 55 | 11 | 21 | 7 | 18 | Os03g0133300 protein (Fragment) |
| Nitab4.5_0003914g0100 | 9 | 6 | 7 | 12 | 6 | 4 | 2 | 3 | AT4G35080-like protein (Fragment) |
| Nitab4.5_0002907g0030 | 2 | 1 | 1 | 2 | 1 | 0 | 1 | 0 | Serine_threonine kinase receptor |
| Nitab4.5_0003965g0070 | 5 | 7 | 6 | 8 | 1 | 2 | 3 | 5 | Nitrilase 1 like protein |
| Nitab4.5_0000208g0400 | 15 | 8 | 10 | 8 | 7 | 4 | 3 | 4 | Receptor-like protein kinase At3g21340 |
| Nitab4.5_0003304g0010 | 8 | 9 | 7 | 10 | 6 | 2 | 3 | 4 | DEP domain-containing protein 1B |
| Nitab4.5_0004554g0020 | 18 | 12 | 10 | 35 | 10 | 5 | 9 | 9 | Exoribonuclease R |
| Nitab4.5_0003180g0060 | 23 | 18 | 16 | 17 | 6 | 3 | 10 | 13 | BHLH transcription factor |
| Nitab4.5_0002765g0010 | 23 | 10 | 13 | 24 | 8 | 4 | 10 | 8 | Caffeoyl-CoA O-methyltransferase |
| Nitab4.5_0000051g0310 | 3 | 6 | 9 | 9 | 3 | 4 | 2 | 3 | Thaumatin-like protein 12104-13574 |
| Nitab4.5_0004152g0010 | 53 | 35 | 13 | 61 | 32 | 11 | 13 | 15 | Unknown Protein |
| Nitab4.5_0003227g0030 | 4 | 6 | 4 | 4 | 1 | 4 | 2 | 2 | Beta-mannosidase |
| Nitab4.5_0002632g0030 | 6 | 19 | 20 | 7 | 9 | 3 | 6 | 4 | Unknown Protein |
| Nitab4.5_0000718g0300 | 4 | 9 | 3 | 9 | 3 | 3 | 2 | 2 | -- |
| Nitab4.5_0000582g0180 | 10 | 19 | 12 | 23 | 7 | 7 | 5 | 8 | Phenylalanine ammonia-lyase |
| Nitab4.5_0000514g0110 | 3 | 4 | 9 | 4 | 2 | 3 | 3 | 1 | Glucose-6P_phosphate translocator |
| Nitab4.5_0002649g0010 | 3 | 2 | 4 | 10 | 2 | 1 | 4 | 2 | Alkaline alpha galactosidase |
| Nitab4.5_0004195g0080 | 58 | 42 | 58 | 28 | 37 | 12 | 16 | 17 | Nodulin-like protein (Fragment) |
| Nitab4.5_0000231g0070 | 5 | 7 | 8 | 4 | 2 | 3 | 2 | 4 | Ribosome maturation factor rimP |
| Nitab4.5_0007385g0010 | 20 | 21 | 11 | 10 | 3 | 10 | 9 | 6 | RING finger protein 38 |
| Nitab4.5_0001315g0010 | 4 | 5 | 1 | 9 | 2 | 2 | 2 | 3 | Peptide transporter 1 |
| Nitab4.5_0000867g0110 | 5 | 4 | 5 | 3 | 2 | 4 | 1 | 1 | -- |
| Nitab4.5_0000368g0350 | 25 | 14 | 12 | 11 | 8 | 7 | 9 | 3 | GRAS family transcription factor |
| Nitab4.5_0007178g0030 | 3 | 6 | 8 | 3 | 3 | 3 | 1 | 1 | Microtubule-associated protein TORTIFOLIA1 |
| Nitab4.5_0000170g0440 | 170 | 133 | 79 | 275 | 65 | 56 | 76 | 95 | Unknown Protein |
| Nitab4.5_0000355g0180 | 3 | 2 | 6 | 8 | 2 | 1 | 3 | 2 | F-box_LRR-repeat protein 3 |

| | | | | | | | | | |
|---|---|---|---|---|---|---|---|---|---|
| Nitab4.5_0000551g0260 | 18 | 13 | 17 | 6 | 9 | 5 | 5 | 5 | -- |
| Nitab4.5_0004646g0060 | 3 | 3 | 5 | 2 | 2 | 1 | 0 | 2 | GRAS family transcription factor |
| Nitab4.5_0001200g0090 | 56 | 58 | 34 | 75 | 28 | 18 | 30 | 24 | Hydroxymethylglutaryl-CoA synthase |
| Nitab4.5_0002761g0020 | 7 | 12 | 8 | 5 | 4 | 4 | 2 | 4 | Solute carrier family 22 member 5 (Predicted) |
| Nitab4.5_0001046g0030 | 10 | 3 | 5 | 10 | 3 | 3 | 4 | 3 | Neurogenic locus notch protein-like |
| Nitab4.5_0000473g0120 | 5 | 8 | 7 | 4 | 4 | 1 | 3 | 3 | Mitochondrial carrier family |
| Nitab4.5_0005528g0030 | 7 | 4 | 3 | 3 | 3 | 2 | 0 | 3 | Beta-lactamase domain protein |
| Nitab4.5_0002229g0170 | 14 | 15 | 13 | 13 | 5 | 7 | 7 | 6 | DDRGK domain-containing protein 1 |
| Nitab4.5_0000075g0120 | 3 | 2 | 2 | 1 | 1 | 2 | 0 | 0 | Cc-nbs-lrr, resistance protein |
| Nitab4.5_0001496g0050 | 37 | 32 | 23 | 29 | 24 | 8 | 10 | 13 | Lysine ketoglutarate reductase trans-splicing related 1 |
| Nitab4.5_0001373g0080 | 11 | 11 | 4 | 8 | 2 | 4 | 5 | 5 | RER1 protein |
| Nitab4.5_0000028g0060 | 17 | 24 | 14 | 7 | 11 | 6 | 3 | 8 | Glycosyl transferase family 17 protein |
| Nitab4.5_0003527g0010 | 7 | 9 | 13 | 8 | 6 | 3 | 3 | 5 | Pterin-4-alpha-carbinolamine dehydratase |
| Nitab4.5_0000354g0130 | 19 | 22 | 14 | 47 | 8 | 6 | 15 | 18 | Hydroxycinnamoyl CoA quinate transferase 2 |
| Nitab4.5_0002003g0070 | 60 | 63 | 44 | 67 | 30 | 23 | 20 | 34 | LOB domain protein 38 |
| Nitab4.5_0003034g0050 | 5 | 3 | 4 | 6 | 2 | 2 | 2 | 2 | Os05g0264200 protein (Fragment) |
| Nitab4.5_0000363g0070 | 10 | 5 | 4 | 12 | 8 | 2 | 2 | 3 | Protein tolB |
| Nitab4.5_0001234g0050 | 21 | 16 | 12 | 16 | 11 | 6 | 6 | 6 | CBL-interacting protein kinase 11 |
| Nitab4.5_0000187g0040 | 17 | 17 | 13 | 47 | 11 | 9 | 12 | 12 | Plant-specific domain TIGR01615 family protein |
| Nitab4.5_0000006g0530 | 7 | 7 | 5 | 10 | 4 | 3 | 3 | 3 | CONSTANS-like zinc finger protein |
| Nitab4.5_0000048g0080 | 43 | 71 | 105 | 25 | 36 | 35 | 31 | 11 | WRKY transcription factor 6 |
| Nitab4.5_0000080g0290 | 10 | 9 | 10 | 11 | 1 | 5 | 4 | 9 | Cysteine-type peptidase |
| Nitab4.5_0001600g0120 | 95 | 112 | 79 | 83 | 59 | 34 | 32 | 47 | LRR receptor-like serine_threonine-protein kinase, RLP |
| Nitab4.5_0000696g0030 | 23 | 16 | 14 | 8 | 13 | 4 | 5 | 6 | WRKY transcription factor |
| Nitab4.5_0000080g0010 | 6 | 9 | 8 | 25 | 4 | 7 | 7 | 4 | Dehydration-responsive family protein |
| Nitab4.5_0000308g0310 | 13 | 8 | 6 | 7 | 7 | 4 | 3 | 2 | Genomic DNA chromosome 5 TAC clone K1F13 |

| | | | | | | | | | |
|---|---|---|---|---|---|---|---|---|---|
| Nitab4.5_0000651g0250 | 8 | 11 | 8 | 16 | 6 | 3 | 4 | 6 | Os07g0587200 protein (Fragment) |
| Nitab4.5_0000171g0310 | 27 | 21 | 21 | 37 | 15 | 11 | 14 | 10 | Unknown Protein |
| Nitab4.5_0000207g0430 | 6 | 6 | 7 | 5 | 4 | 4 | 2 | 2 | Serine_threonine kinase receptor |
| Nitab4.5_0000021g0020 | 76 | 49 | 43 | 134 | 37 | 28 | 35 | 42 | O-methyltransferase |
| Nitab4.5_0003858g0010 | 320 | 303 | 193 | 253 | 170 | 87 | 102 | 147 | Peroxidase |
| Nitab4.5_0000742g0100 | 47 | 46 | 36 | 65 | 24 | 28 | 20 | 19 | Unknown Protein |
| Nitab4.5_0001320g0070 | 3 | 2 | 3 | 2 | 0 | 1 | 2 | 2 | Myosin |
| Nitab4.5_0000377g0160 | 11 | 5 | 8 | 8 | 4 | 4 | 4 | 3 | Unknown Protein |
| Nitab4.5_0000618g0030 | 15 | 7 | 7 | 7 | 9 | 4 | 5 | 1 | Glycosyl transferase family 8 glycogenin |
| Nitab4.5_0001450g0190 | 29 | 21 | 24 | 31 | 12 | 16 | 11 | 11 | U-box domain-containing protein 14 |
| Nitab4.5_0001373g0040 | 7 | 5 | 8 | 6 | 3 | 4 | 1 | 6 | Receptor like kinase, RLK |
| Nitab4.5_0003247g0060 | 12 | 6 | 9 | 8 | 3 | 5 | 4 | 5 | Zinc finger-homeodomain protein 1 (Fragment) |
| Nitab4.5_0005478g0020 | 8 | 8 | 6 | 7 | 6 | 4 | 1 | 2 | Farnesyl pyrophosphate synthase |
| Nitab4.5_0001244g0100 | 4 | 4 | 3 | 1 | 2 | 1 | 2 | 1 | -- |
| Nitab4.5_0002245g0130 | 26 | 24 | 14 | 18 | 9 | 10 | 8 | 12 | Transmembrane protein 222 (Fragment) |
| Nitab4.5_0001558g0060 | 10 | 6 | 10 | 4 | 1 | 2 | 6 | 5 | Oligopeptide transporter 4 |
| Nitab4.5_0000438g0070 | 76 | 52 | 41 | 45 | 30 | 24 | 27 | 23 | Calcium-dependent protein kinase |
| Nitab4.5_0002080g0050 | 28 | 43 | 21 | 50 | 14 | 15 | 13 | 26 | F-box protein PP2-B1 |
| Nitab4.5_0000306g0170 | 231 | 169 | 124 | 156 | 114 | 83 | 72 | 59 | ATP-binding cassette transporter |
| Nitab4.5_0004506g0060 | 7 | 6 | 7 | 5 | 2 | 4 | 3 | 3 | WRKY transcription factor 6 |
| Nitab4.5_0000092g0120 | 12 | 9 | 7 | 12 | 4 | 6 | 4 | 5 | GRAS family transcription factor |
| Nitab4.5_0001175g0140 | 87 | 85 | 65 | 93 | 58 | 32 | 30 | 40 | Binding protein |
| Nitab4.5_0000041g0380 | 13 | 11 | 10 | 8 | 3 | 7 | 3 | 8 | Serine_threonine-protein phosphatase 2A regulatory subunit delta 1 isoform |
| Nitab4.5_0000622g0170 | 127 | 83 | 92 | 139 | 105 | 42 | 40 | 28 | 1-aminocyclopropane-1-carboxylate oxidase |
| Nitab4.5_0000883g0070 | 3 | 3 | 4 | 2 | 3 | 1 | 1 | 1 | Receptor-like kinase |
| Nitab4.5_0001231g0120 | 5 | 4 | 4 | 8 | 4 | 2 | 2 | 2 | RING finger protein |

| | | | | | | | | | |
|---|---|---|---|---|---|---|---|---|---|
| Nitab4.5_0001822g0080 | 9 | 6 | 4 | 5 | 2 | 4 | 2 | 3 | Pentatricopeptide repeat-containing protein-like protein |
| Nitab4.5_0000101g0080 | 24 | 22 | 16 | 13 | 10 | 10 | 8 | 8 | Receptor-like kinase |
| Nitab4.5_0001301g0070 | 126 | 175 | 97 | 233 | 59 | 66 | 81 | 102 | E3 ubiquitin-protein ligase MARCH6 |
| Nitab4.5_0002399g0010 | 19 | 26 | 16 | 26 | 13 | 8 | 12 | 9 | Transmembrane protein 56 |
| Nitab4.5_0004087g0030 | 6 | 7 | 5 | 8 | 4 | 4 | 2 | 3 | Receptor-like kinase |
| Nitab4.5_0000519g0310 | 17 | 11 | 9 | 13 | 8 | 8 | 5 | 3 | Genomic DNA chromosome 3 P1 clone MLD14 |
| Nitab4.5_0000188g0340 | 58 | 57 | 108 | 65 | 44 | 22 | 25 | 51 | Endochitinase (Chitinase) |
| Nitab4.5_0000326g0340 | 5 | 4 | 3 | 8 | 3 | 3 | 2 | 2 | Myosin XI-2 |
| Nitab4.5_0000005g0190 | 6 | 5 | 3 | 3 | 1 | 4 | 2 | 2 | Transcription factor WRKY |
| Nitab4.5_0000052g0220 | 67 | 67 | 56 | 172 | 56 | 37 | 44 | 42 | Aspartate aminotransferase |
| Nitab4.5_0000033g0230 | 19 | 20 | 21 | 30 | 19 | 8 | 9 | 8 | CHP-rich zinc finger protein-like |
| Nitab4.5_0007482g0040 | 23 | 13 | 19 | 17 | 3 | 9 | 12 | 12 | Mediator of RNA polymerase II transcription subunit 21 |
| Nitab4.5_0001288g0100 | 3 | 4 | 7 | 4 | 5 | 1 | 2 | 1 | Receptor like kinase, RLK |
| Nitab4.5_0000676g0060 | 1 | 6 | 11 | 6 | 3 | 2 | 3 | 5 | Serine_threonine-protein phosphatase 6 regulatory ankyrin repeat subunit A |
| Nitab4.5_0000721g0200 | 17 | 21 | 18 | 13 | 14 | 6 | 7 | 7 | Calcium dependent protein kinase 1 |
| Nitab4.5_0000297g0160 | 9 | 7 | 11 | 27 | 5 | 7 | 5 | 9 | Uncharacterized secreted protein |
| Nitab4.5_0003316g0030 | 12 | 23 | 12 | 23 | 11 | 6 | 10 | 8 | WRKY transcription factor 2 |
| Nitab4.5_0002320g0020 | 34 | 46 | 44 | 45 | 32 | 21 | 15 | 16 | Hydroxycinnamoyl CoA shikimate_quinate hydroxycinnamoyltransferase |
| Nitab4.5_0000055g0100 | 11 | 13 | 8 | 35 | 11 | 6 | 8 | 9 | Saccharopine dehydrogenase (NAD(+) L-glutamate-forming) |
| Nitab4.5_0001317g0070 | 186 | 130 | 112 | 202 | 88 | 75 | 73 | 79 | Arginine decarboxylase |
| Nitab4.5_0001713g0020 | 18 | 12 | 22 | 12 | 40 | 39 | 24 | 28 | Os12g0604200 protein (Fragment) |
| Nitab4.5_0001913g0140 | 2 | 4 | 4 | 3 | 9 | 6 | 6 | 3 | Genomic DNA chromosome 3 TAC clone K10D20 |
| Nitab4.5_0003477g0040 | 11 | 13 | 13 | 10 | 34 | 27 | 21 | 12 | Transcription factor |
| Nitab4.5_0001735g0040 | 2 | 3 | 3 | 2 | 2 | 4 | 10 | 5 | Glutamate decarboxylase |
| Nitab4.5_0003746g0030 | 20 | 21 | 24 | 24 | 42 | 62 | 43 | 31 | RAG1-activating protein 1 homolog |
| Nitab4.5_0003855g0010 | 2 | 2 | 2 | 2 | 3 | 4 | 2 | 5 | Serine_threonine protein kinase-like |

| | | | | | | | | | |
|---|---|---|---|---|---|---|---|---|---|
| Nitab4.5_0002403g0050 | 3 | 6 | 4 | 14 | 12 | 10 | 18 | 14 | Fasciclin-like arabinogalactan protein 9 |
| Nitab4.5_0000603g0070 | 5 | 3 | 6 | 7 | 15 | 8 | 12 | 7 | Copper transport protein 86 |
| Nitab4.5_0000104g0010 | 10 | 11 | 9 | 5 | 19 | 20 | 20 | 14 | Phototropic-responsive NPH3 family protein |
| Nitab4.5_0004599g0090 | 2 | 11 | 10 | 5 | 9 | 18 | 17 | 12 | BZIP transcription factor family protein |
| Nitab4.5_0001586g0100 | 24 | 21 | 25 | 12 | 46 | 42 | 39 | 36 | Chitinase |
| Nitab4.5_0000037g0170 | 3 | 2 | 5 | 3 | 4 | 4 | 6 | 11 | Unknown Protein |
| Nitab4.5_0001044g0050 | 1 | 3 | 3 | 1 | 4 | 6 | 3 | 5 | Leucine Rich Repeat family protein expressed |
| Nitab4.5_0000895g0210 | 8 | 6 | 8 | 9 | 10 | 16 | 25 | 11 | Pentatricopeptide repeat-containing protein |
| Nitab4.5_0001823g0020 | 13 | 14 | 20 | 11 | 28 | 24 | 31 | 33 | Malate dehydrogenase, cytoplasmic |
| Nitab4.5_0000107g0120 | 25 | 16 | 30 | 17 | 31 | 73 | 55 | 19 | O-methyltransferase |
| Nitab4.5_0001560g0050 | 41 | 30 | 32 | 34 | 91 | 83 | 67 | 36 | Serine_threonine protein kinase |
| Nitab4.5_0000132g0510 | 2 | 2 | 3 | 5 | 4 | 7 | 7 | 8 | Ankyrin repeat domain 1 |
| Nitab4.5_0000203g0030 | 26 | 8 | 10 | 13 | 25 | 36 | 37 | 19 | BURP domain-containing protein (Fragment) |
| Nitab4.5_0000046g0200 | 502 | 564 | 874 | 481 | 1278 | 1545 | 1249 | 865 | Non-specific lipid-transfer protein |
| Nitab4.5_0001120g0010 | 2 | 1 | 3 | 1 | 4 | 5 | 4 | 1 | Multidrug resistance protein ABC transporter family |
| Nitab4.5_0002632g0060 | 2 | 3 | 1 | 1 | 3 | 3 | 4 | 4 | Phosphoglycerate kinase |
| Nitab4.5_0000173g0030 | 6 | 7 | 8 | 16 | 10 | 29 | 20 | 18 | Aquaporin-like protein |
| Nitab4.5_0000363g0310 | 7 | 8 | 11 | 4 | 23 | 15 | 15 | 8 | DNAJ heat shock N-terminal domain-containing protein |
| Nitab4.5_0000004g0060 | 3 | 4 | 3 | 2 | 4 | 4 | 9 | 8 | mRNA binding protein Pumilio 2 |
| Nitab4.5_0000080g0080 | 11 | 7 | 8 | 9 | 21 | 28 | 18 | 5 | Long-chain-fatty-acid--CoA ligase |
| Nitab4.5_0002552g0010 | 19 | 10 | 15 | 14 | 33 | 39 | 21 | 24 | Lipase |
| Nitab4.5_0000831g0030 | 7 | 23 | 16 | 9 | 23 | 35 | 30 | 26 | BZIP transcription factor family protein |
| Nitab4.5_0002085g0130 | 10 | 16 | 19 | 9 | 30 | 24 | 28 | 29 | GDSL esterase_lipase At5g33370 |
| Nitab4.5_0001797g0030 | 3 | 3 | 3 | 2 | 4 | 8 | 5 | 6 | Glycosyltransferase |
| Nitab4.5_0000048g0070 | 5 | 4 | 8 | 7 | 8 | 8 | 17 | 17 | GDSL esterase_lipase At1g28590 |
| Nitab4.5_0000633g0070 | 2 | 3 | 4 | 8 | 8 | 12 | 7 | 9 | AT-hook motif nuclear localized protein 1 |

| | | | | | | | | | |
|---|---|---|---|---|---|---|---|---|---|
| Nitab4.5_0002482g0070 | 53 | 53 | 58 | 23 | 78 | 137 | 107 | 66 | Polyphenol oxidase |
| Nitab4.5_0000202g0470 | 30 | 29 | 24 | 15 | 43 | 60 | 50 | 55 | Glutathione S-transferase |
| Nitab4.5_0002393g0070 | 9 | 3 | 8 | 8 | 11 | 11 | 24 | 12 | 50S ribosomal protein L28 |
| Nitab4.5_0004979g0130 | 9 | 6 | 7 | 4 | 15 | 11 | 13 | 14 | FIP1 |
| Nitab4.5_0001082g0080 | 10 | 4 | 6 | 7 | 8 | 20 | 18 | 11 | Neutral invertase like protein |
| Nitab4.5_0005622g0060 | 3 | 1 | 2 | 2 | 4 | 1 | 5 | 4 | Proline-, glutamic acid- and leucine-rich protein 1 |
| Nitab4.5_0000391g0390 | 2 | 1 | 1 | 2 | 4 | 5 | 3 | 2 | Xenotropic and polytropic retrovirus receptor |
| Nitab4.5_0005169g0010 | 4 | 1 | 3 | 2 | 6 | 6 | 8 | 2 | Octicosapeptide_Phox_Bem1p domain-containing protein |
| Nitab4.5_0002165g0040 | 4 | 2 | 6 | 4 | 5 | 11 | 8 | 10 | Replication protein A subunit |
| Nitab4.5_0000083g0140 | 3 | 4 | 4 | 2 | 10 | 4 | 7 | 7 | Pentatricopeptide repeat-containing protein At4g21190 |
| Nitab4.5_0006360g0010 | 7 | 14 | 13 | 18 | 33 | 25 | 32 | 19 | Heat stress transcription factor A3-type, DNA-binding |
| Nitab4.5_0001231g0030 | 2 | 2 | 3 | 1 | 4 | 2 | 4 | 8 | DNA primase |
| Nitab4.5_0000243g0070 | 2 | 4 | 4 | 4 | 7 | 9 | 7 | 7 | Glucosyltransferase-2 |
| Nitab4.5_0003649g0020 | 7 | 14 | 13 | 11 | 38 | 23 | 15 | 19 | Outer membrane lipoprotein blc |
| Nitab4.5_0000976g0170 | 4 | 5 | 14 | 5 | 17 | 16 | 14 | 13 | Sec14-like (Fragment) |
| Nitab4.5_0001034g0170 | 8 | 6 | 3 | 2 | 11 | 13 | 8 | 11 | ATP-dependent RNA helicase fal1 |
| Nitab4.5_0002927g0060 | 2 | 3 | 3 | 2 | 5 | 5 | 4 | 6 | Exosome complex exonuclease RRP4 |
| Nitab4.5_0000178g0330 | 4 | 1 | 3 | 6 | 6 | 8 | 10 | 7 | Mannan endo-1 4-beta-mannosidase |
| Nitab4.5_0001970g0020 | 12 | 12 | 28 | 17 | 20 | 19 | 38 | 68 | Calcium_proton exchanger |
| Nitab4.5_0003559g0050 | 5 | 9 | 12 | 7 | 16 | 21 | 19 | 14 | Unknown Protein |
| Nitab4.5_0002265g0180 | 2 | 5 | 4 | 3 | 8 | 7 | 11 | 5 | Beta-1,3-galactosyl-O-glycosyl-glycoprotein beta-1,6-N-acetylglucosaminyltransferase 4 |
| Nitab4.5_0003077g0030 | 4 | 5 | 2 | 3 | 5 | 5 | 13 | 10 | Unknown Protein |
| Nitab4.5_0000516g0020 | 1 | 2 | 2 | 1 | 4 | 2 | 4 | 3 | Unknown Protein |
| Nitab4.5_0000614g0050 | 21 | 8 | 5 | 13 | 24 | 41 | 25 | 13 | Unknown Protein |
| Nitab4.5_0003978g0050 | 1 | 6 | 6 | 4 | 7 | 7 | 15 | 7 | Ganglioside-induced differentiation-associated protein 1 |

| | | | | | | | | | |
|---|---|---|---|---|---|---|---|---|---|
| Nitab4.5_0001728g0060 | 4 | 1 | 2 | 1 | 5 | 2 | 5 | 4 | Ribonucleoside-diphosphate reductase |
| Nitab4.5_0001989g0070 | 4 | 13 | 7 | 5 | 9 | 15 | 12 | 28 | 3-oxo-5-alpha-steroid 4-dehydrogenase family protein |
| Nitab4.5_0000094g0020 | 42 | 26 | 31 | 20 | 75 | 71 | 66 | 50 | Hydrolase alpha_beta fold family protein expressed |
| Nitab4.5_0000855g0100 | 2 | 1 | 2 | 2 | 7 | 2 | 3 | 4 | MORC family CW-type zinc finger 3 |
| Nitab4.5_0003062g0020 | 2 | 1 | 1 | 3 | 6 | 3 | 4 | 3 | Cc-nbs-lrr, resistance protein with an R1 specific domain |
| Nitab4.5_0002616g0040 | 3 | 4 | 3 | 2 | 5 | 7 | 8 | 7 | Unknown Protein |
| Nitab4.5_0001268g0020 | 9 | 13 | 10 | 20 | 6 | 22 | 27 | 59 | Calcium-dependent protein kinase 2 |
| Nitab4.5_0001256g0060 | 5 | 1 | 4 | 2 | 9 | 8 | 6 | 3 | Malonyl CoA anthocyanin 3-O-glucoside-6_apos_apos-O-malonyltransferase |
| Nitab4.5_0004813g0050 | 9 | 9 | 7 | 12 | 19 | 30 | 22 | 11 | WD-40 repeat family protein |
| Nitab4.5_0001494g0050 | 13 | 13 | 15 | 6 | 22 | 38 | 24 | 18 | Transcription factor |
| Nitab4.5_0002252g0080 | 2 | 1 | 1 | 2 | 3 | 3 | 3 | 5 | DnaJ homolog subfamily C member 7 |
| Nitab4.5_0000129g0190 | 1 | 2 | 2 | 0 | 2 | 2 | 3 | 3 | WD-repeat protein |
| Nitab4.5_0001163g0010 | 8 | 7 | 12 | 9 | 12 | 19 | 17 | 32 | Self-pruning interacting protein 1 |
| Nitab4.5_0000154g0280 | 3 | 5 | 5 | 4 | 4 | 13 | 14 | 7 | Hydrolase alpha_beta fold family protein |
| Nitab4.5_0002762g0010 | 11 | 18 | 7 | 15 | 23 | 24 | 30 | 36 | Beta-xylosidase 1 |
| Nitab4.5_0000187g0160 | 30 | 30 | 13 | 15 | 68 | 49 | 37 | 39 | Auxin responsive protein |
| Nitab4.5_0001892g0050 | 7 | 5 | 4 | 4 | 13 | 17 | 9 | 4 | Multi-sensor hybrid histidine kinase |
| Nitab4.5_0000287g0320 | 7 | 9 | 4 | 7 | 15 | 12 | 18 | 15 | Peroxisomal multifunctional enzyme type 2 |
| Nitab4.5_0000928g0090 | 20 | 20 | 8 | 20 | 33 | 39 | 44 | 35 | Amino acid transporter |
| Nitab4.5_0001030g0140 | 6 | 11 | 8 | 6 | 17 | 22 | 15 | 14 | Glycosyltransferase family GT8 protein |
| Nitab4.5_0004651g0010 | 33 | 25 | 23 | 28 | 74 | 56 | 72 | 42 | Unknown Protein |
| Nitab4.5_0000307g0240 | 17 | 31 | 47 | 13 | 65 | 78 | 69 | 34 | BHLH transcription factor |
| Nitab4.5_0003381g0060 | 2 | 4 | 5 | 2 | 7 | 10 | 4 | 8 | Iaa-amino acid hydrolase 9 |
| Nitab4.5_0000567g0070 | 2 | 4 | 2 | 2 | 4 | 2 | 8 | 7 | Serine carboxypeptidase |
| Nitab4.5_0000396g0260 | 2 | 2 | 1 | 1 | 3 | 3 | 3 | 4 | Glycogen synthase |
| Nitab4.5_0002421g0040 | 7 | 3 | 7 | 7 | 13 | 8 | 16 | 14 | Pentatricopeptide repeat-containing protein |

| | | | | | | | | | |
|---|---|---|---|---|---|---|---|---|---|
| Nitab4.5_0005376g0030 | 1 | 2 | 3 | 3 | 4 | 6 | 6 | 3 | Pentatricopeptide repeat-containing protein |
| Nitab4.5_0001934g0020 | 2 | 5 | 4 | 2 | 14 | 3 | 3 | 7 | Cytochrome P450 |
| Nitab4.5_0000218g0060 | 3 | 4 | 3 | 1 | 5 | 12 | 6 | 1 | Acyltransferase-like protein |
| Nitab4.5_0007418g0010 | 3 | 4 | 12 | 7 | 15 | 16 | 8 | 22 | -- |
| Nitab4.5_0004787g0010 | 65 | 52 | 45 | 34 | 91 | 155 | 136 | 65 | Unknown Protein |
| Nitab4.5_0000635g0100 | 156 | 153 | 215 | 66 | 307 | 486 | 352 | 202 | Auxin-responsive GH3-like |
| Nitab4.5_0000603g0060 | 9 | 4 | 4 | 14 | 13 | 24 | 15 | 19 | Os12g0236050 protein (Fragment) |
| Nitab4.5_0001356g0110 | 6 | 14 | 5 | 9 | 17 | 31 | 13 | 17 | GTP binding protein |
| Nitab4.5_0002577g0050 | 3 | 3 | 4 | 2 | 6 | 4 | 9 | 6 | Ribonucleoside-diphosphate reductase |
| Nitab4.5_0000085g0320 | 4 | 3 | 3 | 2 | 6 | 6 | 6 | 10 | Homocysteine s-methyltransferase |
| Nitab4.5_0000353g0010 | 1 | 1 | 2 | 5 | 5 | 3 | 12 | 4 | Heat stress transcription factor A3-type, DNA-binding |
| Nitab4.5_0000194g0290 | 3 | 7 | 5 | 5 | 18 | 15 | 11 | 4 | Unknown Protein |
| Nitab4.5_0002070g0010 | 150 | 145 | 105 | 68 | 345 | 327 | 284 | 125 | -- |
| Nitab4.5_0000785g0140 | 2 | 5 | 1 | 1 | 3 | 6 | 7 | 6 | Genomic DNA chromosome 5 TAC clone K19B1 |
| Nitab4.5_0000410g0370 | 15 | 5 | 7 | 7 | 17 | 28 | 23 | 12 | Hydrolase alpha_beta fold family protein |
| Nitab4.5_0001701g0170 | 36 | 39 | 37 | 40 | 61 | 86 | 99 | 108 | Fasciclin-like arabinogalactan protein 19 |
| Nitab4.5_0002352g0050 | 2 | 5 | 4 | 2 | 6 | 6 | 11 | 5 | Phytochrome kinase substrate 1 |
| Nitab4.5_0000315g0010 | 2 | 8 | 6 | 4 | 10 | 9 | 15 | 13 | Unknown Protein |
| Nitab4.5_0000081g0120 | 5 | 3 | 3 | 6 | 8 | 6 | 10 | 16 | Germin-like protein |
| Nitab4.5_0000073g0350 | 12 | 4 | 5 | 7 | 22 | 19 | 17 | 7 | class I heat shock protein |
| Nitab4.5_0001556g0100 | 24 | 21 | 12 | 17 | 57 | 43 | 39 | 36 | Auxin responsive protein |
| Nitab4.5_0003138g0030 | 0 | 1 | 1 | 1 | 2 | 1 | 3 | 1 | Receptor like kinase, RLK |
| Nitab4.5_0002649g0040 | 2 | 5 | 4 | 3 | 8 | 7 | 8 | 9 | Histone-lysine N-methyltransferase NSD3 |
| Nitab4.5_0000008g0230 | 21 | 10 | 19 | 29 | 46 | 78 | 42 | 20 | Unknown Protein |
| Nitab4.5_0000246g0010 | 2 | 3 | 1 | 3 | 7 | 6 | 5 | 5 | E3 SUMO-protein ligase NSE2 |
| Nitab4.5_0000956g0070 | 6 | 9 | 10 | 9 | 10 | 17 | 26 | 28 | Phosphatidylinositol-specific phospholipase c |

| | | | | | | | | | |
|---|---|---|---|---|---|---|---|---|---|
| Nitab4.5_0007026g0060 | 14 | 4 | 9 | 14 | 40 | 28 | 19 | 9 | Inositol 1 4 5-trisphosphate 5-phosphatase |
| Nitab4.5_0000104g0330 | 2 | 1 | 7 | 4 | 8 | 9 | 7 | 8 | Genomic DNA chromosome 5 P1 clone MQN23 |
| Nitab4.5_0000679g0110 | 1 | 1 | 3 | 1 | 2 | 5 | 4 | 2 | 5_apos-AMP-activated protein kinase subunit beta-2 |
| Nitab4.5_0000102g0020 | 14 | 8 | 1 | 3 | 20 | 20 | 18 | 5 | Os06g0207500 protein (Fragment) |
| Nitab4.5_0003259g0080 | 144 | 170 | 125 | 203 | 283 | 331 | 425 | 510 | ChaC cation transport regulator-like 1 |
| Nitab4.5_0000257g0230 | 1 | 3 | 2 | 1 | 6 | 6 | 2 | 4 | Polymerase (DNA directed) mu |
| Nitab4.5_0004230g0010 | 35 | 25 | 37 | 31 | 121 | 111 | 61 | 20 | Unknown Protein |
| Nitab4.5_0000770g0100 | 1 | 2 | 1 | 2 | 1 | 3 | 5 | 6 | Laccase |
| Nitab4.5_0000006g0100 | 4 | 1 | 5 | 9 | 11 | 15 | 14 | 7 | Glucose-6-phosphate_phosphate translocator 2 |
| Nitab4.5_0003039g0040 | 11 | 1 | 3 | 9 | 17 | 22 | 14 | 5 | Ethylene-responsive transcription factor 13 |
| Nitab4.5_0000709g0070 | 24 | 29 | 26 | 33 | 76 | 94 | 61 | 40 | ACT domain-containing protein |
| Nitab4.5_0000283g0170 | 93 | 62 | 45 | 62 | 280 | 114 | 126 | 125 | Gibberellin-regulated protein 2 |
| Nitab4.5_0000246g0100 | 2 | 7 | 8 | 4 | 8 | 19 | 15 | 9 | BZIP transcription factor family protein |
| Nitab4.5_0000209g0090 | 3 | 3 | 13 | 8 | 10 | 22 | 19 | 18 | Ethylene-responsive transcription factor 11 |
| Nitab4.5_0000401g0150 | 1 | 2 | 3 | 1 | 2 | 3 | 4 | 8 | Methyltransferase |
| Nitab4.5_0004924g0030 | 36 | 12 | 19 | 47 | 76 | 89 | 66 | 50 | Calmodulin-binding protein |
| Nitab4.5_0000123g0610 | 1 | 1 | 1 | 1 | 1 | 2 | 6 | 2 | -- |
| Nitab4.5_0004160g0020 | 1 | 1 | 1 | 2 | 4 | 2 | 3 | 3 | BEL1-like homeodomain protein 3 |
| Nitab4.5_0000682g0080 | 3 | 2 | 3 | 2 | 5 | 11 | 3 | 5 | Peroxidase |
| Nitab4.5_0000091g0510 | 12 | 19 | 13 | 24 | 36 | 62 | 44 | 31 | Xyloglucan endotransglucosylase_hydrolase 1 |
| Nitab4.5_0004425g0020 | 2 | 1 | 2 | 3 | 4 | 3 | 7 | 5 | Cytochrome P450 |
| Nitab4.5_0005206g0010 | 12 | 7 | 7 | 10 | 28 | 37 | 22 | 5 | Cellulose synthase-like C1-2 glycosyltransferase family 2 protein |
| Nitab4.5_0001952g0250 | 11 | 7 | 4 | 7 | 12 | 37 | 17 | 7 | Peroxidase 65 |
| Nitab4.5_0000630g0080 | 17 | 8 | 11 | 6 | 18 | 23 | 40 | 26 | Unknown Protein |
| Nitab4.5_0001492g0030 | 4 | 1 | 2 | 3 | 5 | 5 | 7 | 7 | Caffeoyl-CoA O-methyltransferase |
| Nitab4.5_0001657g0020 | 2 | 1 | 2 | 2 | 4 | 4 | 3 | 6 | Receptor like kinase, RLK |

| | | | | | | | | |
|---|---|---|---|---|---|---|---|---|
| Nitab4.5_0000647g0120 | 3 | 5 | 2 | 2 | 7 | 11 | 6 | 7 | Receptor like protein kinase |
| Nitab4.5_0000978g0100 | 4 | 3 | 4 | 2 | 4 | 12 | 8 | 7 | Chitinase-like protein |
| Nitab4.5_0002700g0020 | 2 | 3 | 2 | 10 | 18 | 7 | 11 | 8 | Unknown Protein |
| Nitab4.5_0000038g0160 | 17 | 10 | 8 | 7 | 32 | 38 | 27 | 12 | Bile acid sodium symporter family protein |
| Nitab4.5_0000565g0270 | 2 | 1 | 1 | 0 | 2 | 3 | 3 | 2 | UDP-glucuronosyltransferase |
| Nitab4.5_0002415g0020 | 17 | 13 | 22 | 55 | 92 | 88 | 67 | 27 | Unknown Protein |
| Nitab4.5_0001612g0090 | 1 | 1 | 1 | 1 | 1 | 5 | 2 | 4 | Zinc ion binding protein |
| Nitab4.5_0000764g0020 | 4 | 1 | 1 | 3 | 6 | 9 | 6 | 4 | Inositol 1 4 5-trisphosphate 5-phosphatase |
| Nitab4.5_0003328g0020 | 47 | 38 | 32 | 59 | 103 | 135 | 129 | 91 | Fasciclin-like arabinogalactan protein 10 |
| Nitab4.5_0004557g0020 | 8 | 10 | 7 | 18 | 31 | 38 | 32 | 13 | Tyrosine-protein kinase transforming protein Src |
| Nitab4.5_0000134g0030 | 19 | 21 | 29 | 16 | 93 | 50 | 30 | 50 | photosystem II polypeptide |
| Nitab4.5_0000256g0410 | 2 | 2 | 1 | 2 | 6 | 4 | 5 | 3 | AT-hook motif nuclear localized protein 13 |
| Nitab4.5_0000040g0410 | 2 | 4 | 2 | 10 | 17 | 11 | 6 | 13 | Unknown Protein |
| Nitab4.5_0000047g0180 | 2 | 1 | 1 | 1 | 3 | 2 | 5 | 3 | Phosphatidylinositol transfer protein SFH5 |
| Nitab4.5_0002315g0170 | 2 | 9 | 8 | 5 | 12 | 23 | 11 | 17 | DNA-3-methyladenine glycosylase I |
| Nitab4.5_0000568g0030 | 7 | 2 | 13 | 8 | 8 | 32 | 21 | 17 | Expansin |
| Nitab4.5_0002654g0010 | 3 | 4 | 3 | 2 | 8 | 7 | 12 | 6 | SNARE associated Golgi protein |
| Nitab4.5_0000124g0040 | 1 | 2 | 4 | 3 | 8 | 7 | 7 | 5 | Gibberellin 20-oxidase-1 |
| Nitab4.5_0000170g0250 | 5 | 3 | 3 | 6 | 10 | 22 | 12 | 4 | LRR receptor-like serine_threonine-protein kinase, RLP |
| Nitab4.5_0000578g0100 | 2 | 3 | 3 | 2 | 7 | 4 | 10 | 4 | Pentatricopeptide repeat-containing protein |
| Nitab4.5_0000036g0030 | 2 | 2 | 2 | 0 | 4 | 8 | 3 | 3 | Speckle-type POZ protein |
| Nitab4.5_0000630g0170 | 6 | 3 | 2 | 10 | 12 | 18 | 16 | 11 | Unknown Protein |
| Nitab4.5_0001196g0050 | 3 | 7 | 7 | 9 | 15 | 9 | 19 | 26 | -- |
| Nitab4.5_0007026g0020 | 5 | 2 | 4 | 1 | 13 | 10 | 6 | 2 | Peptide transporter-like protein |
| Nitab4.5_0002340g0070 | 9 | 3 | 4 | 10 | 16 | 27 | 16 | 10 | C4-dicarboxylate transporter_malic acid transport family protein |
| Nitab4.5_0000302g0040 | 0 | 2 | 4 | 1 | 5 | 3 | 4 | 7 | Syntaxin |

| | | | | | | | | | |
|---|---|---|---|---|---|---|---|---|---|
| Nitab4.5_0003264g0030 | 15 | 7 | 16 | 21 | 46 | 65 | 41 | 7 | Unknown Protein |
| Nitab4.5_0000969g0040 | 2 | 1 | 3 | 0 | 5 | 8 | 2 | 2 | U-box domain-containing protein 13 |
| Nitab4.5_0001591g0040 | 7 | 3 | 4 | 20 | 16 | 25 | 23 | 27 | Blue copper protein |
| Nitab4.5_0002816g0090 | 2 | 7 | 1 | 3 | 9 | 11 | 8 | 6 | Unknown Protein |
| Nitab4.5_0001968g0130 | 2 | 0 | 1 | 2 | 3 | 7 | 2 | 1 | Solute carrier family 15 member 4 |
| Nitab4.5_0000866g0020 | 2 | 14 | 1 | 8 | 16 | 15 | 16 | 21 | Male sterility 5 family protein (Fragment) |
| Nitab4.5_0000568g0130 | 6 | 7 | 5 | 6 | 22 | 32 | 9 | 5 | Xenotropic and polytropic retrovirus receptor |
| Nitab4.5_0000375g0160 | 1 | 1 | 1 | 2 | 2 | 3 | 4 | 6 | CBL-interacting protein kinase 13 |
| Nitab4.5_0001952g0140 | 7 | 5 | 7 | 10 | 19 | 35 | 20 | 6 | Cellulose synthase-like C1-2 glycosyltransferase family 2 protein |
| Nitab4.5_0002314g0090 | 3 | 4 | 5 | 1 | 11 | 11 | 8 | 6 | U-box domain-containing protein 4 |
| Nitab4.5_0000208g0100 | 4 | 3 | 6 | 6 | 12 | 17 | 17 | 6 | Serine_threonine-protein kinase bud32 (EC 2.7.11.1) |
| Nitab4.5_0004588g0010 | 3 | 2 | 2 | 3 | 2 | 4 | 13 | 6 | Os03g0366700 protein (Fragment) |
| Nitab4.5_0004399g0080 | 4 | 2 | 4 | 5 | 7 | 7 | 10 | 18 | Unknown Protein |
| Nitab4.5_0001748g0050 | 10 | 7 | 5 | 9 | 11 | 31 | 27 | 15 | C4-dicarboxylate transporter_malic acid transport family protein |
| Nitab4.5_0001438g0070 | 3 | 3 | 5 | 3 | 16 | 7 | 8 | 5 | Auxin-responsive protein |
| Nitab4.5_0007027g0010 | 2 | 2 | 1 | 1 | 6 | 5 | 5 | 2 | Protein kinase 5 |
| Nitab4.5_0001820g0070 | 1 | 1 | 3 | 2 | 7 | 4 | 4 | 3 | Serine carboxypeptidase K10B2.2 |
| Nitab4.5_0000080g0250 | 40 | 35 | 23 | 38 | 114 | 104 | 82 | 79 | ATP binding _ serine-threonine kinase |
| Nitab4.5_0000804g0070 | 5 | 1 | 4 | 4 | 8 | 13 | 6 | 11 | Aluminum-activated malate transporter (Fragment) |
| Nitab4.5_0001693g0130 | 2 | 1 | 1 | 2 | 3 | 5 | 6 | 4 | Aldo_keto reductase family protein |
| Nitab4.5_0001315g0320 | 2 | 2 | 1 | 0 | 4 | 4 | 6 | 1 | Os01g0786800 protein (Fragment) |
| Nitab4.5_0000116g0440 | 0 | 3 | 2 | 2 | 2 | 2 | 8 | 7 | Phosphoserine phosphatase |
| Nitab4.5_0002031g0120 | 1 | 2 | 2 | 2 | 6 | 10 | 3 | 1 | Receptor protein kinase-like protein |
| Nitab4.5_0001439g0100 | 78 | 48 | 45 | 157 | 164 | 348 | 270 | 143 | Arabinogalactan |
| Nitab4.5_0004554g0010 | 2 | 1 | 0 | 2 | 9 | 4 | 2 | 1 | Xenotropic and polytropic retrovirus receptor |
| Nitab4.5_0000136g0450 | 1 | 2 | 1 | 1 | 3 | 3 | 4 | 4 | Chromodomain-helicase-DNA-binding protein 6 |

| | | | | | | | | | |
|---|---|---|---|---|---|---|---|---|---|
| Nitab4.5_0003603g0040 | 175 | 52 | 20 | 146 | 352 | 419 | 275 | 80 | Expansin-like protein |
| Nitab4.5_0001423g0100 | 8 | 17 | 18 | 9 | 27 | 31 | 46 | 46 | Glucose transporter 8 |
| Nitab4.5_0001049g0080 | 1 | 2 | 1 | 1 | 5 | 1 | 4 | 3 | Survival motor neuron containing protein |
| Nitab4.5_0000509g0080 | 3 | 1 | 1 | 2 | 5 | 6 | 9 | 3 | Unknown protein DS12 from 2D-PAGE of leaf, chloroplastic |
| Nitab4.5_0002629g0030 | 112 | 81 | 42 | 211 | 280 | 501 | 309 | 221 | Fasciclin-like arabinogalactan protein 4 |
| Nitab4.5_0000535g0030 | 3 | 2 | 7 | 5 | 24 | 9 | 9 | 12 | Aluminum-activated malate transporter (Fragment) |
| Nitab4.5_0000257g0150 | 1 | 1 | 0 | 1 | 2 | 6 | 2 | 2 | IAA-amino acid hydrolase |
| Nitab4.5_0002859g0150 | 2 | 5 | 8 | 3 | 18 | 11 | 7 | 14 | D-isomer specific 2-hydroxyacid dehydrogenase |
| Nitab4.5_0001361g0130 | 1 | 1 | 2 | 1 | 1 | 6 | 4 | 2 | Ternary complex factor MIP1 |
| Nitab4.5_0000307g0100 | 2 | 0 | 1 | 1 | 6 | 4 | 1 | 3 | Pentatricopeptide repeat-containing protein |
| Nitab4.5_0000357g0250 | 2 | 2 | 1 | 0 | 2 | 3 | 4 | 4 | Exocyst complex component EXO70 |
| Nitab4.5_0000386g0240 | 5 | 1 | 7 | 8 | 15 | 27 | 15 | 7 | Digalactosyldiacylglycerol synthase 2, chloroplastic |
| Nitab4.5_0000157g0160 | 1 | 1 | 1 | 0 | 1 | 2 | 4 | 3 | 60S ribosomal protein L5, mitochondrial |
| Nitab4.5_0001789g0130 | 3 | 3 | 4 | 4 | 14 | 17 | 7 | 6 | Protein kinase domain containing protein |
| Nitab4.5_0002613g0030 | 27 | 25 | 16 | 33 | 77 | 118 | 76 | 33 | Xyloglucan endotransglucosylase_hydrolase 1 |
| Nitab4.5_0000167g0070 | 2 | 2 | 2 | 3 | 7 | 0 | 10 | 8 | RING finger protein 13 |
| Nitab4.5_0000306g0030 | 5 | 1 | 3 | 4 | 15 | 11 | 6 | 5 | Male sterility MS5 family protein |
| Nitab4.5_0001700g0060 | 22 | 11 | 10 | 29 | 60 | 87 | 55 | 14 | Expressed protein (Fragment) |
| Nitab4.5_0000678g0050 | 1 | 0 | 1 | 1 | 4 | 2 | 1 | 2 | Sulfate transporter |
| Nitab4.5_0003625g0030 | 0 | 7 | 7 | 1 | 15 | 4 | 11 | 17 | -- |
| Nitab4.5_0002356g0060 | 7 | 2 | 7 | 7 | 20 | 30 | 15 | 8 | Hydroxycinnamoyl CoA quinate transferase |
| Nitab4.5_0000137g0100 | 1 | 2 | 2 | 3 | 8 | 4 | 5 | 4 | Lipoxygenase |
| Nitab4.5_0000082g0130 | 1 | 2 | 2 | 3 | 6 | 4 | 3 | 10 | tRNA-splicing endonuclease subunit sen54 |
| Nitab4.5_0002234g0100 | 19 | 24 | 5 | 24 | 72 | 68 | 55 | 22 | BCL-2 binding anthanogene-1 |
| Nitab4.5_0000302g0200 | 2 | 0 | 1 | 4 | 6 | 6 | 4 | 4 | Myb-related transcription factor |
| Nitab4.5_0000610g0050 | 3 | 0 | 2 | 2 | 7 | 1 | 12 | 3 | Genomic DNA chromosome 5 P1 clone MMN10 |

| | | | | | | | | | |
|---|---|---|---|---|---|---|---|---|---|
| Nitab4.5_0001685g0060 | 1 | 4 | 1 | 0 | 5 | 3 | 7 | 5 | F-box family protein |
| Nitab4.5_0002579g0060 | 25 | 4 | 8 | 28 | 64 | 68 | 57 | 14 | Xyloglucan endotransglucosylase_hydrolase 2 |
| Nitab4.5_0000243g0040 | 1 | 0 | 3 | 3 | 8 | 6 | 6 | 1 | -- |
| Nitab4.5_0000343g0160 | 27 | 31 | 17 | 37 | 76 | 132 | 91 | 52 | Mps one binder kinase activator-like 1A |
| Nitab4.5_0000111g0120 | 3 | 1 | 4 | 0 | 6 | 4 | 7 | 7 | Pentatricopeptide repeat-containing protein |
| Nitab4.5_0000022g0430 | 2 | 0 | 2 | 1 | 2 | 4 | 4 | 7 | Sodium_calcium exchanger protein (Fragment) |
| Nitab4.5_0000678g0040 | 1 | 0 | 0 | 1 | 1 | 2 | 2 | 1 | Serine_threonine kinase |
| Nitab4.5_0002240g0060 | 2 | 0 | 3 | 1 | 4 | 6 | 5 | 4 | Lysine ketoglutarate reductase trans-splicing related 1-like |
| Nitab4.5_0003338g0080 | 4 | 2 | 3 | 2 | 15 | 6 | 6 | 10 | Rhomboid family protein |
| Nitab4.5_0000227g0020 | 84 | 48 | 49 | 97 | 256 | 327 | 226 | 90 | Receptor like kinase, RLK |
| Nitab4.5_0002022g0020 | 2 | 0 | 2 | 3 | 3 | 8 | 4 | 11 | Pathogenesis-related protein |
| Nitab4.5_0003553g0130 | 2 | 0 | 2 | 1 | 2 | 4 | 6 | 5 | Homeobox-leucine zipper protein |
| Nitab4.5_0001885g0030 | 1 | 1 | 2 | 1 | 4 | 4 | 5 | 5 | DNA-directed RNA polymerase |
| Nitab4.5_0001580g0070 | 5 | 4 | 2 | 9 | 17 | 16 | 16 | 18 | Serine_threonine-protein kinase 38 |
| Nitab4.5_0000929g0060 | 3 | 1 | 1 | 2 | 4 | 10 | 5 | 3 | Receptor like kinase, RLK |
| Nitab4.5_0002823g0080 | 5 | 0 | 2 | 1 | 7 | 7 | 9 | 3 | Receptor-like kinase |
| Nitab4.5_0001003g0160 | 2 | 2 | 1 | 0 | 3 | 3 | 7 | 4 | At1g65470_F5I14_33 (Fragment) |
| Nitab4.5_0000166g0100 | 4 | 1 | 0 | 1 | 13 | 4 | 4 | 2 | Peptide transporter-like protein |
| Nitab4.5_0000358g0030 | 5 | 2 | 5 | 3 | 14 | 5 | 13 | 18 | 60S ribosomal protein L21-like protein |
| Nitab4.5_0003384g0010 | 3 | 6 | 3 | 0 | 8 | 8 | 14 | 9 | Unknown Protein |
| Nitab4.5_0000021g1000 | 1 | 1 | 1 | 0 | 2 | 3 | 1 | 6 | Inositol-tetrakisphosphate 1-kinase 1 |
| Nitab4.5_0000232g0060 | 2 | 0 | 3 | 1 | 5 | 5 | 4 | 7 | A_IG002N01.30 protein (Fragment) |
| Nitab4.5_0001911g0060 | 1 | 1 | 1 | 1 | 2 | 4 | 2 | 2 | Pentatricopeptide repeat-containing protein |
| Nitab4.5_0001039g0020 | 1 | 0 | 1 | 4 | 7 | 6 | 7 | 4 | Os06g0524700 protein (Fragment) |
| Nitab4.5_0000002g0330 | 1 | 1 | 1 | 1 | 6 | 3 | 0 | 3 | CAS1 domain containing 1 |
| Nitab4.5_0002016g0020 | 1 | 1 | 2 | 4 | 7 | 13 | 5 | 1 | Multidrug resistance protein mdtK |

| | | | | | | | | | |
|---|---|---|---|---|---|---|---|---|---|
| Nitab4.5_0000145g0040 | 0 | 1 | 1 | 1 | 4 | 3 | 1 | 1 | Cc-nbs-lrr, resistance protein with an R1 specific domain |
| Nitab4.5_0000232g0150 | 3 | 2 | 1 | 7 | 13 | 5 | 12 | 12 | -- |
| Nitab4.5_0003328g0010 | 6 | 3 | 2 | 1 | 11 | 19 | 8 | 5 | Unknown Protein |
| Nitab4.5_0001816g0070 | 1 | 1 | 2 | 2 | 7 | 7 | 1 | 5 | Uncharacterized conserved membrane protein |
| Nitab4.5_0000037g0180 | 212 | 123 | 80 | 236 | 594 | 794 | 549 | 326 | Xyloglucan endotransglucosylase_hydrolase 8 |
| Nitab4.5_0000563g0180 | 6 | 19 | 11 | 8 | 35 | 52 | 46 | 19 | Cyclin-dependent protein kinase regulator Pho80 |
| Nitab4.5_0000284g0090 | 1 | 2 | 3 | 1 | 10 | 4 | 3 | 2 | Speckle-type POZ protein |
| Nitab4.5_0000103g0070 | 0 | 0 | 1 | 0 | 1 | 2 | 2 | 1 | SET and MYND domain containing 3 |
| Nitab4.5_0000986g0060 | 2 | 2 | 0 | 1 | 9 | 1 | 2 | 3 | Uncharacterized membrane protein |
| Nitab4.5_0000262g0180 | 0 | 1 | 1 | 1 | 2 | 1 | 2 | 4 | Pectinesterase |
| Nitab4.5_0000092g0030 | 2 | 1 | 1 | 1 | 5 | 7 | 10 | 1 | 4-alpha-glucanotransferase |
| Nitab4.5_0000008g0360 | 0 | 1 | 1 | 2 | 0 | 5 | 5 | 8 | Zinc finger family protein (Fragment) |
| Nitab4.5_0000351g0070 | 0 | 1 | 1 | 1 | 2 | 4 | 3 | 1 | ATP-dependent DNA helicase Ta0057, Rad3 type |
| Nitab4.5_0000209g0140 | 1 | 2 | 2 | 2 | 8 | 4 | 12 | 3 | EPIDERMAL PATTERNING FACTOR-like protein 6 |
| Nitab4.5_0005893g0010 | 29 | 17 | 24 | 23 | 64 | 125 | 95 | 55 | 1-aminocyclopropane-1-carboxylate oxidase 1 |
| Nitab4.5_0000052g0280 | 2 | 0 | 2 | 6 | 10 | 12 | 10 | 8 | Uncharacterized basic helix-loop-helix protein At1g06150 |
| Nitab4.5_0001293g0050 | 2 | 1 | 0 | 3 | 4 | 5 | 15 | 1 | Cyclin-dependent protein kinase regulator Pho80 |
| Nitab4.5_0000845g0120 | 1 | 0 | 0 | 2 | 6 | 5 | 1 | 2 | Cysteine-rich repeat secretory protein 3 |
| Nitab4.5_0003098g0060 | 1 | 2 | 0 | 1 | 1 | 4 | 3 | 5 | Mitochondrial glycoprotein |
| Nitab4.5_0000009g0500 | 1 | 0 | 1 | 3 | 5 | 6 | 3 | 3 | Expansin-like protein |
| Nitab4.5_0002287g0070 | 1 | 1 | 2 | 1 | 6 | 2 | 5 | 3 | Tetraacyldisaccharide 4_apos-kinase family protein |
| Nitab4.5_0001840g0060 | 2 | 2 | 1 | 3 | 8 | 15 | 5 | 2 | Purple acid phosphatase |
| Nitab4.5_0003976g0070 | 0 | 1 | 1 | 1 | 2 | 3 | 4 | 6 | Cyclin-dependent protein kinase regulator-like protein |
| Nitab4.5_0002252g0060 | 2 | 1 | 1 | 1 | 8 | 9 | 3 | 1 | Pollen allergen Phl p 11 |
| Nitab4.5_0001935g0050 | 1 | 0 | 1 | 1 | 5 | 3 | 2 | 1 | Prolyl 3-hydroxylase 1 |
| Nitab4.5_0000263g0090 | 2 | 1 | 11 | 2 | 15 | 27 | 12 | 11 | DNA binding protein |

| | | | | | | | | | |
|---|---|---|---|---|---|---|---|---|---|
| Nitab4.5_0002904g0030 | 1 | 1 | 0 | 2 | 3 | 0 | 5 | 3 | -- |
| Nitab4.5_0000101g0200 | 7 | 13 | 8 | 4 | 46 | 44 | 29 | 9 | Cytokinin riboside 5_apos-monophosphate phosphoribohydrolase LOG |
| Nitab4.5_0000244g0350 | 7 | 7 | 7 | 14 | 34 | 59 | 40 | 5 | Xyloglucan endotransglucosylase_hydrolase 9 |
| Nitab4.5_0002954g0050 | 1 | 1 | 1 | 0 | 3 | 3 | 2 | 1 | RING finger protein 24 |
| Nitab4.5_0000768g0030 | 4 | 1 | 5 | 1 | 15 | 11 | 14 | 4 | BHLH transcription factor |
| Nitab4.5_0001034g0150 | 3 | 2 | 2 | 3 | 4 | 18 | 13 | 4 | Multidrug resistance protein mdtK |
| Nitab4.5_0000479g0030 | 1 | 1 | 0 | 2 | 4 | 3 | 5 | 5 | Snurportin-like protein |
| Nitab4.5_0000980g0290 | 78 | 24 | 20 | 35 | 396 | 74 | 107 | 74 | -- |
| Nitab4.5_0002195g0010 | 8 | 0 | 9 | 1 | 13 | 34 | 20 | 9 | DNA binding protein |
| Nitab4.5_0000307g0280 | 1 | 0 | 1 | 1 | 5 | 3 | 2 | 1 | Phosphatidylinositol-4-phosphate 5-kinase family protein |
| Nitab4.5_0000790g0120 | 0 | 0 | 1 | 0 | 1 | 1 | 2 | 2 | Glycosyltransferase family 77 protein |
| Nitab4.5_0000252g0180 | 0 | 0 | 1 | 0 | 2 | 1 | 1 | 2 | Unknown Protein |
| Nitab4.5_0001039g0080 | 0 | 0 | 1 | 0 | 1 | 3 | 3 | 1 | Microtubule plus-end binding protein |
| Nitab4.5_0002152g0060 | 1 | 2 | 2 | 3 | 9 | 12 | 11 | 4 | Zinc finger family protein (Fragment) |
| Nitab4.5_0000358g0040 | 1 | 0 | 1 | 1 | 1 | 6 | 5 | 4 | Unknown Protein |
| Nitab4.5_0000934g0180 | 0 | 0 | 1 | 0 | 3 | 2 | 2 | 1 | Cyclin A-like protein |
| Nitab4.5_0002972g0030 | 0 | 1 | 2 | 0 | 3 | 2 | 4 | 4 | Protein serine_threonine kinase |
| Nitab4.5_0000071g0140 | 0 | 1 | 1 | 1 | 2 | 2 | 6 | 5 | 1-(5-phosphoribosyl)-5-((5-phosphoribosylamino)methylideneamino)imidazole-4-carboxamide isomerase |
| Nitab4.5_0001797g0070 | 1 | 0 | 1 | 3 | 9 | 7 | 4 | 3 | Squamosa promoter binding protein 3 |
| Nitab4.5_0001553g0010 | 1 | 0 | 0 | 1 | 2 | 2 | 3 | 0 | Nbs-lrr, resistance protein |
| Nitab4.5_0000184g0070 | 0 | 1 | 0 | 0 | 2 | 1 | 1 | 2 | Essential meiotic endonuclease 1B |
| Nitab4.5_0001323g0050 | 0 | 0 | 2 | 1 | 8 | 4 | 3 | 0 | -- |
| Nitab4.5_0000127g0020 | 2 | 3 | 1 | 0 | 6 | 4 | 13 | 3 | Unknown Protein |
| Nitab4.5_0000481g0010 | 5 | 2 | 0 | 1 | 10 | 20 | 8 | 1 | Ethylene-responsive transcription factor 7 |
| Nitab4.5_0000776g0070 | 0 | 1 | 1 | 1 | 2 | 2 | 2 | 4 | LRR receptor-like serine_threonine-protein kinase, RLP |

| | | | | | | | | | |
|---|---|---|---|---|---|---|---|---|---|
| Nitab4.5_0002321g0020 | 1 | 1 | 0 | 1 | 2 | 1 | 3 | 6 | Unknown Protein |
| Nitab4.5_0001151g0060 | 1 | 2 | 1 | 0 | 3 | 6 | 7 | 2 | Receptor-like protein kinase At3g21340 |
| Nitab4.5_0002918g0050 | 0 | 1 | 0 | 0 | 3 | 1 | 2 | 1 | Protein phosphatase 2C |
| Nitab4.5_0000396g0030 | 1 | 2 | 4 | 0 | 11 | 18 | 6 | 4 | EPIDERMAL PATTERNING FACTOR-like protein 2 |
| Nitab4.5_0004479g0040 | 0 | 0 | 1 | 1 | 5 | 4 | 3 | 0 | Telomere repeat-binding protein 4 |
| Nitab4.5_0000034g0260 | 0 | 0 | 2 | 2 | 4 | 0 | 12 | 5 | Sentrin-specific protease 2 |
| Nitab4.5_0001402g0150 | 2 | 0 | 0 | 1 | 3 | 2 | 8 | 3 | Gpi-anchor transamidase |
| Nitab4.5_0005625g0020 | 6 | 5 | 5 | 3 | 31 | 49 | 22 | 6 | Purple acid phosphatase |
| Nitab4.5_0001639g0040 | 0 | 1 | 0 | 1 | 6 | 5 | 4 | 2 | F-box family protein |
| Nitab4.5_0000849g0010 | 0 | 1 | 0 | 0 | 1 | 2 | 2 | 5 | RAG1-activating protein 1 homolog |
| Nitab4.5_0000332g0130 | 107 | 65 | 60 | 46 | 433 | 678 | 400 | 87 | -- |
| Nitab4.5_0001921g0030 | 4 | 0 | 4 | 3 | 30 | 21 | 8 | 2 | Phi-1 protein (Fragment) |
| Nitab4.5_0002716g0020 | 3 | 0 | 1 | 0 | 15 | 6 | 3 | 4 | Unknown Protein |
| Nitab4.5_0000563g0300 | 0 | 0 | 0 | 2 | 2 | 2 | 2 | 8 | Blue copper protein |
| Nitab4.5_0000725g0060 | 0 | 0 | 0 | 1 | 5 | 1 | 0 | 2 | O-methyltransferase 1 |
| Nitab4.5_0000848g0010 | 0 | 0 | 2 | 0 | 8 | 5 | 5 | 0 | Glutathione S-transferase |
| Nitab4.5_0001531g0030 | 1 | 0 | 0 | 0 | 2 | 1 | 3 | 4 | L-lactate dehydrogenase |
| Nitab4.5_0000980g0200 | 0 | 0 | 0 | 0 | 2 | 0 | 2 | 1 | Tripeptidyl peptidase II |
| Nitab4.5_0000065g0070 | 0 | 1 | 1 | 2 | 9 | 8 | 0 | 11 | -- |
| Nitab4.5_0001529g0020 | 0 | 0 | 0 | 2 | 2 | 8 | 5 | 1 | Protease inhibitor_seed storage_lipid transfer protein family protein |
| Nitab4.5_0004977g0030 | 0 | 0 | 0 | 2 | 0 | 4 | 4 | 11 | Unknown Protein |
| Nitab4.5_0001329g0130 | 0 | 0 | 0 | 0 | 2 | 1 | 5 | 1 | DNA repair and recombination protein radA |
| Nitab4.5_0001365g0050 | 0 | 0 | 0 | 1 | 4 | 1 | 0 | 4 | Epidermal growth factor receptor substrate 15 |
| Nitab4.5_0003061g0020 | 0 | 0 | 0 | 0 | 3 | 2 | 0 | 2 | Myb-related transcription factor |
| Nitab4.5_0000401g0220 | 0 | 0 | 0 | 0 | 0 | 2 | 3 | 1 | HAPp48 5 protein (Fragment) |
| Nitab4.5_0001642g0050 | 0 | 0 | 1 | 2 | 6 | 9 | 7 | 2 | Gibberellin-regulated protein 2 |

| | | | | | | | | | |
|---|---|---|---|---|---|---|---|---|---|
| Nitab4.5_0001756g0030 | 0 | 0 | 0 | 0 | 4 | 2 | 0 | 3 | Protection of telomeres 1 protein |
| Nitab4.5_0000343g0220 | 0 | 0 | 0 | 0 | 4 | 3 | 0 | 1 | Unknown Protein |
| Nitab4.5_0003374g0070 | 0 | 0 | 0 | 0 | 0 | 0 | 3 | 2 | BHLH transcription factor |
| Nitab4.5_0000850g0170 | 0 | 0 | 1 | 0 | 4 | 0 | 6 | 3 | Actin-related protein 2_3 complex subunit 4 |
| Nitab4.5_0001239g0070 | 0 | 0 | 0 | 0 | 3 | 2 | 0 | 0 | Unknown Protein |
| Nitab4.5_0002438g0030 | 0 | 0 | 0 | 0 | 2 | 8 | 4 | 0 | High affinity sulfate transporter 2 |

**Table S4. Primers used in this study**

| Genes | Forward primer (5'-3') | Reverse primer (5'-3') |
|---|---|---|
| **Primers used in qPCR** | | |
| *BtActin* | TCTTCCAGCCATCCTTCTTG | CGGTGATTTCCTTCTGCATT |
| *BtRDP* | TTGGCTTTCCTTGTCCTCGC | CGTCGCAGAGTTCGTAGTCA |
| *Nttubulin* | AAGTACATGGCTTGCTGCCT | ATCAATGCGCGAGAAGACCT |
| *NtGAPDH* | GCAGTGAACGACCCATTTATCTC | AACCTTCTTGGCACCACCCT |
| *NtPAL* | AAGAAGCGTTCCGTGTTGCTG | TCGGGCTTTCCATTCATCACC |
| *NtNPR1* | GCTGTAGCGTTCCTTGTTGA | AGGCCTTATCAAGGGTTATG |
| *NtFAD7* | CATGTGGCTTGACTTAGTTACCTACT | CCCTGACTTCTTTGGCTCCTT |
| *NtPDF1.2* | GGAAATGGCAAACTCCATGCG | ATCCTTCGGTCAGACAAACG |
| *NtRLP4* | ATTCCTGAAAGCCTTGGGCA | AGCGTAAGATGGGTTCCACA |
| *NtSOBIR1* | GCGAAAGGCAAACAGAACACA | GGACCTCCAGTTTACGGCAT |
| *SlRLP4* | TGACCTCATTACGGACGCTG | CATGCCCCTAATCCGATCCC |
| *OsRLP4* | TCTTGACAGCGAGCCGATTT | TGGATCACCATGACCAGTGC |
| *NlSP104* | GTCTTAGCCGTGTGCATAGC | TGAAGAGGCGGATTGTTGGA |
| **Primers used in double stranded RNA synthesis** | | |
| *GFP* | TAATACGACTCACTATAGGGAGAATGAGT AAAGGAGAAGAACTTTTC | TAATACGACTCACTATAGGGAGATTTGTAT AGTTCATCCATGCCATGT |

| | | |
|---|---|---|
| *BtRDP* | TAATACGACTCACTATAGGGTTGGCCGTCCCAACCGCCACCC | TAATACGACTCACTATAGGGTCACAAGTAGATGCTCGGGTTG |
| *NlSP104* | TAATACGACTCACTATAGGGAACCCCAACCCGAAGCCGAT | TAATACGACTCACTATAGGGCTACAAGAAAGGCATGTATTGTC |
| **Primers used in binary vector construction** | | |
| *BtRDP-flag/mCherry* | ATGCACAAATTATTGGCTTTCCATGCACAAATTATTGGCTTTCC | GAGGAGAAGAGCCGTCGCAAGTAGATGCTCGGGTTG |
| *BtRDP*$^{-sp}$*- flag* | CGACGACAAGACCGTCACCATGTTGGCCGTCCCAACCGCCACCC | GAGGAGAAGAGCCGTCGCAAGTAGATGCTCGGGTTG |
| *NtRLP4-myc/gfp* | CGACGACAAGACCGTCACCATGATGAGATTCCACTATGGTTTCT | GAGGAGAAGAGCCGTCGGGTAAGCAAGGGAGGTC |
| *NtRLP4-mCherry* | CGACGACAAGACCGTCACCATGGATCCATATGTAATGCGAATAAGC | GAGGAGAAGAGCCGTCGGGTAAGCAAGGGAGGTC |
| *NtSOBIR1-gfp* | GACGAGCTGTACAAGGGTACCATGGCCTTCACTGCTTCACAAATCC | GCGGACTCTAGTTCATCTAGATTAATGCTTGATCTGAGTTAAC |
| *NtSOBIR1-flag* | CGACGACAAGACCGTCACCATGAAACTAAATCTCTATCCACC | GAGGAGAAGAGCCGTCGATGCTTGATCTGAGTTAAC |
| *RFP-mCheery* | CTTCGACGACAAGACCGGGCCCATGGCCTCCTCCGAGAACGTCA | AGTGAGGAGAAGAGCCGGGCCCACAGGAACAGGTGGTGGCGG |

| | | |
|---|---|---|
| *NLSP104-flag* | CGACGACAAGACCGTCACCATGCATCCAAGAACGATCATCCG | GAGGAGAAGAGCCGTCGTGCCGGACTACCCCCACCT |
| *BtFTSP-flag* | CGACGACAAGACCGTCACCATGTCAGCCCTAAGCTTTACTG | GAGGAGAAGAGCCGTCGCAAGAAAGGCATGTATTGTC |
| *SLRLP4-myc* | CGACGACAAGACCGTCACCATGCGCCATGAGCCATATGTAATG | GAGGAGAAGAGCCGTCGGGTAAGCAAGGGAGGTCC |
| *OsRLP4-myc* | CGACGACAAGACCGTCACCATGGCGGATCCTAGCAAAGAGCC | GAGGAGAAGAGCCGTCGGGAAGGAAGCAAATGTGG |
| **Primers used in Y2H vector construction** | | |
| *AD-NtRLP4$_{(23-541)}$* | GTACCAGATTACGCTCATATGGATCCATATGTAATGCGAATAAGC | CAGCTCGAGCTCGATGGATCCTTAACACGTTGGTAATCCCG |
| *AD-NtRLP4$_{(31-336)}$* | GTACCAGATTACGCTCATATGATAAGCTGTGGAGCTCGAC | CAGCTCGAGCTCGATGGATCCTTAAATCTCAAAAATTTC |
| *AD-NtRLP4$_{(375-541)}$* | GTACCAGATTACGCTCATATGCCTGACGAAGTCAAGGG | CAGCTCGAGCTCGATGGATCCTTAACACGTTGGTAATCCCG |
| *AD-NtRLP4$_{(573-625)}$* | GTACCAGATTACGCTCATATGGGAACCCATCTTACGCTTG | CAGCTCGAGCTCGATGGATCCTCAGGTAAGCAAGGGAGG |
| *AD-SLRLP4$_{(23-546)}$* | GTACCAGATTACGCTCATATGATAAGCTGTGGAGCTCGAC | CAGCTCGAGCTCGATGGATCCTTAATCATGTAAGATGTGTTCCGCAAG |

| | | |
|---|---|---|
| *AD-OsRLP4*$_{(29\text{-}551)}$ | GTACCAGATTACGCTCATATGGCGGATCCTAGCAAAGAGCC | CAGCTCGAGCTCGATGGATCCTGCCGCATTCATGTAAACCGG |
| *AD-OsRLP4*$_{(40\text{-}374)}$ | GTACCAGATTACGCTCATATGATAAGCTGTGGGAGTTTTG | CAGCTCGAGCTCGATGGATCCTCTCAAAGACCTCAATAGC |
| *AD-OsRLP4*$_{(377\text{-}551)}$ | GTACCAGATTACGCTCATATGGGCCGAAAAGAAAACTTTAAC | CAGCTCGAGCTCGATGGATCCTGCCGCATTCATGTAAACCGG |
| *AD-OsRLP4*$_{(552\text{-}582)}$ | GTACCAGATTACGCTCATATGCCGCATTTATCTGTGGCTGC | CAGCTCGAGCTCGATGGATCCTCTAGGAAGGAAGCAAATGTGG |
| *BK-BtRDP*$^{-sp}$ | TCAGAGGAGGACCTGCATATGTTGGCCGTCCCAACCGCCACCC | CCGCTGCAGGTCGACGGATCCTCACAAGTAGATGCTCGGGTTG |
| *BK-BtSP37.4*$^{-sp}$ | TCAGAGGAGGACCTGCATATGGACAGTGGAGCTACAGCGCC | CCGCTGCAGGTCGACGGATCCTCTAAAAGTGGTGGGTGGTTTGT |
| *BK-BtSP16.3*$^{-sp}$ | TCAGAGGAGGACCTGCATATGTTGCCGTCGAAGGAACCTGGA | CCGCTGCAGGTCGACGGATCCAGAGAGCACCATTTAAGATCCTC |
| *BK-BtFTSP*$^{-sp}$ | TCAGAGGAGGACCTGCATATGGGCAAAGATGAAGGCAAAGG | CCGCTGCAGGTCGACGGATCCTTATGCCGGACTACCCCCAC |
| *BK-NLSP104*$^{-sp}$ | TCAGAGGAGGACCTGCATATGAACCCCAACCCGAAGCCGAT | CCGCTGCAGGTCGACGGATCCCTACAAGAAAGGCATGTATTGTC |
| *BK-NlSP32030*$^{-sp}$ (XP_039291719.1) | TCAGAGGAGGACCTGCATATGAAATATGTAACCTCGGCAGTGC | CCGCTGCAGGTCGACGGATCCTTATTGCTTGGTGGCCCCTGCAG |

| | | |
|---|---|---|
| *BK-NlSP706^-sp^* (MF278706.1) | TCAGAGGAGGACCTGCATATGACTAGCGATGACGATTGCAGAG | CCGCTGCAGGTCGACGGATCCTTACAAAAGTCCCATATCTGCTAA |
| *BK-NlSP8^-sp^* (KU365967.1) | TCAGAGGAGGACCTGCATATGAGGTATCCTGGCTTTGGCGGG | CCGCTGCAGGTCGACGGATCCCTATGGGAATCCTGGGTATGG |
| *BK-NlSP711^-sp^* (MF278711.1) | TCAGAGGAGGACCTGCATATGGCAAGTACGGACATGGAATTC | CCGCTGCAGGTCGACGGATCCTCAGAAAGTAACATCCATTCCC |
| *BK-NlSP714^-sp^* (MF278714.1) | TCAGAGGAGGACCTGCATATGGAAACGTCTGAAGTCTACCTC | CCGCTGCAGGTCGACGGATCCTTATTCGTTTAGAATTCTGGCAAG |
| *BK-NlSP715^-sp^* (MF278715.1) | TCAGAGGAGGACCTGCATATGACGGGAAAATTTGATTTTGGAG | CCGCTGCAGGTCGACGGATCCTCAAGCACTGATTTTCGCTTC |
| *BK- NlSP720^-sp^* (MF278720.1) | TCAGAGGAGGACCTGCATATGGTACAGGAATCTAAATCATGGGC | CCGCTGCAGGTCGACGGATCCTCACTTAAAAAGCGGACTGTAG |
| *BK-NlSP28^-sp^* (XP_022195702.1) | TCAGAGGAGGACCTGCATATGACGAAGGGCGTAGAAGATGTG | CCGCTGCAGGTCGACGGATCCTAGTACTTGGGGATTGGAATC |
| *BK-NlSP474^-sp^* (XP_022196818.2) | TCAGAGGAGGACCTGCATATGTGTGTGGAGGGTACAGCAGC | CCGCTGCAGGTCGACGGATCCTCACGAGAATTTCGCATTCCC |
| *BK-NlSP170^-sp^* (XP_039284081.1) | TCAGAGGAGGACCTGCATATGTCTGGAACAGCAATGACTTCTG | CCGCTGCAGGTCGACGGATCCCCACCACCAATAATAAGGTACT |
| *BK-NlSP2^-sp^* (XP_022192221.2) | TCAGAGGAGGACCTGCATATGGCAGCTGTGGACCTTAGCTTC | CCGCTGCAGGTCGACGGATCCTCAACTGAAATCGACATCTCC |

| | | |
|---|---|---|
| *BK-NlSP3244*$^{-sp}$ (XP_022207944.2) | TCAGAGGAGGACCTGCATATGGCGAATATTGCTGACCATGGA | CCGCTGCAGGTCGACGGATCCCAATTGATAGTCAGTTTGACG |
| *BK-NlSHP*$^{-sp}$ (XP_022207944.2) | TCAGAGGAGGACCTGCATATGTTCCCATTTCCACTTCTCAGCC | CCGCTGCAGGTCGACGGATCCTTAGAAGGTCAATGACTGGAAGG |
| *BK-LsSP397*$^{-sp}$ (RZF44823.1) | TCAGAGGAGGACCTGCATATGGAATCAGATGATGTGCCGGTC | CCGCTGCAGGTCGACGGATCCCTATGCAGCTGGTGGTGTTGTG |
| *BK-LsSP19899*$^{-sp}$ (RZF42644.1) | TCAGAGGAGGACCTGCATATGGATTGCGAGACGAAACCGCATC | CCGCTGCAGGTCGACGGATCCTTATCCAGCATTTCCAACCAAG |
| *BK-LsSP3*$^{-sp}$ (RZF33006.1) | TCAGAGGAGGACCTGCATATGCGTTACGCCTCATATGAAAAGC | CCGCTGCAGGTCGACGGATCCCTAACATTTTCCACATTTTCCC |
| *BK-LsSP4*$^{-sp}$ (RZF48570.1) | TCAGAGGAGGACCTGCATATGGTGACATGTTTCCCGTTCCCC | CCGCTGCAGGTCGACGGATCCTTAGAAGGTCAATGACATGAC |
| *BK-LsSP5*$^{-sp}$ (RZF42817.1) | TCAGAGGAGGACCTGCATATGGCGGCTGTGCTACCAGCTAAC | CCGCTGCAGGTCGACGGATCCTCAAGGTATGGGTGGAGTTAGG |
| *BK-LsSP6*$^{-sp}$ (RZF33751.1) | TCAGAGGAGGACCTGCATATGGGGCCCAAATCCGACCGGAAAC | CCGCTGCAGGTCGACGGATCCTTAGTAGACAAGTTGTGGTTGC |
| **Primers used in prokaryotic expression vector construction** | | |
| *BtRDP*$^{-sp}$*-his* | CTGGTGCCGCGCGGCAGCCATATGTTGGCCGTCCCAACCGCCACCC | ACGGAGCTCGAATTCGAATCCTCACAAGTAGATGCTCGGGTTG |

| | | |
|---|---|---|
| *GFP-his* | CAGCAAATGGGTCGCGGATCCATGGTGAG CAAGGGCGAGGAG | GTGGTGGTGGTGGTGCTCGAGTGTACAGC TCGTCCATGCCGA |
| **Primers used in BIFC vector construction** | | |
| *nYFP-NtRLP4* | ATCGAGGACTCCGGAGTCGACATGGATCC ATATGTAATGCGAATAAGC | GATCGGGGAAATTCGAGCTCTCAGGTAAG CAAGGGAGGTC |
| *nYFP-NtCf-9* | CTGTACAAGTCCGGAGTCGACATGGATTA TGAAAATCTTGCA | GATCGGGGAAATTCGAGCTCTCTAATATCT TTTCTTGTGTTTTTTC |
| *cYFP-BtRDP$^{-sp}$* | CTGTACAAGTCCGGAGTCGACATGTTGGC CGTCCCAACCGCCACCC | GATCGGGGAAATTCGAGCTCTCACAAGTA GATGCTCGGGTTG |
| *cYFP-BtFTSP$^{-sp}$* | CTGTACAAGTCCGGAGTCGACATGGGCAA AGATGAAGGCAAAG | GATCGGGGAAATTCGAGCTCTTATGCCGG ACTACCCCCAC |
